# A reference tissue atlas for the human kidney

**DOI:** 10.1101/2020.07.23.216507

**Authors:** Jens Hansen, Rachel Sealfon, Rajasree Menon, Michael T. Eadon, Blue B. Lake, Becky Steck, Dejan Dobi, Samir Parikh, Tara K. Sigdel, Guanshi Zhang, Dusan Velickovic, Daria Barwinska, Theodore Alexandrov, Priyanka Rashmi, Edgar A. Otto, Michael P. Rose, Christopher R. Anderton, John P. Shapiro, Annapurna Pamreddy, Seth Winfree, Yongqun He, Ian H. de Boer, Jeffrey B. Hodgin, Laura Barisoni, Abhijit S. Naik, Kumar Sharma, Minnie M. Sarwal, Kun Zhang, Jonathan Himmelfarb, Brad Rovin, Tarek M. El-Achkar, Zoltan Laszik, John Cijiang He, Pierre C. Dagher, M. Todd Valerius, Sanjay Jain, Lisa Satlin, Olga G. Troyanskaya, Matthias Kretzler, Ravi Iyengar, Evren U. Azeloglu, for the Kidney Precision Medicine Project

## Abstract

Kidney Precision Medicine Project (KPMP) is building a spatially-specified human tissue atlas at the single-cell resolution with molecular details of the kidney in health and disease. Here, we describe the construction of an integrated reference tissue map of cells, pathways and genes using unaffected regions of nephrectomy tissues and undiseased human biopsies from 55 subjects. We use single-cell and -nucleus transcriptomics, subsegmental laser microdissection bulk transcriptomics and proteomics, near-single-cell proteomics, 3-D nondestructive and CODEX imaging, and spatial metabolomics data to hierarchically identify genes, pathways and cells. Integrated data from these different technologies coherently describe cell types/subtypes within different nephron segments and interstitium. These spatial profiles identify cell-level functional organization of the kidney tissue as indicative of their physiological functions and map different cell subtypes to genes, proteins, metabolites and pathways. Comparison of transcellular sodium reabsorption along the nephron to levels of mRNAs encoding the different sodium transporter genes indicate that mRNA levels are largely congruent with physiological activity.This reference atlas provides an initial framework for molecular classification of kidney disease when multiple molecular mechanisms underlie convergent clinical phenotypes.

## Introduction

The kidney has one of the most diverse cellular populations in the human body, and it is critical in maintaining the physiological homeostasis by regulating fluid and electrolyte balance, osmolarity and pH. The basic unit of organization in the kidney is the nephron embedded in the interstitium; the human kidney has between 210,000 to 2.7 million nephrons (*1*). There are multiple cell types in the nephron and the interstitium including those that comprise the blood vessels and capillaries (such as endothelial cells and vascular smooth muscle cells) and many types of immune cells. From the development of a structure based standard nomenclature (*2*), to a recent review (*3*), there has been a sustained effort to develop a detailed understanding of structure-function relations within the kidney tissue to understand its physiology and pathophysiology.

Over the past decade, with the advent of single-cell (sc) RNAseq technologies, substantial advances have been made in enumerating the different cell types in the human and mouse kidney (*4–17*). Computational analyses and modeling of single-cell transcriptomic data, and other types of omics data are starting to provide rich and deep insight into different kidney disease processes including kidney cancers (*17*) and fibrosis (*9*). These studies demonstrate the power of omics technologies in developing atlases that map structure-function relationships at the single-cell level within tissues.

Data sets from different omics technologies provide an unparalleled opportunity to understand how the diversity of cell types and their constituents underlie physiological functions and how they are altered in different disease states. The Kidney Precision Medicine Project (KPMP) is a consortium funded by the National Institute of Diabetes and Digestive and Kidney Diseases (NIDDK). Using kidney biopsies that are ethically and safely obtained from participants with kidney disease, KPMP aims at the creation of a kidney atlas in health and disease. Such an atlas can allow the identification of critical cells, pathways, and targets for novel therapies and preventive strategies (*18, 19*). To identify and understand disease states, it is necessary to have a detailed atlas of tissues that do not show disease phenotype by standard clinical histological evaluation. We call such an atlas a reference atlas. Using multiple kidney reference sources, different groups in the consortium have generated diverse types of data. Among these are single-nucleus (*20*) and single-cell (*21*) transcriptomics, regional bulk transcriptomics, proteomics and metabolomics as well as multiple complementary types of imaging methods. We have analyzed and integrated these different data types obtained from reference kidney tissue specimens, as evaluated by standard pathology analysis, from 55 human subjects. We have constructed maps of the different cell types in the kidney and the molecular entities as well as functional pathways within these cell types to develop an early version of a reference human kidney atlas. To determine if the molecular details in the atlas enables new insight into physiological activity we compare the transcellular sodium reabsorption along the nephron that is important for the maintenance of normal blood pressure in individuals with hypertension (*22, 23*). We find substantial congruence between physiological activity and the sum of the mRNA levels of different sodium transporters indicating that these such a molecular atlas can provide deep insight into molecular and cellular basis of physiological processes. This atlas is now available to serve as a starting point from which datasets emerging from disease states can be used to project into the integrated functional context and to drive new molecular classification of kidney diseases.

## Results

The KPMP Consortium conducted different types of omics as well as low throughput immunohistochemistry experiments at different sites for these reference atlas studies. Although it is impossible to definitely characterize tissue as healthy, clinical pathologists adjudicated that specimens used in this study show no signs of disease manifestations. Nevertheless, since we use unaffected tissue regions from nephrectomies as well as biopsies from both living donors and transplant recipients (i.e., surveillance biopsies), we use the general term reference tissue (Suppl. Table 1). In future studies, these can be compared to diseased tissue specimens.

There were four transcriptomic, two proteomic, two imaging-based, and one spatial metabolomics tissue interrogation assays deployed on the shared tissue samples. These assays yielded 3 to 48 different datasets obtained from 3 to 22 subjects per assay for a total of 55 different human subjects (Suppl. Table 1). The assays and their detailed tissue pre-analytical, tissue processing, data acquisition and analytical data processing pipelines are schematically depicted as a flowchart in Figure 1. We also summarize, in the integration segment of our flowchart, the steps by which the data sets from the different assays were integrated and harmonized. This is shown in the upper right side of this descriptive map.

**Figure 1.**
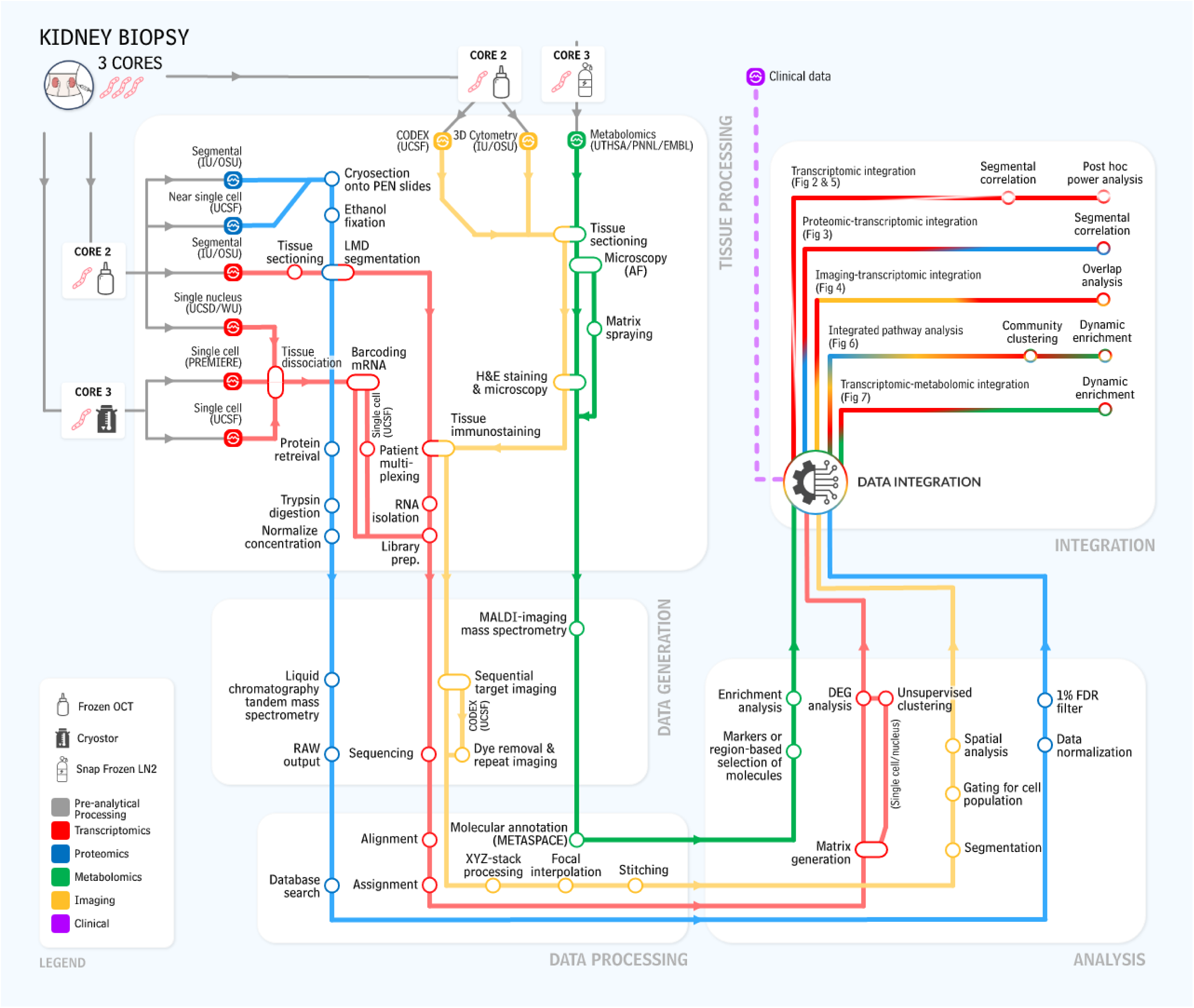
Graphic outline of KPMP data integration and harmonization procedures. The “subway map” representation of the experimental and analytical protocols used within KPMP is shown in operational flow from kidney biopsy to the integrated multimodal data represented in this manuscript. The kidney biopsy, which is processed through three different tissue processing methods, is shared among TISes that generate the data. Four key modalities of molecular data are generated: transcriptomic (red), proteomic (blue), imaging (yellow) and metabolomic (green). Biopsy core 2 and 3 are used for the molecular analysis, biopsy core 1 (not depicted) is used for histological analysis.

### Integration of multiple transcriptomic interrogation techniques shows agreement and technological synergy between assays

Separate as well as integrated analysis of single-cell (sc), multiplexed single-cell and single-nucleus (sn) transcriptomic datasets confirmed all known major kidney tissue cell types of the nephron (*20, 21*) and multiple immune cells (Figure 2A). Clustering algorithms used to separately analyze the sc and sn RNAseq data identified multiple subtypes for several cells. We observed differences between the numbers of subtypes in the sc versus sn data as different cutoffs were used in the initial analyses (*20, 21*). Nevertheless, when sc and sn RNAseq data were analyzed in an integrated manner, all major cell types were identified, as shown in the central panel in Figure 2A. Here, combined processing of 17,529 and 13,130 cells along with 17,657 nuclei yielded 16 main clusters (note that some clusters contain multiple closely related subtypes). These clusters were annotated to 14 cell types based on cluster specific marker gene expression. Each cluster contained cells and nuclei from every dataset, documenting consistency of our transcriptomic datasets (Suppl. Figure 1). To provide spatial context with respect to different regions of the nephron, we compared the sc and sn transcriptomic datasets with nephron segment specific bulk transcriptomic datasets that were obtained after Laser Microdissection (LMD) of kidney segments (*24*) (Suppl. Table 2). Cross-assay Pearson correlation analysis allowed us to map each single cell and nucleus to the nearest LMD segment (Figure 2B). We find that there is strong concordance across the data obtained by the different technologies, whereby the majority of the cells and nuclei from each cluster were assigned to the correct corresponding LMD subsegment in an unbiased manner. For example, proximal tubule (PT) cells were assigned to the PT subsegment, while podocytes were assigned to the glomerular subsegment.

**Figure 2.**
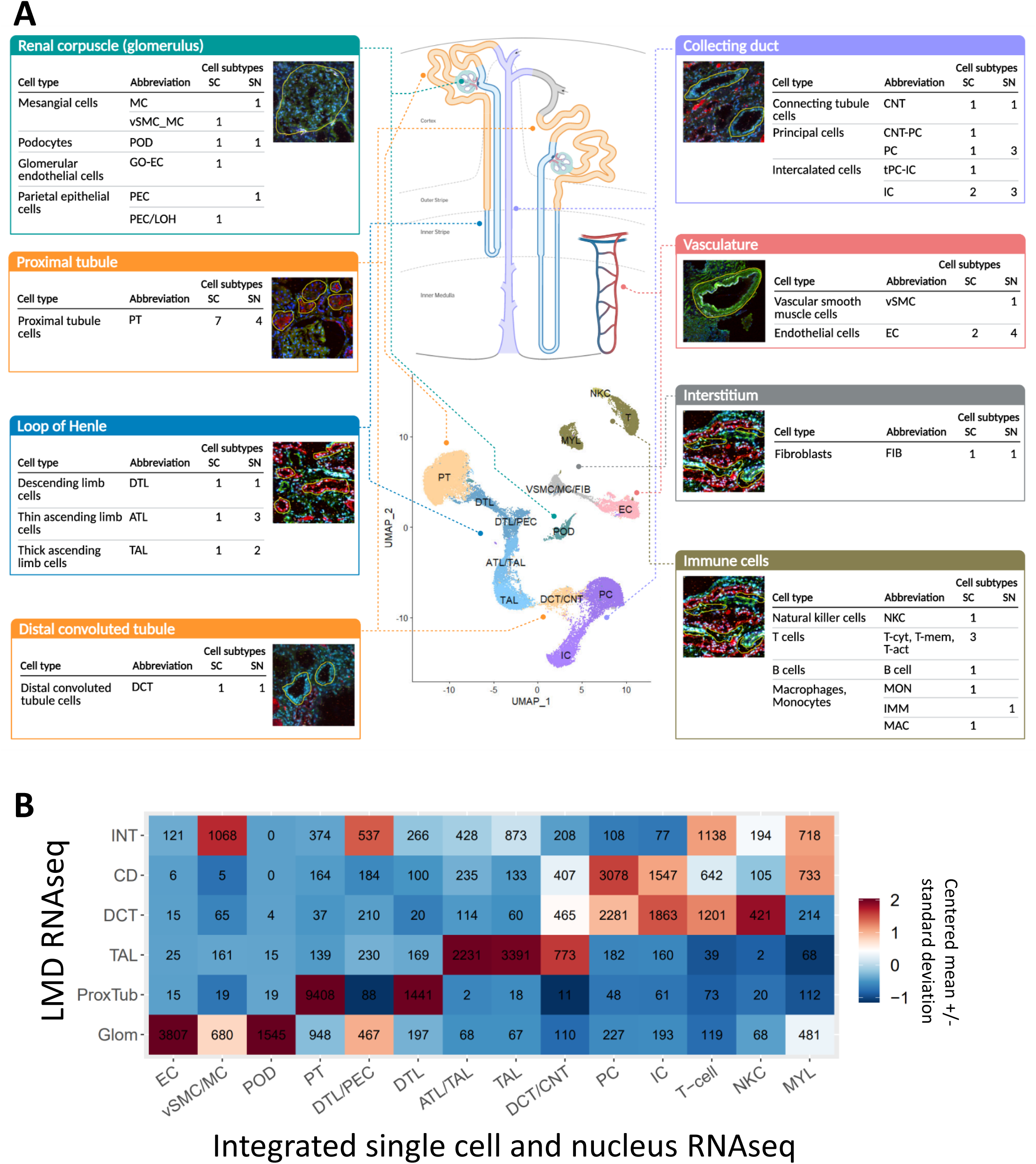
Integrated transcriptomic analysis reveals coherent cell-type-specific signatures. **(A)** Scheme showing the major nephron segments as identified in our datasets. Sc and sn datasets were either analyzed separately (*20, 21*) or combined. UMAP documents the results of the combined analysis. Cell subtype counts were obtained from the separated analyses (Suppl. Figure 4A/B). The corresponding LMD segments shown include the markers used to identify each subsegment: Phalloidin – FITC labeled phalloidin for dissection of glomeruli and other structures; LRP2 – Megalin with AlexaFluor 568 secondary (red); UMOD – directly conjugated AlexaFluor 546 Ab to uromodulin (red); fluorescein labeled PNA – Peanut Agglutinin labels collecting ducts (green); DAPI included for nuclei (blue). **(B)** Each cell or nucleus in the combined transcriptomic sc/sn analysis is mapped to the closest subsegment (subsegment with highest Pearson correlation of gene expression) in the LMD RNAseq data. To compute the Pearson correlation between the gene expression profiles of cells and LMD segments, the gene profiles were restricted to genes shared between the two datasets and showing variable expression in the single-cell dataset. Correlations were computed between the logarithm of the mean ratio vector for each LMD segment and the scaled expression profile of each cell in the sc/sn dataset. For each sc/sn cluster and LMD subsegment, the number of cells/nuclei from that cluster assigned to the corresponding segment is displayed in the heatmap. The heatmap is colored according to the number of cells/nuclei assigned to each LMD subsegment, scaled so each column has mean of 0 and standard deviation of 1. For the overlap between cell type annotations in the combined and integrated analyses see Supplementary Figure 4C/D and for the LMD mappings based on the separated analyses see Supplementary Figure 4E/F.

The total numbers of cells analyzed are small by current standards and hence we determined, if other independent orthogonal technologies support our overall atlas framework. Hence, we used integration of different omic technologies as well as posthoc power analyses to determine the validity of the atlas.

### Proteomic and transcriptomic assays produce biologically complementary descriptions of subsegmental molecular composition

In addition to transcriptomic profiles, we obtained subsegment specific protein expression profiles using two different proteomic assays. These assays identify protein expression in the glomerulus and the tubulointerstitium (LMD proteomics) or proximal tubule (Near Single Cell, NSC, proteomics) (Suppl. Tables 3 and 4, respectively). We then compared the proteomic data sets with the transcriptomic data sets. For an unbiased cross-platform comparison, we focused on podocyte/glomerular and proximal tubule (PT) cells and subsegments in the four transcriptomic datasets. To reduce assay related biases, we calculated, for each subject within each assay, the logarithmic ratios of gene or protein expression values for the glomerular versus tubular cell types or subsegments (Suppl. Figure 2A). Pairwise correlation of these logarithmic ratios, followed by hierarchical clustering, resulted in grouping of the data sets by appropriate regions of the kidney (Figure 3A). Within this broad classification, the subgroupings by different assays could be readily identified and are shown (Right side labels in Figure 3A). From this clustering, we conclude that irrespective of the assay, we can readily identify groups of genes or proteins associated with the appropriate anatomical region (i.e. glomerulus versus tubulointerstitium). This pattern is observed with or without removal of genes or proteins that are not identified by all technologies (Suppl. Figure 2B). In contrast, if we cluster by absolute expression values, the clustering is primarily driven by the assay used rather than the anatomical region. This is irrespective of whether we use datasets with and without removal of genes or proteins not detected by all technologies (Suppl. Figure 2C and 2D, respectively). These results suggest that rather than absolute presence or absence of the different genes or proteins, the relative expression levels are more indicative of the corresponding anatomical region of the kidney. It documents the high quality of our data, since technological bias can be overcome by a relatively simple algorithm. Correlation analysis of averaged log_2_ fold changes between all combined RNAseq datasets and combined proteomic datasets further supports the conclusions that similar entities are identified by different assays (Figure 3B). The 0.6 correlation value that we obtain is in agreement with the canonical value across mammalian tissues as described (*25*), though our comparison is based on fold changes and not absolute mRNA and protein abundancies. As such, integration of multiple datasets increases accuracy of the results, since integrated RNAseq and proteomic datasets show a higher correlation with each other than any individual RNAseq and proteomic datasets. Nevertheless, correlations between the same technologies when the assay was conducted at different sites is quite high (Figure 3C).

**Figure 3.**
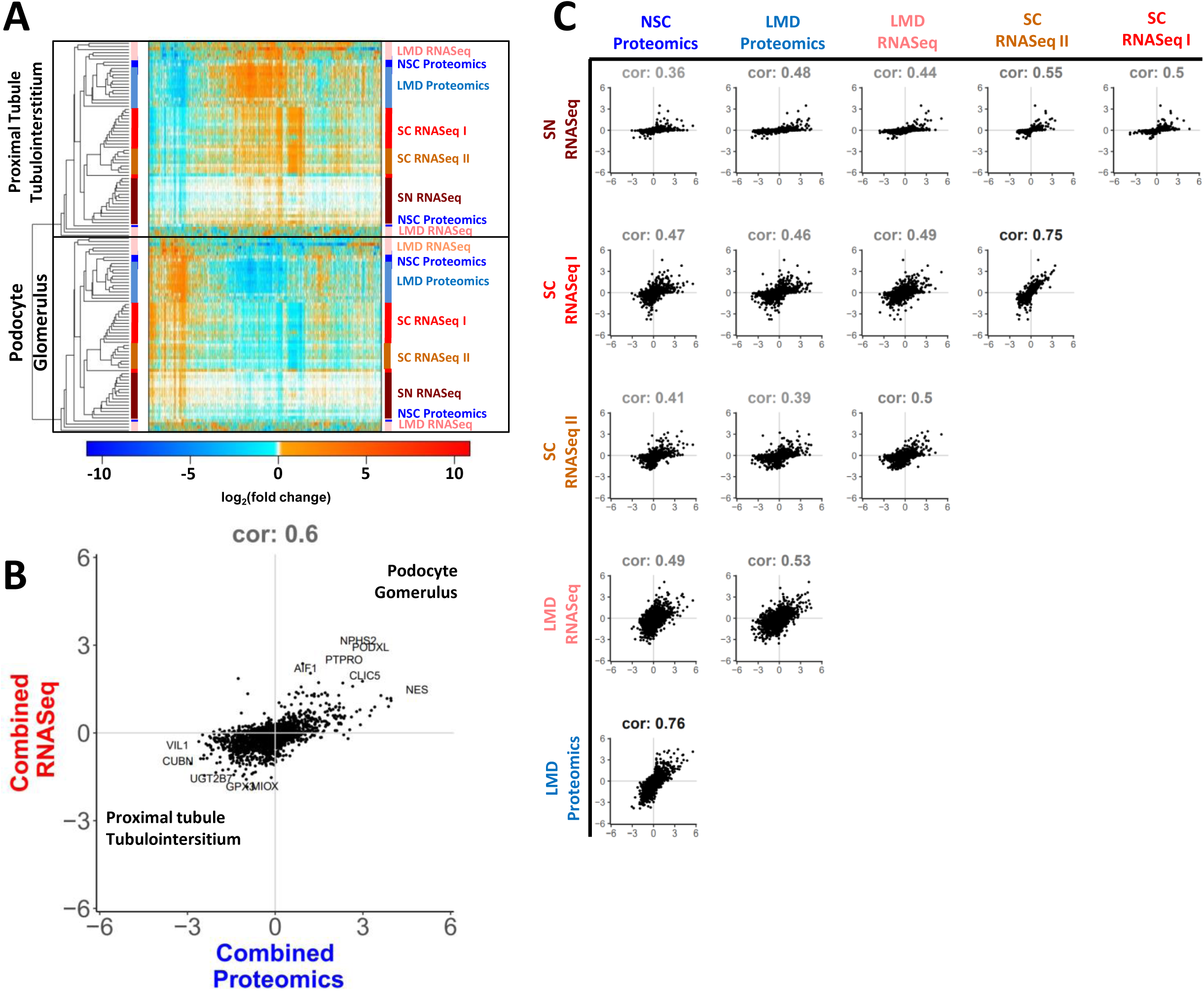
Correlation analyses demonstrate concordance across different omics technologies. Log_2_(fold changes) between podocyte (or glomerulus) and proximal tubule cells (or tubulointerstitium) were calculated for each subject based on each assay. Common genes/proteins identified by each assay subjected to comparative analysis. **(A)** Hierarchical clustering of pairwise correlation coefficients between the log_2_(fold changes) groups samples based on cell type/segment. Heatmap shows up- and downregulated genes/proteins of each sample in red and blue, respectively. Genes and proteins were rearranged according to the clustering results. White spots indicate undetected genes or no expression differences. Genes and proteins that are not consistently detected across all six technologies were removed. Nevertheless, observed grouping of samples by anatomical region is independent of this removal (Suppl. Figure 2B). **(B)** Log_2_(fold changes) obtained by the same assay were averaged across all subjects, followed by averaging of the results across all four transcriptomics and two proteomics assays. Positive (negative) log_2_(fold changes) indicate podocyte/glomerular (PT/tubulointerstitial) expression. In arbitrarily selected cases we replaced the dots by the official NCBI gene symbols. **(C)** Pairwise correlations between the sc/sn RNAseq and proteomic datasets document highest concordance between both proteomic and single-cell assays. Positive (negative) log_2_(fold changes) indicate podocyte/glomerular (PT/tubulointerstitial) expression.

### Imaging-based molecular data and non-spatial proteomic and transcriptomic assays together produce spatial marker expression signatures

Imaging assays can provide spatial specification of omics data, such as bulk proteomics (*26*) and confirm contextual framework for cell types inferred through sc transcriptomics (*27*). Those with well-characterized markers can identify the spatial localization of individual cells, which can be independently identified from gene expression patterns. By analyzing the relationship between cells identified from sc/sn sequencing technologies and CODEX imaging of canonical markers, we establish the concordance between the assay types for independently identifying cell types and inferring molecular profiles for spatially localized cells. We constructed a mapping matrix to transform the cell-type specific protein (i.e., marker) expression profiles measured using CODEX to cell type-specific gene expression profiles measured using sc and sn transcriptomic assays (Suppl. Figure 3). An entry in the mapping matrix is high if the corresponding imaging cell type is highly weighted in the linear combination of imaging cell type expression profiles that approximate the expression profile of a cell type in the single-cell transcriptomic dataset. We find that this mapping approach performs well for cell types with well-characterized cell type-specific canonical markers in the imaging dataset, such as endothelial cells and podocytes (Figure 4).

**Figure 4.**
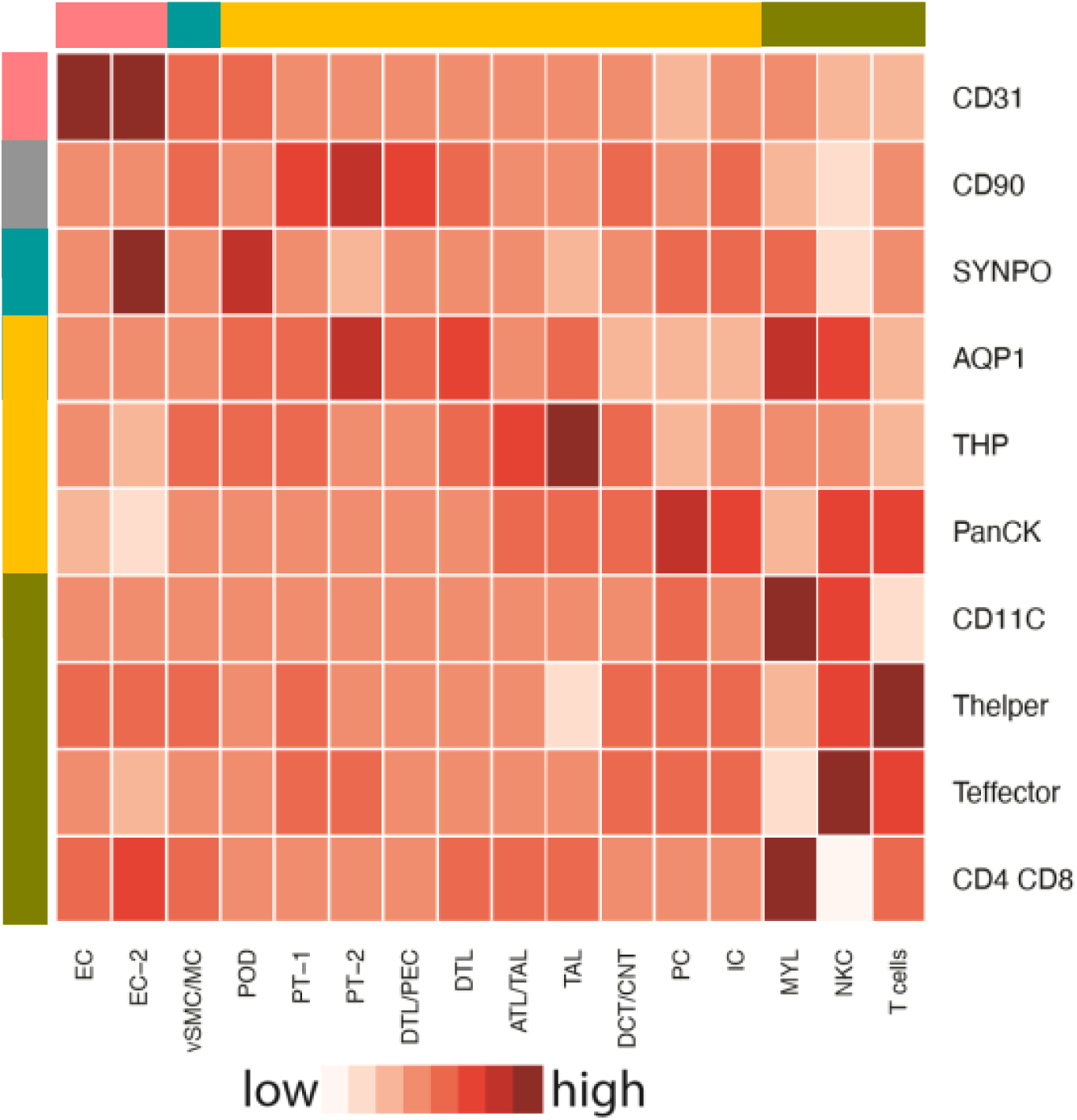
Imaging-based and transcriptomic assays show consistent cell-type-specific marker signatures. Mapping matrix showing relationship between markers characterizing CODEX cell-type clusters and transcriptomic cell-type clusters. Colorbars to the top and left of the heatmap show broad segmental/cell-type categories (red = endothelial, gray = fibroblast/mesangial, turquoise = podocyte, orange = tubular, gold = immune). See figure 2A for cell type abbreviations.

Both the mapping of single cell and nucleus expression profiles to LMD segments and the mapping of single cell and nucleus expression profiles to the imaging assays allow assignment of single cells and nuclei to anatomical regions within the kidney. Using these comparisons we can arrange the cells along the nephron and in the interstitium, allowing documentation of the order by which they encounter the glomerular ultrafiltrate.

### Integrated pathway enrichment analysis enables identification of functional capabilities of different cell types of the kidney

After establishing the consistency between transcriptomic, proteomic and imaging datasets, we used these integrated data to identify the cell-type specific functional pathways and network modules. Pathways and modules give rise to subcellular processes that together produce whole cell-level biochemical and physiological function. This pathway based approach that connects genes to cell level physiological function will serve as the basis for molecular classification of disease states. We started by using individual analyses of the sc and sn RNAseq datasets and identifying the pathways inferred from the expressed genes (*20, 21*) (Suppl. Figure 4A and B, respectively). In contrast to our integrated analysis of these datasets described above, the individual analyses used more relaxed quality control cutoffs such as allowing up to 50% mitochondrial gene expression so the cell subtype and type specific gene expression obtained by the single cell RNAseq dataset was based on 22,264 cells instead of 17,529 cells. These single cell technology analyses also allowed us to ascertain that all of the cell types could be observed independently of the method by which the reference tissue was obtained. We find that all major kidney cell types can be identified in nephrectomy, living donor biopsy and transplant surveillance biopsy tissues (Suppl. Figures 4A and 4B). An exception to this finding is that immune cells were mostly identified only within the sc RNAseq dataset, while only one cluster of the sn RNAseq dataset that contained less than 1% of all nuclei was annotated to an immune cell type, i.e. immature macrophage (Suppl. Figure 4B).

Individual analyses of sc and sn transcriptomic data ensure that these two related technologies do not computationally influence the ranking of combined pathways in ways that are not fully identifiable. Most cells identified from sc or sn RNAseq data sets in the individual analyses were annotated to the same cell types as in the combined analysis (Suppl. Figure 4C and 4D, respectively) and mapped to the appropriate LMD segment as well (Suppl. Figure 4E and 4F, respectively). A less stringent cutoff for mitochondrial gene expression (50% instead of 20%) allowed consideration of additional cells that were excluded from the combined analysis.

### *Post hoc* power analysis documents consistent cell-type detection

Before focusing on cell-type specific functions that we predict from pathway enrichment analysis and module mappings, we evaluated how many reference subject samples need to be processed to obtain consistently reproducible results. 24 and 47 libraries obtained from 22 and 15 subjects were subjected to sc (*21*) and sn (*20*) RNAseq, yielding 22,264 cells and 12,100 nuclei after quality control (Suppl. Table 1), respectively. We separately subjected both RNAseq datasets, with and without random and progressive removal of libraries, to a standardized sc and sn RNAseq analysis pipeline (Suppl. Figure 5A). Results obtained for the down sampled datasets were compared to those obtained for the complete datasets (Suppl. Figure 5B). Our results indicate that for a consistent detection of podocytes and mesangial cells (i.e. in at least 95% of all down sampled datasets with the same library counts), at least 9 (∼8,250 cells) or 7 libraries (1,837 nuclei) are needed if subjected to sc RNAseq (Figure 5A) or sn RNAseq (Figure 5B), respectively. The observed higher identification rate by the sn RNAseq assay is in agreement with a previous report that compared sn and sc RNAseq results obtained from mouse kidneys (*16*). Proximal tubule cells, thick ascending limb cells, principal cells, intercalated cells, T-cells were always detected in the downsampled sc RNAseq datasets. Macrophages were consistently detected, if 3 libraries (2,843 cells) were analyzed. In the sn RNAseq datasets we consistently detected proximal tubule cells, thick ascending limb cells, principal cells and intercalated cells in 4, 7, 9 and 6 libraries (1,013; 1,832; 2,323 and 1,527 nuclei), respectively. For additional cell types, see Figures 5A and 5B. Additionally, our results suggest that the accuracy of sc or sn assignments to the selected cell types is relatively stable as documented by the low number of cells that are assigned as different cell types or mapped to an unrelated tissue subsegment in the downsampled sc and sn datasets (Suppl. Figures 5C and 5D, respectively). Similarly, pearson correlation between cell type specific DEGs in the down sampled and full datasets follow the same trend. These analyses establish the rigor with which we are able to assign pathways and physiological functions to the different cell types.

**Figure 5:**
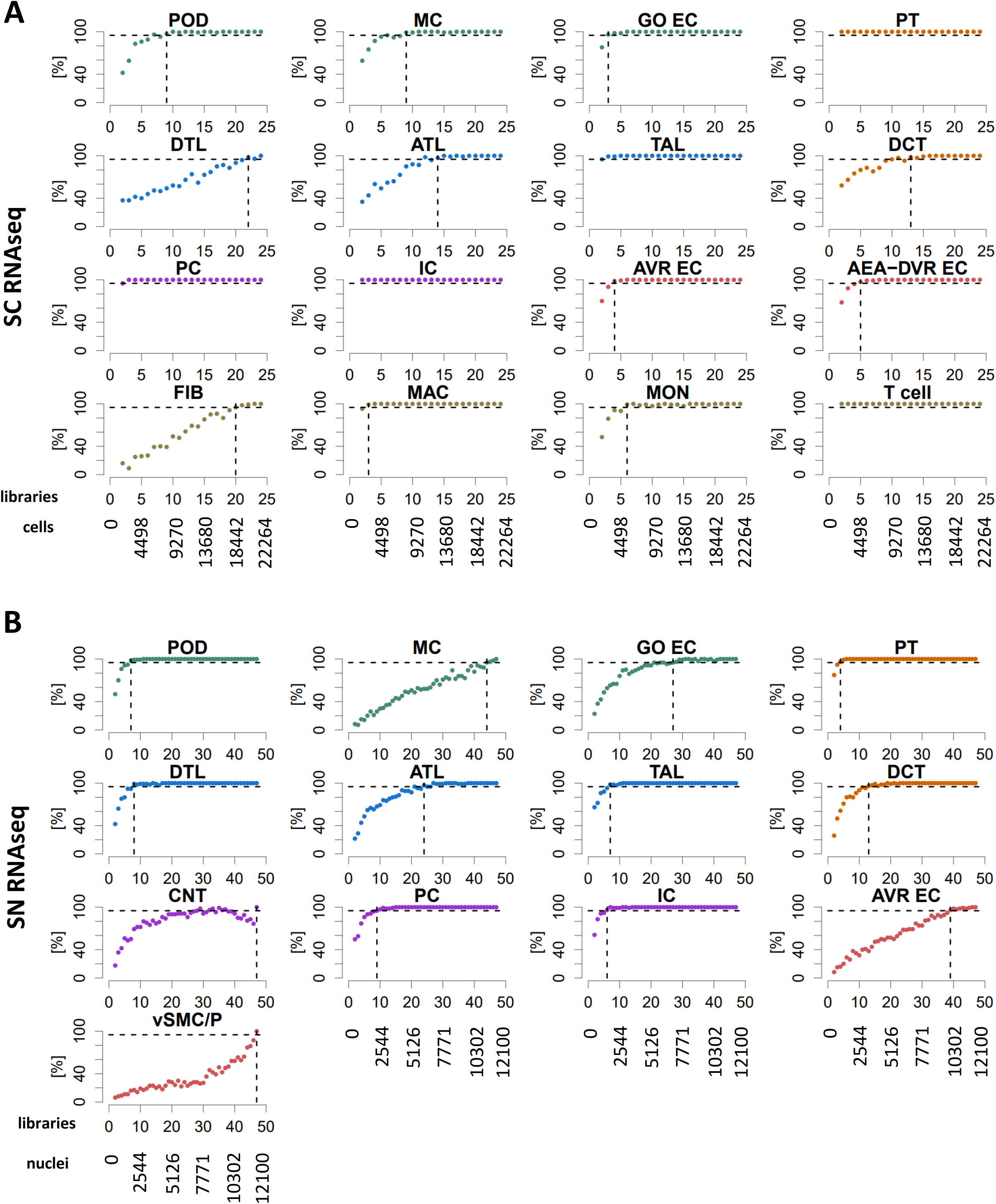
Single-cell/nucleus transcriptomic *post hoc* power analyses show that nine libraries are sufficient to identify most major kidney cell types. Subject libraries (or samples) were randomly and progressively removed from **(A)** the sc (24 libraries) and **(B)** sn (47 libraries) RNAseq to generate at max 100 non-overlapping random groups for the remaining samples. Sc and sn datasets were subjected to an automated data analysis pipeline (Suppl. Figure 5A). To assign cell types to the identified clusters we compared cluster specific markers of each analysis with literature curated cell type specific genes (Suppl. Figure 5B). We counted how many analyses based on the same number of remaining libraries that have identified a particular cell type. Horizontal dashed lines mark the 95% plateau; vertical dashed lines indicate the minimum number of libraries needed to identify a given cell type with a probability of 95%. See Suppl. Figure 5 for complete post hoc power analysis results. See figure 2A for cell type abbreviations.

### Pathway enrichment analysis and module identification

The top 300 significant gene and protein markers of each cell type or subtype and subsegment (Suppl. Table 5) were subjected to dynamic enrichment analysis using the Molecular Biology of the Cell Ontology (MBCO) (*28*) (Suppl. Table 7). In many cases, less than 300 markers were significant (Suppl. Table 6) and we consequently used only those for our downstream analysis. Dynamic enrichment analysis is a novel enrichment algorithm that considers dependencies between functionally related subcellular processes (SCPs), thereby addressing a limitation of standard enrichment analysis (*29*). In contrast to standard enrichment analysis that determines if a set of experimentally observedgenes enriches for genes annotated to a single SCP, dynamic enrichment analysis determines if gene set enriches for genes annotated to multiple functionally related SCPs. We comparatively assigned mRNAs (cognate proteins) to functionally related pathways enabling the formation of subnetworks that underlie subcellular processes (SCPs) that give rise to whole-cell physiological function (*28*). Functional relationships are defined in the MBCO network of subcellular processes that are predicted based on prior knowledge from primary literature of functional activities. Cell type and corresponding segment specific networks were merged. Non-glomerular and glomerular metabolites (Suppl. Table 8) were subjected to pathway enrichment analysis using MetaboAnalyst (*30*) (Suppl. Figure 6A and 6B, respectively). For the top eight predicted pathways, we manually determined if pathway specific metabolites in the metabolomics data sets could be identified. When at least one metabolite could be selectively associated with the predicted pathway, we added the pathway to the SCP network identified from the transcriptomic and proteomics datasets. We added two pathways that do not exist in MetaboAnalyst after curation of identified metabolites (Suppl. Figure 6A). This pathway-based integration process allowed us create maps of biochemical and physiological functions of all major cell types in the kidney, setting up the framework for the development of molecular classification of kidney diseases.

Significant mRNA and protein markers were used for community clustering in a kidney-specific functional network using HumanBase (*31, 32*) (Suppl. Table 9). In this network-based module detection analysis, genes are partitioned based on their connectivity in tissue-specific functional networks using a community clustering approach. These tissue-specific functional networks are constructed by integrating thousands of public genomic datasets using a regularized Bayesian framework to predict the probability that every pair of genes in the genome is related in a specific tissue context. Thus, module detection provides a global, data-driven view of which genes are likely to participate in shared functions, pathways, and processes. Enrichment analysis is performed only after this data-driven partitioning, improving power to detect signals in the data and to implicate additional genes in biological processes based on their network connectivity. Thus our approaach uses multiple ontologies to fully map genes to functions.

### Cells of the kidney

#### Proximal tubular cells

Merged proximal tubule SCP networks predict a high level of metabolic activity dependent on β-oxidation of lipids, ammonium metabolism as well as absorption of ions, ion-dependent glucose reabsorption and detoxification mechanisms (Figure 6A). These SCPs, as shown by the different colors, are inferred from multiple technologies. The size of the SCP circle reflects the number of technology types that support the prediction of the SCP, while pie slices represent the individual technologies. In some physiology functions, cases of multiple pie slices are shown for the same technology indicating that this technology predicts the same SCP for multiple subtypes of the PT cells. The solid lines indicate connections between SCPs predicted by MBCO relationships and the dashed lines indicate additional well-known relationships between SCPs. Typically, these edges can represent functional relationships such as enzyme-substrate relationships or cotransport of molecules by symporters. It should be noted that most SCPs consist of multiple gene/gene products/metabolites of which only some are experimentally determined. Both the LMD proteomics and spatial metabolomics assays only distinguish between glomerular and tubulointerstitial regions in the kidney. SCPs that were predicted by these two assays either overlapped with or described similar functions as the SCPs that were identified by the proximal tubule cell or segment-specific datasets (Suppl. Figure 7). This agrees with the observation that most tubulointerstitial cells were proximal tubule cells (Suppl. Figure 4A/B). Consequently, we added all SCPs identified by LMD proteomics and spatial metabolomics to the proximal tubule network as well. The identified predictions are in agreement with the well-established physiological functions of PT cells that include ATP-dependent reabsorption of ions, glucose and other small molecules like amino acids and mono- and dicarboxylates (e.g., lactate or oxalate) (*33*). The pathways also highlight the important role of PT cells in ammonium excretion, drug clearance (*34*) and iron homeostasis pathways (*35*). The latter - among other functions - mitigate kidney damage during AKI (*36*). The prediction of glucose, fructose and glutamine metabolism from integration of transcriptomic, proteomic and metabolomics assays is in agreement with the high levels of PT gluconeogenesis activity (*37, 38*). Beta-oxidation, which is the central pathway for energy generation in the PT cells (*39, 40*), is predicted by four out of six technologies. The identified genes and proteins document involvement of both mitochondrial and peroxisomal beta-oxidation (Suppl. Table 7). These findings support the notion that peroxisomes could be a target in kidney injury (*41*) .

**Figure 6.**
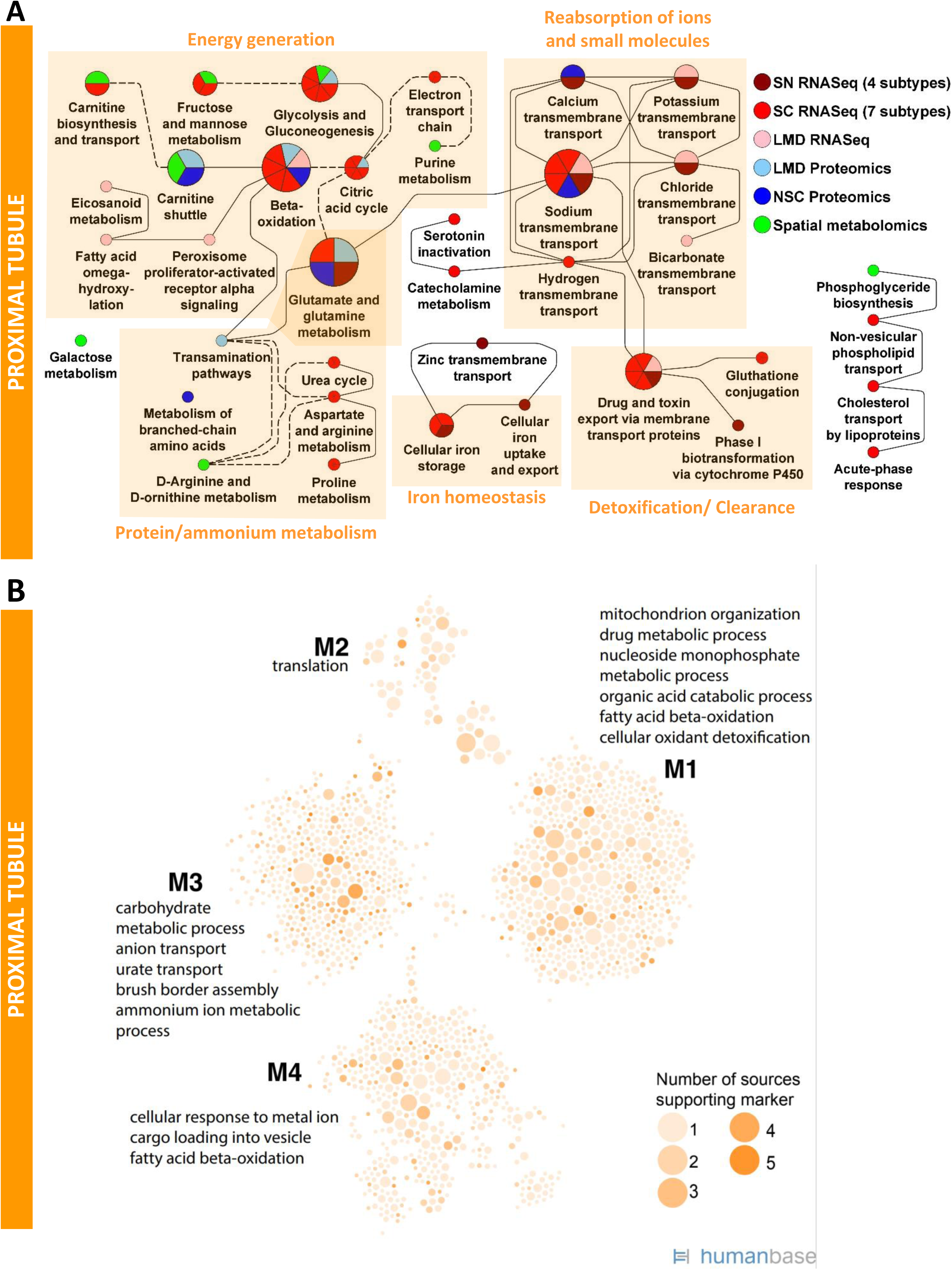
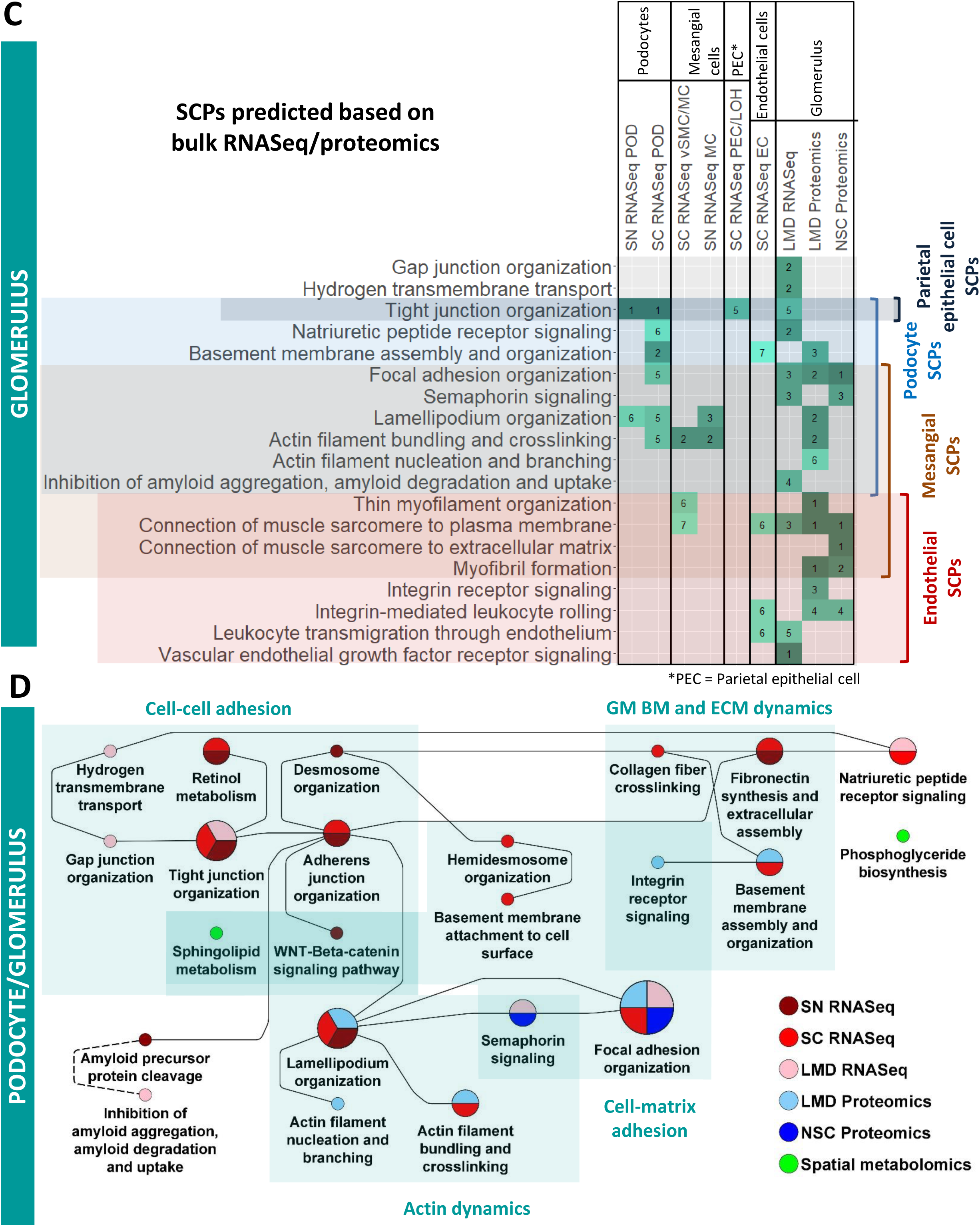
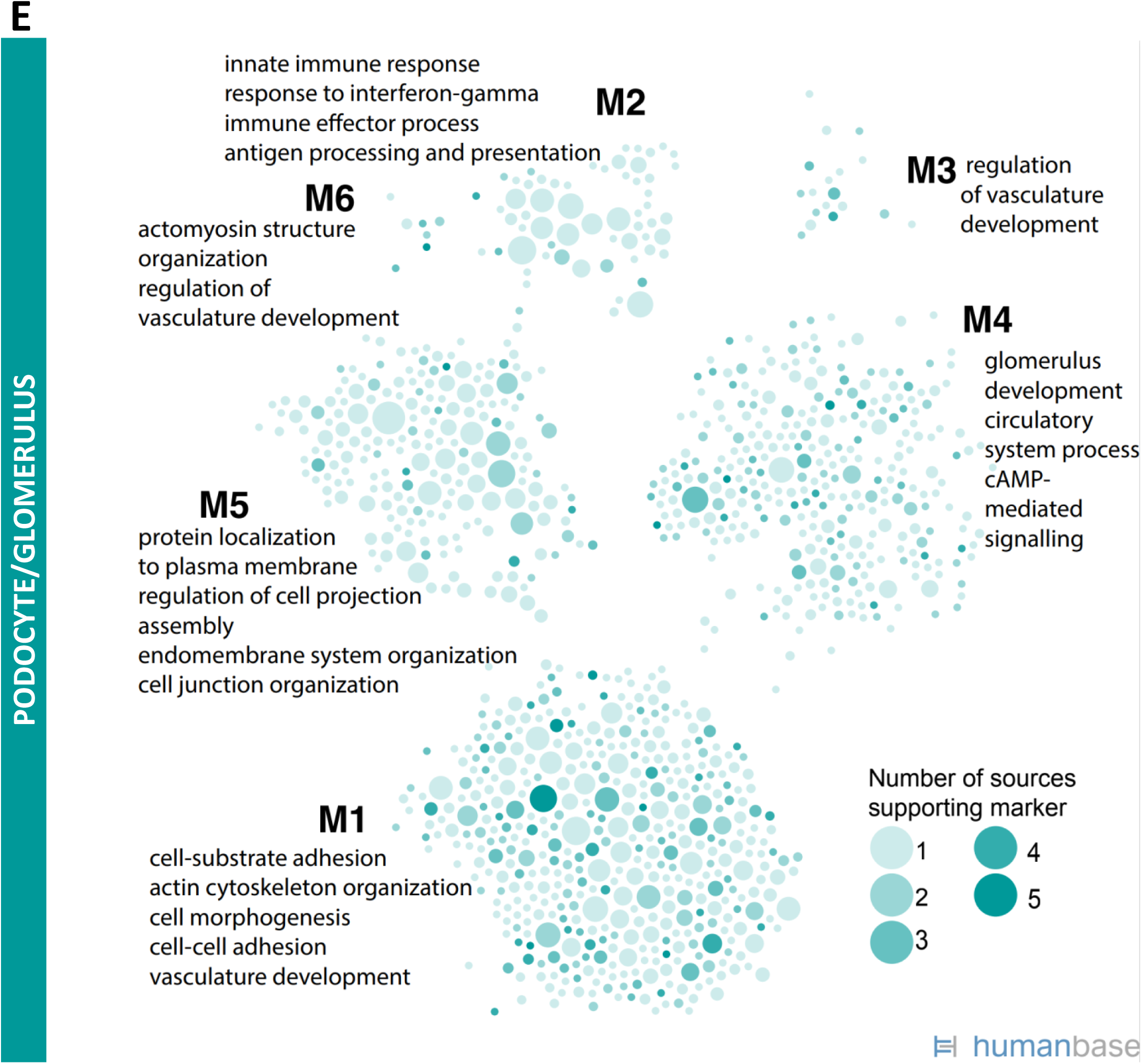
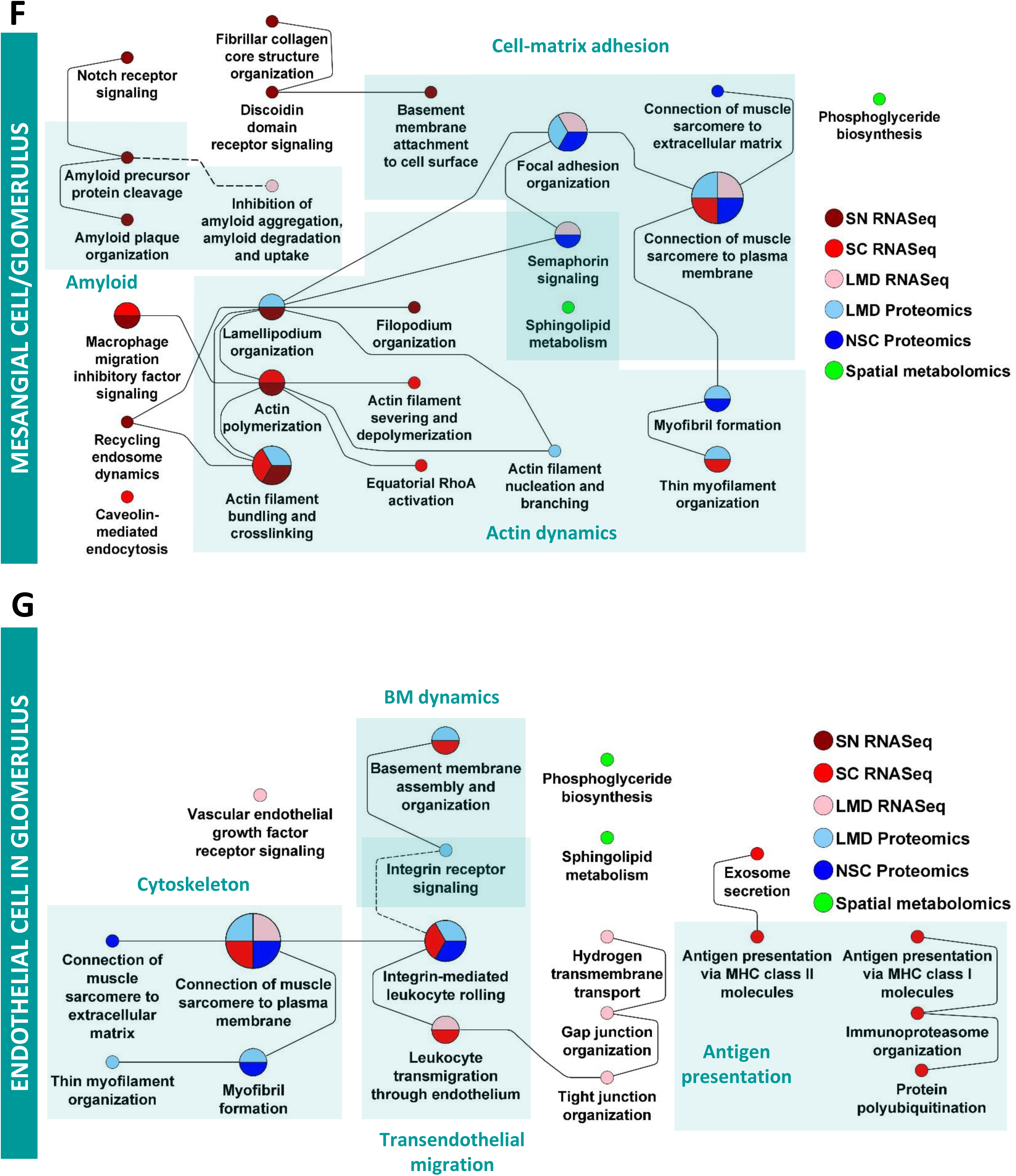
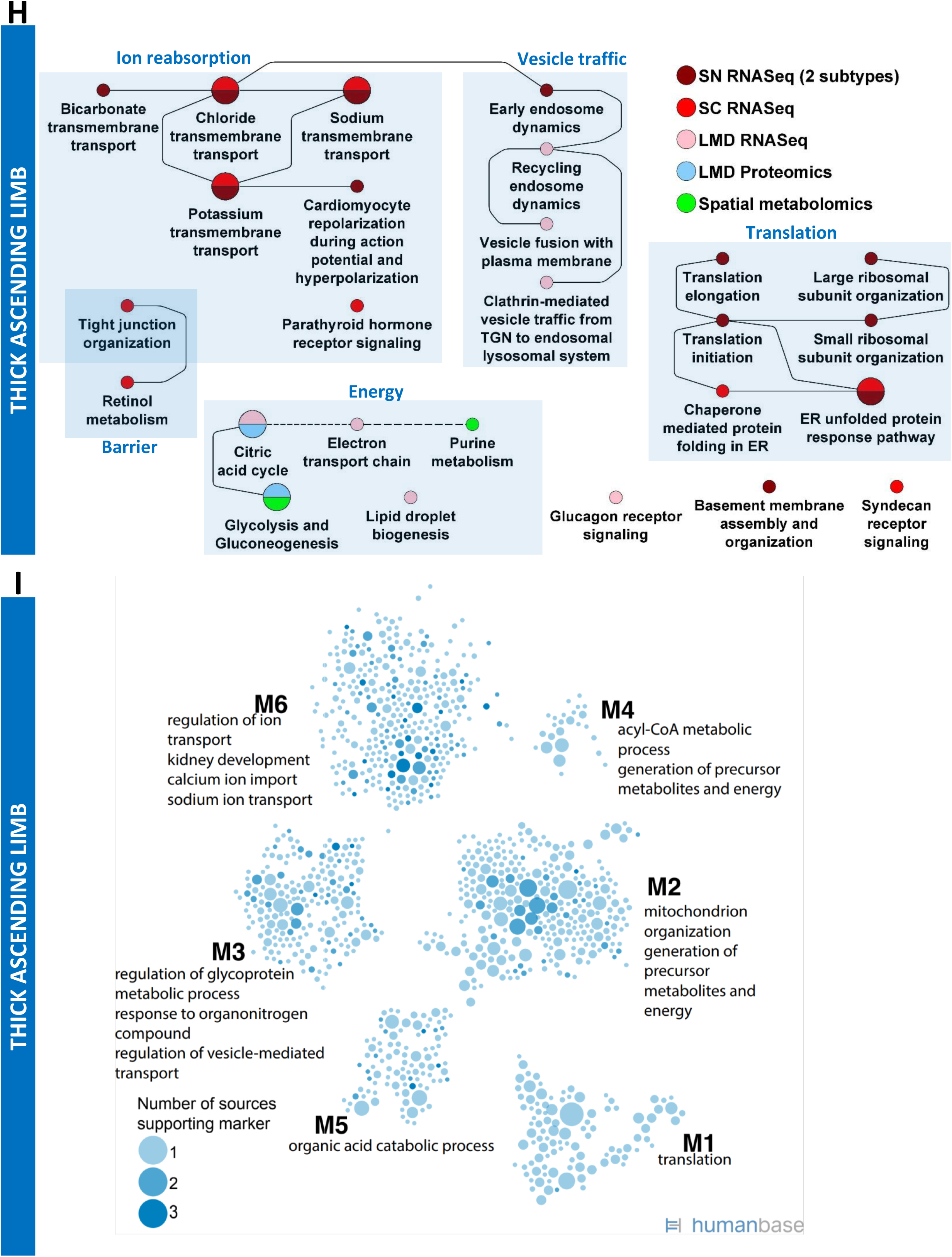
Enrichment analysis of markers for proximal tubule and glomerular cells and segments predicts cell well known functions. Nephrectomy tissues were subjected to single-nucleus (SN) and single-cell (SC) RNAseq, laser microdissected (LMD) RNAseq and proteomics, near single cell (NSC) proteomics and spatial metabolomics. **(A)** DEGs or DEPs of each PT cell subtype or subsegment were subjected to dynamic enrichment analysis using the Molecular Biology of the Cell Ontology (MBCO). Subcellular processes (SCPs) that were among the top seven predictions were connected by dashed lines, if their interaction was part of the top 25% inferred MBCO SCP interactions, and by dotted lines, if their functional relationship was curated from the literature. Supplementary figure 8 shows additional predicted SCPs involved in cell adhesion and translation. Metabolites associated with non-glomerular compartments were subjected to MetaboAnalyst enrichment analysis (Suppl. Figure 6). Any pathway among the top eight predicted pathways that was predicted based on metabolites specifically for that pathway was mapped to MBCO SCPs, if possible, and integrated into the PT SCP network. MBCO SCPs “Carnitine shuttle” and “Carnitine biosynthesis and transport” were added to the predicted MetaboAnalyst pathways, since four and two involved metabolites were among the non-glomerular metabolites (see methods for details). **(B)** Humanbase analysis of DEGs and DEPs. **(C)** SCPs predicted by dynamic enrichment analysis for the glomerular segment by the LMD RNAseq and Proteomics and NSC Proteomics assay were mapped to one of four detected glomerular cell types, because they were either detected in that cell type as well or related to SCPs detected for that cell type. Numbers indicate at which rank a particular SCP was detected. Notify that dynamic enrichment analysis can predict single SCPs or combinations of up to three SCPs, and consequently the same rank can be given to multiple SCPs. SCP network predicted for **(D)** podocytes, **(E)** mesangial cells and **(F)** glomerular endothelial cells by the sn and sc RNAseq datasets were merged with those SCPs that were predicted by the glomerular segment-specific datasets and assigned to each cell type as described above. **(G)** Podocyte specific modules were predicted by combined analysis of the podocyte sc and sn markers and the glomerular DEGs and DEPs. Predicted **(H)** SCP-network and **(I)** modules for the thick ascending limb cells/segment.

Both proteomic datasets of the PT subsegments highlight mitochondrial carnitine shuttle pathway that describes a central transport mechanism involved in both peroxisomal (*42*) and mitochondrial (*43*) beta-oxidation. We identify by spatial metabolomics the central carrier molecule carnitine, as well as acetyl-carnitine and palmitoyl-carnitine that are involved in transport processes during peroxisomal and mitochondrial beta-oxidation, respectively. The identification of carnitine biosynthesis and the carnitine precursor 3-Dehydroxycarnitine predicts that adult kidney - besides apical reabsorption of carnitine - also has the biosynthetic capacity for local carnitine production, as shown for human fetal kidney (*44*). Loss of beta-oxidation and consequently ATP synthesis is a significant contributor to tubulointerstitial fibrosis (*45*). Hence mapping of the variations in these pathways in different patient populations can provide a basis for molecular stratification of kidney fibrosis. Our data indicate the importance of beta-oxidation for proximal tubule function, since the prediction of local carnitine synthesis suggests an alternative carnitine source to dietary carnitine intake that might gain importance under a strictly vegetarian diet (*46*). Prediction of high levels of ATP generation and turnover rate is supported by the spatial metabolites that enrich for a pathway involved in the biosynthesis and degradation of adenine nucleotides. The ability of proximal tubule cells to significantly contribute to gluconeogenesis, especially in states of starvation (*47*) is documented by the identification of many enzymes involved in gluconeogenesis in our datasets. Glycolysis-specific enzymes were not detected, as described by others and in agreement with the low potential for glycolysis in the proximal tubule (*38*). Only a few pathways describing general cell biological functions (such as ECM dynamics, cell adhesion and translation) were predicted by one technology (Suppl. Figure 8).

Consequently, our analyses show that the different technologies describe the same biology, even though they might detect different genes or proteins and analyzed samples from the overlapping and non-overlapping participants (Suppl. Table 1).

Community clustering of PT marker genes in a kidney-specific functional network (Figure 6B) identifies four modules enriched for functions including translation (M2), cellular response to metal ion (M4), mitochondrial organization (M1), brush border assembly (M3), and anion transport (M3). The marker genes were identified across five distinct technologies (sc/sn/LMD transcriptomics, and two independent proteomics datasets), and include genes with a corrected p-value of less than 0.01 in each technology. Genes are shaded per number of technologies identifying each marker. Five genes (ALDH2, ANPEP, LRP2, PDZK1, and SHMT1) were identified as PT markers across all five technologies. Fifty-four genes were identified as PT markers by four of the five technologies, and 106 genes were identified as PT markers by three of the five technologies. Functional enrichments in module clustering provide a picture consistent with the SCP enrichments: key processes enriched in network modules and also identified in SCP enrichments include fatty acid beta-oxidation (M1, M4), ammonium ion metabolic process (M3), glucose metabolic process (M3), detoxification (M1), anion transport (M3), and cellular response to metal ion (M4). While we did not separate between male and female samples in this study, sex specific differences in proximal tubule cells have been described recently (*8*).

#### Glomerular cells

In agreement with a previous study focusing on human and mouse glomerular cells (*7*) we detected all four different glomerular cell types, podocytes, mesangial cells, endothelial cells and parietal epithelial cells. The sc and sn transcriptomic datasets (Figure 2) lead to four glomerular cell type specific SCP-networks. We separately analyzed the LMD transcriptomic and LMD and NSC proteomics and spatial metabolomics datasets (that were obtained from the whole glomerulus thus lacking cell type specificity) and identified glomerular SCP networks (Suppl. Figure 9A). Analyzing the overlap between the glomerular SCP networks with each of the three cell-type specific SCP-networks allows us to assign glomerular SCPs to podocytes, mesangial cells or glomerular endothelial cells (Figure 6C). Ten of the 19 glomerular SCPs are also predicted for at least one glomerular cell-type based on the sc/sn transcriptomic datasets. Seven other SCPs we identified map to particular cell types per functional relationships predicted from the sc/sn RNAseq datasets. These SCPs were added to each of the individual cell type specific SCP-networks. Podocyte SCPs (Figure 6D) focus on cell-cell/cell-matrix adhesion, glomerular basement membrane (GBM) and extracellular matrix (ECM) dynamics as well as actin dynamics. All these pathways are required for foot process maintenance and formation of the glomerular filtration barrier (*48–50*). Metabolomics data identify sphingolipid metabolism that could be involved in cell-cell adhesions as shown in other cell types (*51–54*). LMD segmental proteomics and transcriptomics identified key pathways involved in actin dynamics as well as cell-cell and cell-matrix adhesion. Multiple technologies identify tight junction organization, focal adhesion organization and lamellipodia organization. The glomerular slit diaphragm between mature podocytes develops from epithelial tight and adherens junctions (*55*). It contains many of these junctional protein components and was suggested to be a specialized form of either tight junctions (*56*) or adherens junctions (*57, 58*). This explains the prediction of these two structures from our data, thought they are not morphologically observed in healthy podocytes. We show WNT signaling as a central modulator of podocyte function (*59*). The pathway “Retinol metabolism” was predicted for both sc and sn RNAseq dataset as a regulator of tight junction similar structures. In agreement, retinoic acid has a regulatory effect on tight junctions in the epidermis (*60*) and plays a significant role in mitigating podocyte apoptosis and dedifferentiation during podocyte injury (*61*).

Community clustering of podocyte marker genes in a kidney-specific functional network identifies six modules (Figure 6E). Functional enrichments in these modules included glomerulus development (M4), vasculature development (M3), cell-substrate adhesion (M1), cell-cell adhesion (M1), and actin cytoskeleton organization (M1). Thirteen genes (*AHNAK*, *CLIC5*, *FERMIT2*, *GOLIM4*, *IQGAP2*, *NES*, *NPHS2*, *PDLIM5*, *PODXL*, *PTPRO*, *SLK*, *SYNPO*, and *TJP1*) were identified as podocyte markers by all five technologies surveyed. Forty-one genes were identified by four of the five technologies and 108 genes were identified by three of the five technologies.

Our datasets identify one mesangial and one transitional mesangial/VSMC cell type from the sn and sc RNASeq assays, respectively (Figure 2). LMD transcriptomics and proteomics and NSC proteomics along with sc and sn transcriptomics data identify SCPs involved in actin cytoskeleton dynamics, ECM dynamics, cell adhesion and amyloid plaque generation in these mesangial cells (Figure 6F). Our results are in agreement with their well-known function in blood vessel contraction and ECM support (*62*). In addition, one glomerular endothelial cell type was identified by the sc RNAseq data (Figure 2). Its SCP-network derived from integration of LMD proteomics and transcriptomics and NSC transcriptomics along with sc transcriptomic data identify cytoskeletal, trans-endothelial immune cell migration and antigen presentation pathways (Figure 6G). The assignment of “integrin-mediated leukocyte rolling” to endothelial cells is supported by the presence of the related “leukocyte transmigration through endothelium” SCP by sc and LMD RNA transcriptomics. Sn and sc RNAseq assays identified one parietal epithelial and one parietal epithelial cell type that also shows characteristics of loop of Henle cells, respectively (Figure 2). Parietal epithelial SCP networks contain pathways involved in cell-cell and cell-matrix adhesion and intermediate filament dynamics (Suppl. Figure 9B).

#### Loop of Henle

We identified one descending limb cell subtype by each sc and sn RNAseq assay (Figure 2). SCP networks from sc and sn RNAseq data for the descending limb cells identify cell adhesion functions and cytoskeleton dynamics (Suppl. Figure 10A). The presence of “tight junction organization” is in agreement with barrier formation in the descending limb that can allow for paracellular water reabsorption (*63*) but not for reabsorption of ions such as sodium or chloride (*64*). Community clustering of descending limb marker genes in a kidney-specific functional network identifies six modules enriched in functions including cell-cell adhesion (M6), epithelium development (M3), tube development (M3), response to endoplasmic reticulum stress (M5), and water homeostasis (M6) (Suppl. Figure 10B).

Three thin ascending limb (ATL) cell subtypes are identified by sn RNAseq although only one type was identified by sc RNAseq (Figure 2). SCP-networks obtained for ATL cells from these two technologies describe functions such as cell adhesion, cytoskeleton dynamics and translation (Suppl. Figure 10C). Overall, these SCP networks agree with the known functions of these cells that initiate the formation of dilute urine by the establishment of a water impermeable barrier that is permeable to low levels of ions (*65*). Community clustering of ATL marker genes in a kidney-specific functional network identifies seven modules enriched in functions including translation (M1), kidney morphogenesis (M6), and cell-cell adhesion (M4) (Suppl. Figure 10D).

Sc and sn transcriptomics identified one and two thick ascending limb (TAL) cell subtypes, respectively (Figure 2). TAL cell SCPs indicate sodium, potassium and chloride transport capabilities as detected by sc, sn and LMD transcriptomic technologies (Figure 6H). Tubulointerstitial SCPs identified by the LMD Proteomics and Spatial Metabolomics assays provide evidence for functional capabilities of the SCPs networks (Suppl. Figure 7). These findings are in agreement with the known transcellular reabsorption of sodium and chloride that is initiated by the furosemide sensitive sodium chloride potassium symporter NKCC2 and supported by apical potassium recycling (*66*). The “tight junction organization” SCP is involved in the establishment of a physical barrier that makes this region impermeable to water and thus allows the dilution of urine (*67*). Among the tight junction associated genes are *CLDN10* and *CLDN16* that are involved in the paracellular reabsorption of sodium or calcium/magnesium (*66, 68*), respectively, which supports the well-known physiology of this nephron segment. Involvement of “retinol metabolism” suggests that retinol regulated transcription can play an important role in TAL tight junction maintenance, similarly to its contribution to podocyte integrity. SCPs involved in the late secretory and early endocytic pathway support the known morphologic observation of vesicles below the plasma membrane that contain the furosemide sensitive NKCC2 (*66, 69*) allowing its mobilization and retrieval on demand (*70–72*).

The high energy demand of the TAL cells is reflected by the identification of SCPs involved in mitochondrial energy generation from LMD transcriptomics and proteomics. Spatial metabolomics that identify purine metabolites in the tubulointerstitium also support this conclusion. Community clustering of TAL marker genes in a kidney-specific functional network (Figure 6I) identifies six modules enriched in functions including regulation of ion transport (M6), calcium ion import (M6), sodium ion transport (M6), translation (M1), and mitochondrion organization (M2).

#### Distal convoluted tubules

One distal convoluted (DCT) cell subtype was identified based on each of sc and sn RNAseq assays (Figure 2). Predicted SCPs for the DCT cells from sc, sn and LMD transcriptomics converge on sodium and chloride transmembrane transport (Suppl. Figure 11A). Our results agree with the well-known sodium and chloride reabsorption by this cell type via the thiazide sensitive sodium chloride symporter NCC (*73*). Additionally, sc/sn transcriptomics highlight reabsorption of calcium, potassium, bicarbonate and phosphate. Community clustering of DCT marker genes in a kidney-specific functional network (Suppl. Figure 11B) identifies three modules enriched in functions including regulation of ion transport (M3) and metal ion homeostasis (M2). A recent study focusing on the cells in the distal nephron purified by FACS-enrichment of mouse kidney cells further classifies the DCT cells into multiple subtypes (*4*).

#### Connecting tubules

Each sn and sc assay identified one connecting tubule (CNT) subtype (Figure 2). Both sn and sc transcriptomic datasets for CNT cells indicate that SCPs for sodium, potassium and calcium transmembrane transport activities are enriched (Suppl. Figure 12A), supporting its function in fine tuning electrolyte balances (*74–76*). Other SCPs indicate signaling, endoplasmic reticulum and energy functions in this cell type. Community clustering of CNT marker genes in a kidney-specific functional network (Suppl. Figure 12B) identifies three modules enriched in functions including ion transport (M2), receptor-mediated endocytosis (M3), and mitochondrion organization (M1).

#### Collecting duct

Sc and sn RNAseq show two and three principal cell subtypes, respectively (Figure 2). The principal cell SCP networks were obtained by merging the principal cell specific SCPs predicted from sc and sn transcriptomics with the collecting duct (CD) specific SCPs predicted from LMD transcriptomics (Suppl. Figure 12C). Overlapping or functionally related SCPs identified by LMD Proteomics and Spatial Metabolomics were added as well (Suppl. Figure 7). Both sc and sn technologies identified “Potassium-“ as well as “Sodium-transmembrane transport” SCPs for the principal cells. The SCP ‘Water transmembrane transport’ was identified by both sn and sc RNAseq assays as well, though with a lower rank for sn RNASeq assays that did not pass our applied cutoff. The LMD transcriptomics and proteomics data identified the energy generation SCPs required for the various transport SCPs identified by the sc and sn transcriptomic data. The spatial metabolomics data sets provided support for energy generation pathways identified by the LMD technologies.

Principal cells play an important role in fine tuning ion and water reabsorption and thereby regulate systemic electrolyte and water balance (*76*). The anti-diuretic hormone working with prostaglandins regulates the levels of AQP2 on the apical plasma membrane (*77, 78*) stimulating water reabsorption by the principal cell. Apically reabsorbed water is exported by basal water transporters AQP3 and AQP4. We detect both *AQP2* and *AQP3* in our datasets. Sodium reabsorption is regulated by the amiloride-sensitive sodium channel EnaC whose expression and protein turnover is regulated by aldosterone (*79*). The aldosterone-stimulated reabsorption of sodium is coupled with secretion of potassium (*80*), as highlighted by our data. Additionally, we show calcium transmembrane transport for one cell subtype by both sn and sc RNAseq assays. Both sc and sn technologies identify SCPs involved in drug and toxin transmembrane movement in one of the subtypes of the principal cell, although drug excretion is generally described to occur in the proximal tubule (*34*). Furthermore, community clustering of PC marker genes in a kidney-specific functional network (Suppl. Figure 12D) identifies seven modules enriched in functions including ion transport and homeostasis (M7), regulation of vesicle-mediated transport (M4), and water homeostasis (M6).

We identified multiple subclusters of intercalated cells that could be assigned to IC-A, IC-B and one transitionary subtype, tPC-IC, as well as IC-A1, IC-A2 and IC-B in the sc and sn transcriptomic datasets, respectively (Figure 2). SCPs networks were identified by merging sc and sn transcriptomic data with LMD transcriptomic data obtained from the collecting duct (Suppl. Figure 12E). Additionally we added overlapping or functionally related SCPs predicted by LMD Proteomics and Spatial Metabolomics (Suppl. Figure 7). We find the SCP “Bicarbonate transmembrane transport” in all three sc subtypes and one sn subtype (Suppl. Figure 12E), documenting the importance of the intercalated cells in the regulation of systemic acid-base homeostasis (*81*). Apical and basolateral bicarbonate transport is driven by exchange for chloride (*81*), as indicated by the “Chloride transmembrane transport” SCP identified for one subtype in both sn and sc RNAseq datasets. Community clustering of IC marker genes in a kidney-specific functional network (Suppl. Figure 12F) identifies six modules enriched in functions including regulation of body fluid levels (M3), translation (M1), mitochondrion organization (M2), bicarbonate transport (M5), and cell-cell adhesion (M4). Enrichment analysis using Gene Ontology predicts phagocytic activity (phagosome maturation and acidification) based on subunits of the vacuolar H^+^ATPase (*81*) (Suppl. Figure 12G). In combination with the prediction of SCP involved in actin cytoskeleton our data supports the recent observation of phagocytic activity of the intercalated cells (*82, 83*).

#### Interstitium and the vasculature

##### Endothelial Cells

We find four types of endothelial cells by sn transcriptomics and two by sc transcriptomics, in addition to glomerular endothelial cell identified sc transcriptomics (Figure 2). SCP networks for endothelial cells identified from sc and sn transcriptomic data sets contain pathways involved in cellular adhesion, trans-endothelial migration, actin cytoskeleton dynamics, caveolin-mediated endocytosis, signaling and antigen presentation (Suppl. Figure 13A).

##### Vascular smooth muscle cells

We identified a single type of VSMC by sn RNAseq assay (Figure 2). The sc transcriptomic technology identified a variant of mesangial cells that has VSMC markers. We classified this subtype as a glomerular cell subtype. SCP networks from sn technology highlight cell contraction capabilities for the VSMC (Suppl. Figure 13B).

##### Fibroblasts

We identified a single type of fibroblast from sc and sn RNAseq assays (Figure 2). SCPs in fibroblasts identified from sc, sn and LMD transcriptomics data describe pathways related to ECM dynamics, cell adhesion, cytoskeleton dynamics and the complement pathways (Suppl. Figure 14). The proteomic assays did not detect ECM components related SCPS among the highly ranked pathways.

##### Immune cells

Four types of immune cells are detected by sc or sn RNAseq technologies. These include natural killer cells, three types of T-cells, B-Cells and three types of macrophages and monocytes (Figure 2). SCP-networks for macrophages contains pathways involved in antigen presentation, actin cytoskeleton dynamics and translation (Suppl. Figure 15A). Connection of the SCPs involved in actin cytosekeleton dynamics to the SCP ‘Macrophage migration inhibitory factor (MIF) signaling pathway indicates the potential for chemotactic activity. Macrophage migration is driven by rearrangements in the actin cytoskeleton that are activated by stimulation of the MIF receptor proteins CD74 and CXCR4 (*84, 85*) as identified in our data. The SCP ‘Cellular iron uptake and export’ documents the central role of macrophages in iron homeostasis (*86*). It is predicted based on SLC39A8, a transmembrane transporter involved in transport of multiple divalent metal ions including iron (*87*) and the scavenger receptor CD163 that is also involved in removing hemoglobin or haptoglobin-hemoglobin complexes by splenic red pulp macrophages and Kupffer cells (*88*). This SCP and the SCPs involved in actin dynamics are also identified by LMD transcriptomics of the interstitium. SCPs in the natural killer cells identify antigen presentation, cell migration and actin cytoskeleton dynamics (Suppl. Figure 15B). Similarly, SCP-networks predicted for B-cells and T-cells contain pathways involved in antigen presentation and the immunoproteasome and translation (Suppl. Figure 15C and 15D, respectively). A detailed study of immune cell zonation of the human kidney has been published (*14*), while another single cell sequencing study characterized twelve myeloid cell subtypes associated with progression and regression of kidney disease in an animal injury model (*6*).

Since immune activity was documented for all cell types along the nephron (*14*), we analyzed the fraction of cell type and subtype specific marker genes and proteins that were annotated to immune pathways in Gene Ontology. In agreement with the indicated study, about 5-15% of all marker genes participate in immune cell functions (Suppl. Figure 16). We want to emphasize that in the immune zonation study (14) the highest immune activity was predicted for epithelial cells of the pelvis, while our samples do not contain tissue from the pelvis.

#### Using the atlas to understand the molecular basis of physiological functions

Using a similar approach to our *post hoc* power analysis, we investigated the robustness of the SCP-identified cell biological functions by randomly downsampling libraries from the sc/sn datasets. We subjected the top 300 DEGs of each cell type and subtype in each downsampled dataset to dynamic enrichment analysis. Predictions were ranked by significance. Investigation of the ranks that were obtained for those SCPs in the down-sampled datasets that are among the top seven predictions in the full dataset allows to estimate which SCPs are consistently predicted and probably describe biological core functions. The most consistently predicted SCPs share a high overlap with those SCPs that are predicted from multiple datasets, as described above. In case of the proximal tubule cells, most of the consistently identified SCPs by the sc (Suppl. Figure 17A) or sn RNAseq data (Suppl. Figure 17H) are related to cellular metabolism and energy generation, reabsorption and detoxification. In the case of the podocytes, the consistently identified SCPs are involved in cell-cell and cell-matrix adhesion. ‘Tight junction organization’ and ‘Hemidesmosome organization’ are consistently identified based on the sc (Figure 17B) and ‘Tight junction organization’ and ‘Adherens junction organization’ on the sn RNAseq assay (Figure 17I). These results document the central importance of the glomerular slit diagram that is described as a specialized form of both tight junctions (*56*) and adherens junctions (*58*). Supplementary figure 17 also shows the results obtained for the other cell types.

#### Comparison of variation of oxygen supply and inferred levels of energy metabolism help understand sites of kidney injury

To identify energy generation pathways in the different cells along the renal tubule of the nephron, we generated a focused ontology of metabolic pathways (Suppl. Figure 18A). Our ontology focused on the design of small pathway units that distinguished between reactions specific for a particular pathway (e.g., enzymatic reactions that participate in glycolysis, but not in gluconeogenesis) and reactions shared by two or more pathways (e.g., enzymatic reactions shared by glycolysis and gluconeogenesis). The different pathway units converged on more general parent pathways that contained all set of reactions involved (e.g., specific and shared reactions involved in glycolysis). Pathways were populated by literature curation, parent pathways inherited the genes of their children. Enrichment analysis of cell type, subtype and segment specific marker genes and proteins. Using this ontology allowed us to distinguish between aerobic and anaerobic as well as catabolic and anabolic pathways (Suppl. Figure 18B). To rigorously define the groups, we only considered a parent pathway if its child contains the reactions specific for that pathway among the predictions.

The patterns of expression of different pathways involved in aerobic and anaerobic energy generation along the nephron (Figure 7, Suppl. Table 11) and the varying levels of oxygen availability in the different regions of the nephron (*89*) are shown. This comparison helps us identify potential regions with differential susceptibilities towards hypoxia induced kidney injury. Missing capability for anaerobic energy generation as predicted by the transcriptomics data in humans (Suppl. Figure 18B) and observed in animal experiments (*90*) combined with low pO_2_ suggests S3 segment of the proximal tubule as a site for hypoxia based injury. This reasoning is in agreement with experimental observations (*89*). High levels of capacity for aerobic energy generation activity in the medullary TAL (mTAL), a region with low oxygen supply, is complemented by high capacity for anaerobic energy generation, as also documented in animal experiments (*90*). When the output of the anaerobic energy generation is depleted then the mismatch between oxygen availability and aerobic glycolysis can lead to the accumulation of intermediates that can damage the region. Thus, our conclusions on TAL agrees with the experimental observation that mTAL injury during hypoxia depends on epithelial transport activity (*89*). It can be readily seen that molecular profiles of metabolic pathways in the atlas provide a basis for understanding and predicting kidney injury due to hypoxia.

**Figure 7:**
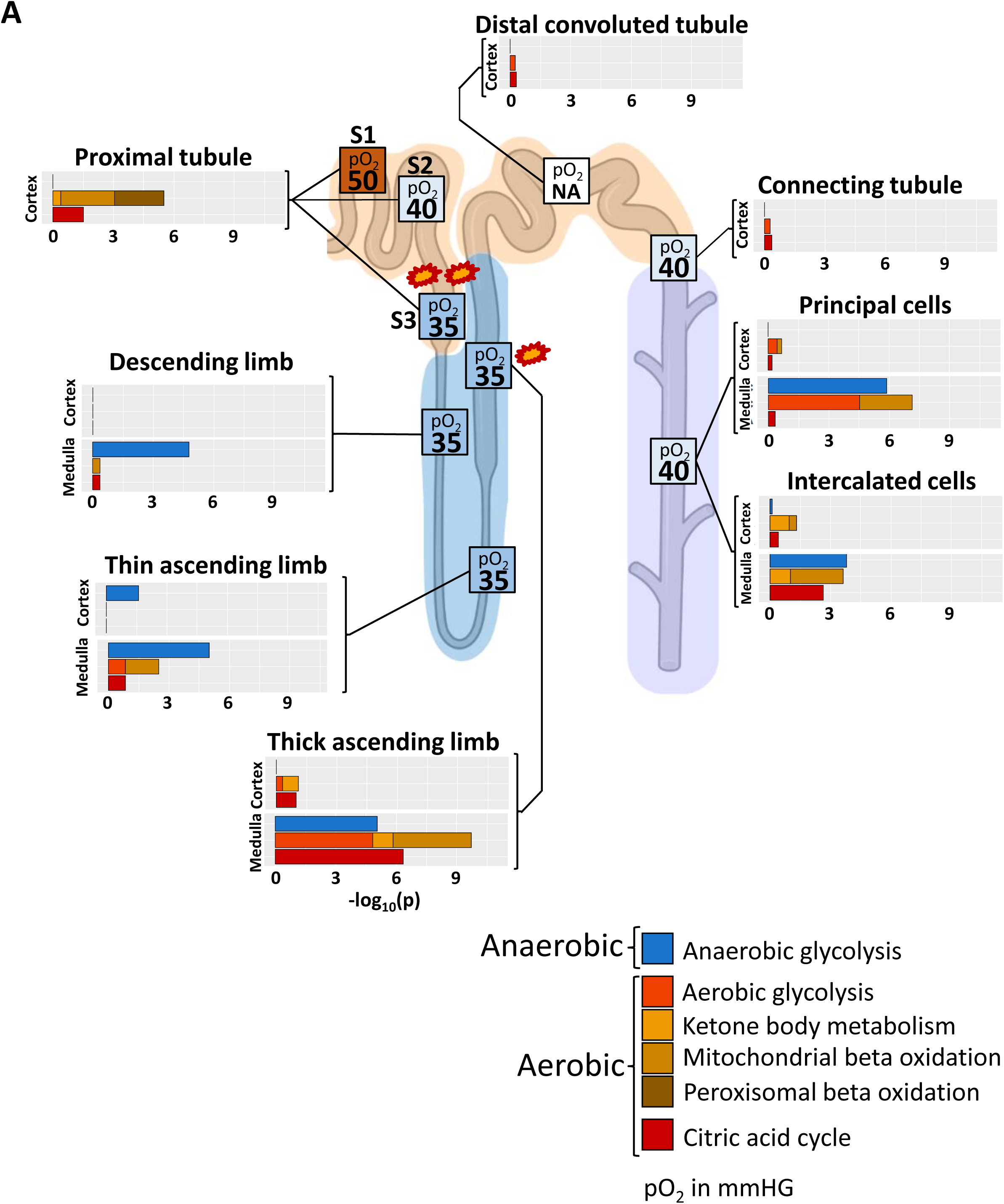
Aerobic and anaerobic energy generation profiles and oxygen supply accurately highlights sites of hypoxia induced injury. To compare energy generation profiles with experimentally determined oxygen supply in the different nephron regions, we generated an ontology that allows the separation of aerobic and anaerobic pathways involved in energy generation. Enrichment analysis of cell type, subtype and subsegment marker genes with this ontology predicts high dependency of proximal tubule cells on aerobic energy generation, suggesting S3 as a primary injury site during hypoxia (marked by two explosions) because of its low oxygen supply under basal conditions. Enrichment results predict a high aerobic energy generation activity for the medullary TAL that can be compensated by anaerobic energy generation. In combination with the already low oxygen saturation in that segment under normal conditions our results suggest that mTAL is the second, though less likely, injury site during hypoxia (marked by one explosion). Enrichment results are combined from those shown in Suppl. Figure 17B. Numbers in boxes indicate pO2 in mmHg taken from (*89*), NA: not available.

Although podocytes are capable of generating energy by anaerobic glycolysis (*91*), we did not identify any podocyte marker genes involved in any of the analyzed energy generation pathways. Since maker genes were determined by comparing cell subtype, type or segment specific gene expression to expression profiles in all other cells or segments, our analysis does not document that these genes are absent in podocytes, but only that they are not expressed in podocytes at higher levels as compared to other kidney cells.

#### Comparision of the physiological activity along the nephron and mRNA levels of transporters provide an understanding of the molecular basis for differential physiological activity

Physiological experiments allow us to determine how much of a filtered ion or small molecule is reabsorbed in a particular nephron segment. Results of these experiments are described in standard medical school physiology textbooks. Typically, the results shown in these textbooks specify the percentage of a filtered ion or molecule reabsorbed in a particular nephron segment (such as the proximal tubule) or finally excreted into the kidney pelvis and ureter. Sodium reabsorption is important for blood pressure control and hence we focused on sodium reabsorption as an example of how a cell level tissue atlas that details levels of the various sodium transporter genes can help us understand physiological homeostasis.

We obtained sodium reabsorption profiles from 4 different standard physiology text books (*67, 92–94*) and averaged the reabsorption percent values for each ion or molecule (Figure 8A-1). An estimated fraction of 1/3 to 2/3 of the total sodium reabsorbed in the proximal tubule is reabsorbed by passive paracellular mechanisms (*95*). Ablation of the tight junction protein claudin2 that facilitates paracellular sodium transport reduces sodium reabsorption in the proximal tubule by 37% (*96*). Paracellular sodium reabsorption in the Loop of Henle is estimated to be below 50% based on electrophysiological considerations (*95*), approximately around 30%. Since we wanted to compare the experimentally determined reabsorption profiles with mRNA levels of the different sodium transporters that mediate transcellular reabsorption, we removed 37% and 30% from the experimental reabsorption profiles for the proximal tubule and Loop of Henle, respectively. Obtained physiology experimental values were readjusted to sum up to 100% to document how much of transcellularly reabsorbed sodium is reabsorbed in each nephron segment (Figure 8A-2). We then calculated the sum of all mRNA levels of plasma membrane transporter genes for sodium for all cells in each segment of the renal tubule of the nephron, using an ontology of kidney sodium transmembrane transport that we generated from prior knowledge (Suppl. Figures 19A/B/C). To determine if results are consistent across data sets, we calculated the mRNA levels from three sn RNAseq datasets, our sn RNAseq dataset (*20*) and two additional sn RNAseq (*97, 98*) datasets from the Humphries laboratory. We hypothesized these sums to represent the total sodium transport capacity for each segment. Detailed methods and assumptions underlying this hypothesis are provided under Methods. We compared the mRNA levels from the sc and sn transcriptomic experiments with the experimentally measured reabsorption profiles of sodium along the nephron without (Figure 8B-1) and with (Figure 8B-2) removal of paracellular sodium reabsorption. There is agreement between the levels of sodium reabsorption seen in physiology experiments with the mRNA levels in the different cell types along the nephron. We see differences mainly in the Loop of Henle. Under consideration that there is most likely spare capacity for sodium reabsorption in the Loop of Henle (*66, 69–72*), our data documents a good agreement between the calculated sodium reabsorption capacities and the experimentally measured reabsorption profiles. Since some nephron segments, such as Loop of Hele and collecting duct contain multiple cell types with different reabsorption mechanisms (*66, 99*) we decided to focus on the different cell types to determine the contribution of different gene products to the overall inferred transport capacity. The relative distribution of mRNAs encoding the different transporter proteins for sodium is shown in Figure 8C. Since some of the mRNA mapped to SCPs that are involved in blood-to-lumen transport (Supp. Table 12), we defined these mRNA levels as negative to account for the opposite direction when compared to lumen-to-blood transport. Consequently, when we add the mRNA levels of all of these individual transporters along each nephron segment, we obtain the total fraction of inferred sodium transport capacity of each segment as documented in figure 8B. A significant contribution of sodium channels to the fine-tuning of sodium reabsorption in the collecting duct (*22*) could explain, why the mRNA levels associated with sodium transporter involved in blood-to-lumen transport are higher than the levels involved in lumen-to-blood transport in this subsegment.

**Figure 8:**
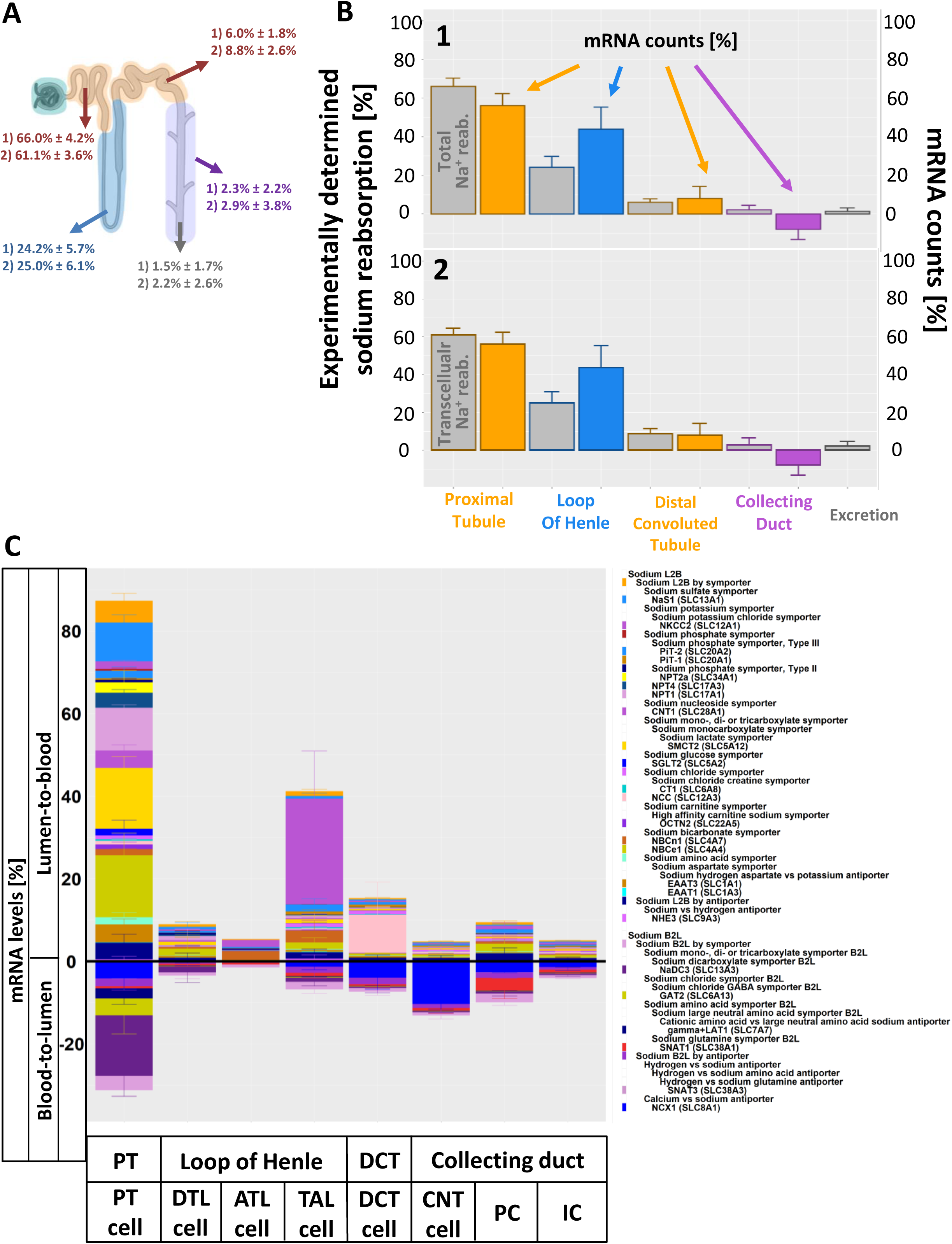
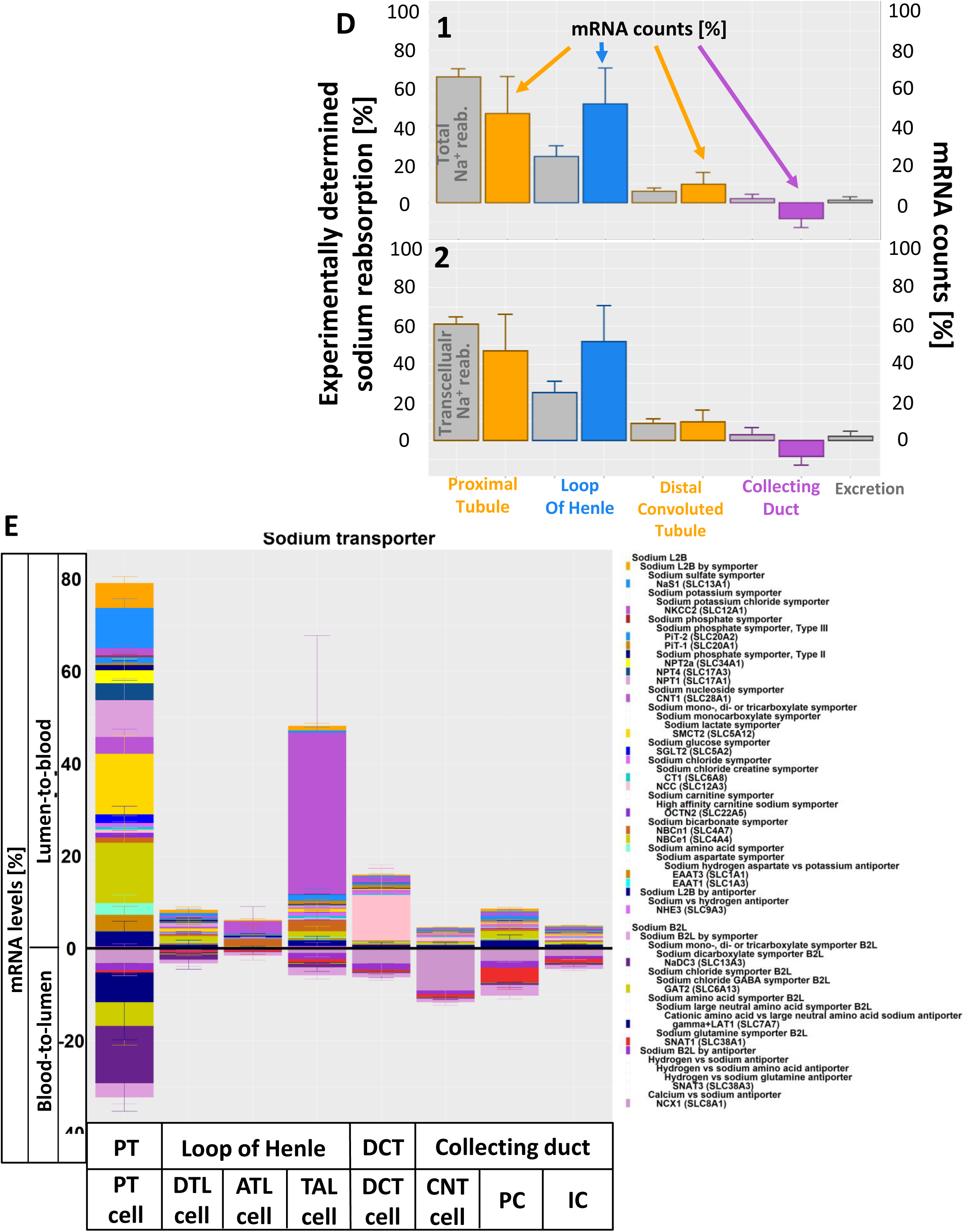
Predicted sodium transport capacities match with experimentally determined reabsorption profiles. **(A)** Sodium reabsorption profiles that document the percentage of the glomerular filtered sodium in each segment of the nephron were obtained from four different standard physiology and medical text books (*92–94, 120*), followed by calculation of the mean and standard deviation values. These values for total reabsorption are shown in (1). About 37% of the sodium reabsorbed in the proximal tubule is paracellular(*96, 134*), while paracellular sodium reabsorption in the Loop of Henle is estimated to be below 50% (*95*). Since we want to compare the experimentally determined sodium reabsorption profiles with mRNA levels involved in transcellular sodium reabsorption, we subtracted 37% and 30% paracellular transport values from the total sodium reabsorption values in the proximal tubule and Loop of Henle, respectively. Profiles were readjusted, so that they sum up to 100%, followed by calculation of means and standard deviations for each segment. These values are shown in (2). **(B)** To compare predicted reabsorption capacities from mRNA levels with those from physiological experiment derived reabsorption profiles, we generated an ontology that assigns genes to the different subcellular processes responsible for sodium movement by different transporter proteins. All transport processes were integrated into a hierarchy that finally converges on lumen-to-blood and blood-to-lumen transport for sodium. For each of three different sn RNAseq datasets (one from KPMP (*20, 21, 135*) and two from the Humphrey laboratory(*97, 98*)) obtained for reference tissue we calculated the sum of all mRNA counts that mapped to genes involved in lumen-to-blood or blood-to-lumen transport of Na+ for each segment of the renal tubule. The net reabsorption capacity for sodium was determined by calculating the difference between both mRNA count levels. Segment specific net reabsorption capacities are expressed in percent of total reabsorption along the nephron, followed by calculation of mean and standard deviations. Physiological experiment derived reabsorption profiles determined in (A) with (I) and without (II) paracellular sodium reabsorption in the proximal tubule are shown in gray. **(C)** Cell type specific transport capacities for sodium. Segment associated single cell or nucleus read counts were summed up for selected transport processes involved in sodium transmembrane transport, followed by normalization of the results towards the net lumen-to-blood transport capacity (see figure (B) for details). Normalized transport capacities from the single cell and single nucleus RNAseq datasets were averaged. All transport capacities that described lumen-to-blood or blood-to-lumen transport were assigned to be positive or negative, respectively. Notify that the positive bars documented in (C) are the sums of all (positive) lumen-to-blood and (negative) blood-to-lumen mRNA levels of each segment. Parent-child relationships between shown subcellular processes (SCPs) are documented in the legend, where children SCPs are written below their parent SCPs and shifted to the right. Parent SCPs only contain mRNA levels mapping to genes that are not assigned to any of the documented children SCPs. For a proper documentation of the SCP hierarchy, we added all offspring SCPs of every selected SCP to the legend, even if the offspring SCPs were not selected. Unselected offspring SCPs whose mRNA levels are not shown in the bar diagrams are not annotated to a color in the legend. Their mRNA levels are part of the next selected ancestor SCP. Notify that in case of multiple parent SCPs we only show one parent in the legend that was arbitrarily selected (Supplemental Table 3). (E) Comparison between reabsorption profiles and reabsorption capacities that were predicted from the 3 sn RNAseq datasets used above and the KPMP sc RNAseq dataset. (F) Sodium reabsorption mechanisms were predicted based on all 4 datasets. See figure 2A for cell type abbreviations.

Calculation of the reabsorption capacities after inclusion of sc RNAseq dataset along with the sn RNAseq data, slightly decreased the match with the physiological reabsorption profiles and mRNA levels (Figure 8D). This is mainly due to the high mRNA levels associated with the basolateral amino acid transporter gamma+LAT1 that exports cationic amino acids into the blood in exchange for flow of large neutral amino acids and sodium into the cell (*100, 101*) (Figure 8D).

Regarding the transport mechanisms involved in sodium reabsorption (Figure 8B/D), we highlight two details here. The proximal tubule is the primary region of the nephron for absorption of many metabolites including different amino acids, organic anions and sugars. Often this absorption is coupled to sodium transport. We identified a large number of distinct gene products that that are responsible for these transport processes. We have grouped these transporters together based on the SCP hierarchy for total transcellular sodium transport. It is this sum of all the mRNA levels that matches the total transcellular sodium transport. Thus, the cell level atlas provides a detailed picture that was up to now not attainable. The second noteworthy feature is that NKKC2 is the major sodium transporter in the TAL cells of the Loop of Henle (*66*). In agreement with this cell physiological knowledge, mRNA encoding NKCC2 is the predominant species of sodium transporter in the TAL cells.

The distribution of mRNAs for the various transporters agrees well with the known levels of reabsorption activities identified in physiological experiments. This agreement suggests that the levels of transporter mRNAs at the single cell level can provide an indication of transcellular transport capacity of the cell type. In support of this conclusion, we find that glucose transport along the nephron agrees with glucose transporter mRNAs (Suppl. Figure 19D/F) and is mainly mediated by SGLT2 (Suppl. Figure 19E/G) that is responsible for ≥ 80% of filtered glucose reabsorption in the proximal tubule (*102*).

## Discussion

The integration of multiple types of omics data allows us to describe in depth multiple subcellular processes and pathways at cell level resolution. From such description we can hypothesize key functions, that when perturbed define disease states. Such disease states could have convergent clinical phenotypes even though the underlying molecular changes are different. Thus, our detailed characterization described here can provide the starting point for a new framework for molecular classification of kidney diseases. For example, the identification of both mitochondrial and peroxisomal β-oxidation and carnitine transport and local biosynthesis pathway in PT cells suggest how individual variations in any of these SCPs can contribute to the effects of kidney injury including fibrosis (*9*). Thus, a convergent clinical phenotype can arise from very different molecular changes related to energy metabolism. Mapping these changes in individual patients may allow for better classification of disease states.

### Integrated view of kidney cellular functions

One advantage of the presented multiomics data integration strategy is the ability to infer how different classes of biomolecules may enable complex multicellular functions leading to potentially predictive biomarkers. Spatial metabolomics identifies N-Palmitoylsphingomyelin (SM d18:0/16:1) as a spatial correlate of glomerular kidney segments (correlation coefficient > 0.9; Figure 9A), as described previously (*103, 104*). To identify the cell types involved in its synthesis, we screened all glomerular cell types for expression of genes involved in ceramide, sphingomyelin and sphingosine metabolism (*105, 106*). Transcriptomics identify *SERINC5* (serine incorporator 5) and *CERS6* (ceramide synthetase 6) as specifically expressed in podocytes or in podocytes and mesangial cells, respectively (Figure 9B). SERINC5 incorporates serine into the membrane of the endoplasmic reticulum, making it available for ceramide and phosphatidylserine synthesis (*105*). *CERS6* is identified by all transcriptomic assays in the podocyte. CERS6 is one of six ceramide synthases that converts sphingosine and acyl-CoA into ceramide. In contrast to the other five ceramide synthases, it has a high substrate specificity towards palmitoyl-CoA (C16:0) (*106*), thereby generating ceramides with the correct acyl chain length to be converted into SM d18:0/16:1. Only one technology, sn RNAseq, shows *CERS6* to be expressed in mesangial cells as well, albeit at lower level of significance (rank 293 in mesangial cells vs 116 and 126 in podocytes; Suppl. Figure 20). Transcriptomic datasets also predict the expression of enzymes involved in sphingomyelin synthesis in non-glomerular cells. In general, only one specific enzyme of this pathway is expressed per cell type. Consequently, podocytes (and mesangial cells) are the most likely synthesis site for this particular sphingomyelin as demonstrated by the spatial metabolomics data. Altered metabolism of multiple sphingolipids including sphingomyelin and its metabolites is observed in several glomerular diseases (*107*). Cellular sphingomyelin is predominantly localized at membranes derived from the trans-Golgi and plasma membrane (*108*). It is involved in multiple functions, including cell signaling (*109*), lipid rafts formation (*110*), caveolar endocytosis (*108*) and apoptosis (*111*). Additionally, it has long been known that sphingomyelin (as part of lipid rafts) is enriched in desmosomes (*51, 110*) and tight junctions (*54*). Given the central importance of foot process interactions between neighboring podocytes, the potential importance of this metabolite in making different types of cell-cell contacts and the enzymes involved in its biosynthesis in podocytes can be readily appreciated. Five technologies that focus on genes and proteins identify cell-cell/cell-matrix adhesion as podocyte key functions, and the sixth technology identifies a specific metabolite that we consequently predict would be involved in the same key functions thus providing integrated support for the role of sphingomyelins in podocyte cell-cell interactions. Sphingomyelin with very long (C24) acyl chains, but not palmitoyl (C16) sphingomyelin is the predominant species associated with tight junctions (*54*). Tight and adherens junctions are morphologically only observed in developing or diseased podocytes (*55, 112*), while in healthy podocytes they have morphed into the glomerular slit diagram (*55*). These observations indicate that we may be able to use decrease in levels of C16 SM and increase in long chain SM as a predictive biomarker for disease progression even prior to changes in glomerular filtration rates. Such hypotheses may be tested in the future.

**Figure 9:**
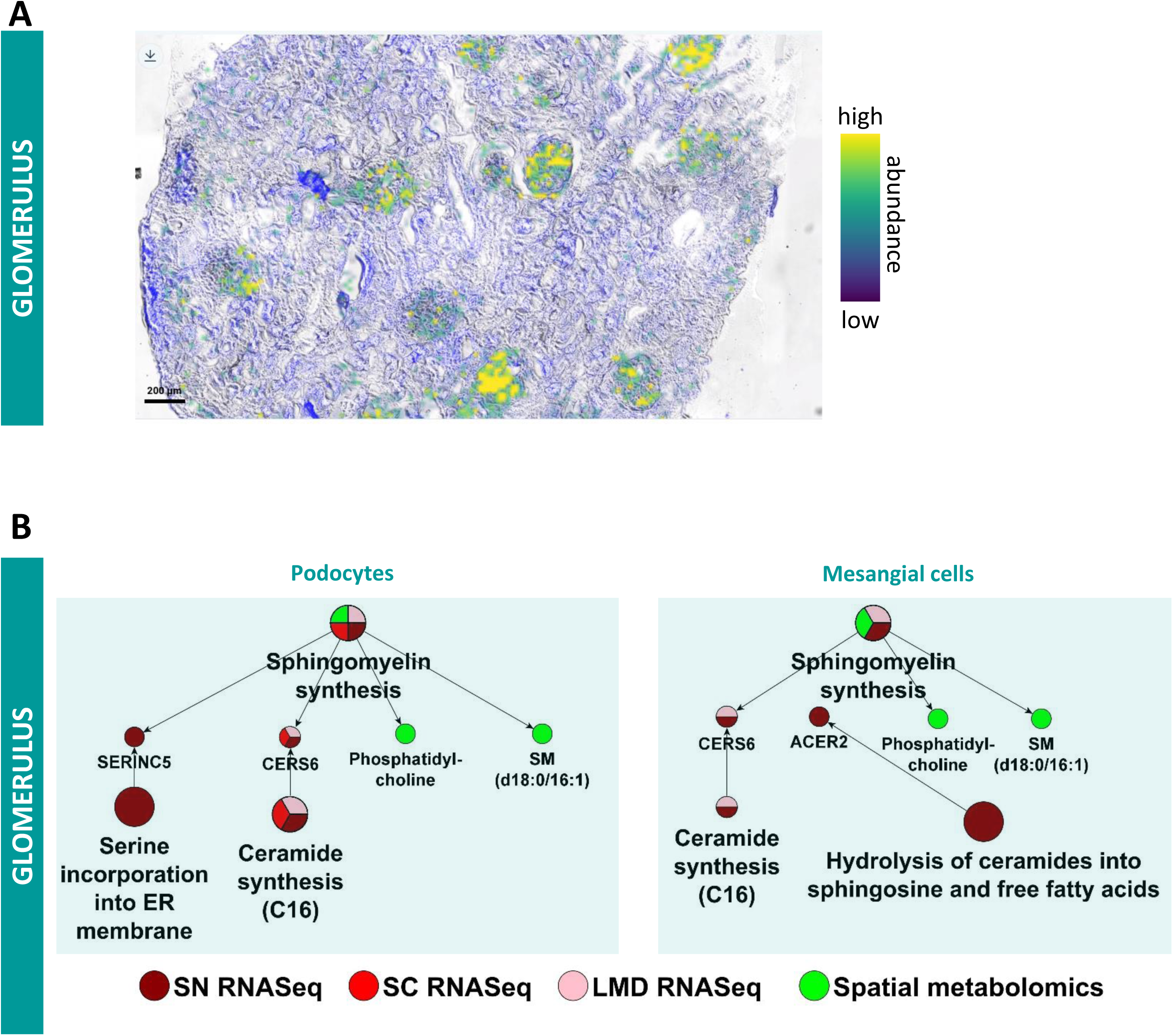
Podocytes are the synthesis site for glomerular sphingomyelin (SM) d18:0/16:1. **(A)** Matrix-assisted laser desorption/ionization mass spectrometry imaging reveals that the ion distribution of SM 18:0/16:1, [M+Na]^+^, correlates with the glomerular kidney regions. **(B)** Podocytes express two genes involved in sphingomyelin synthesis including the genes *CERS6* that is identified by both sn and sc RNAseq datasets and the LMD RNAseq dataset. CERS6 specifically generates C16 ceramides, the direct precursor for SM d18:0/16:1. **(C)** *CERS6* is also expressed in mesangial cell, though only detected by the sn RNAseq dataset. Glomerular expression of the gene *SERINC2* is detected by the LMD RNAseq assay.

### The value of a reference tissue atlas

There have been several valuable studies focused on sc transcriptomic analyses of human kidney tissue (*8-10, 13, 15, 17*) in the context of different diseases. Although each of these studies have provided substantial insight into disease processes, their mapping of undiseased kidney tissue has often been limited and focused on cell types relevant to the disease of interest. In contrast, in this study, we have studied only human kidney specimens without disease; we use multiple omics technologies, including regional and sc/sn transcriptomics, proteomics and spatial metabolomics in conjunction with imaging assays to obtain an extensive, near comprehensive spatial map of the human kidney at the single-cell resolution. Our experiments identify all known major cell types in the kidney as well as recapitulating several known subtypes. Additionally, we are able to identify different types of endothelial cells, vascular smooth muscle cells, fibroblasts and different circulating immune cells. Together, these different cell types and subtypes provide a detailed picture of the cellular and molecular composition of the human kidney. Here, we have extended our bioinformatics analyses beyond ranked lists of genes and associated pathways to identify coherent networks of pathways that give rise to function (Table 1). We have developed our model in a systematic manner such that we identify key functions for each cell type and subtypes. These physiological roles identified through pathways and marker gene lists enable the development of a multiscale atlas that connects expression patterns to whole-cell and tissue level physiological functions. The proteomics and metabolomics as well as the spatial imaging data from CODEX allows for the mapping of the sc/sn RNAseq based cell type identification to canonical cell type markers and appropriate spatial regions. This exercise provides independent orthogonal validation of both the cell type and the spatial localization within the nephron (Figs 2A and 7).

**Table 1.**
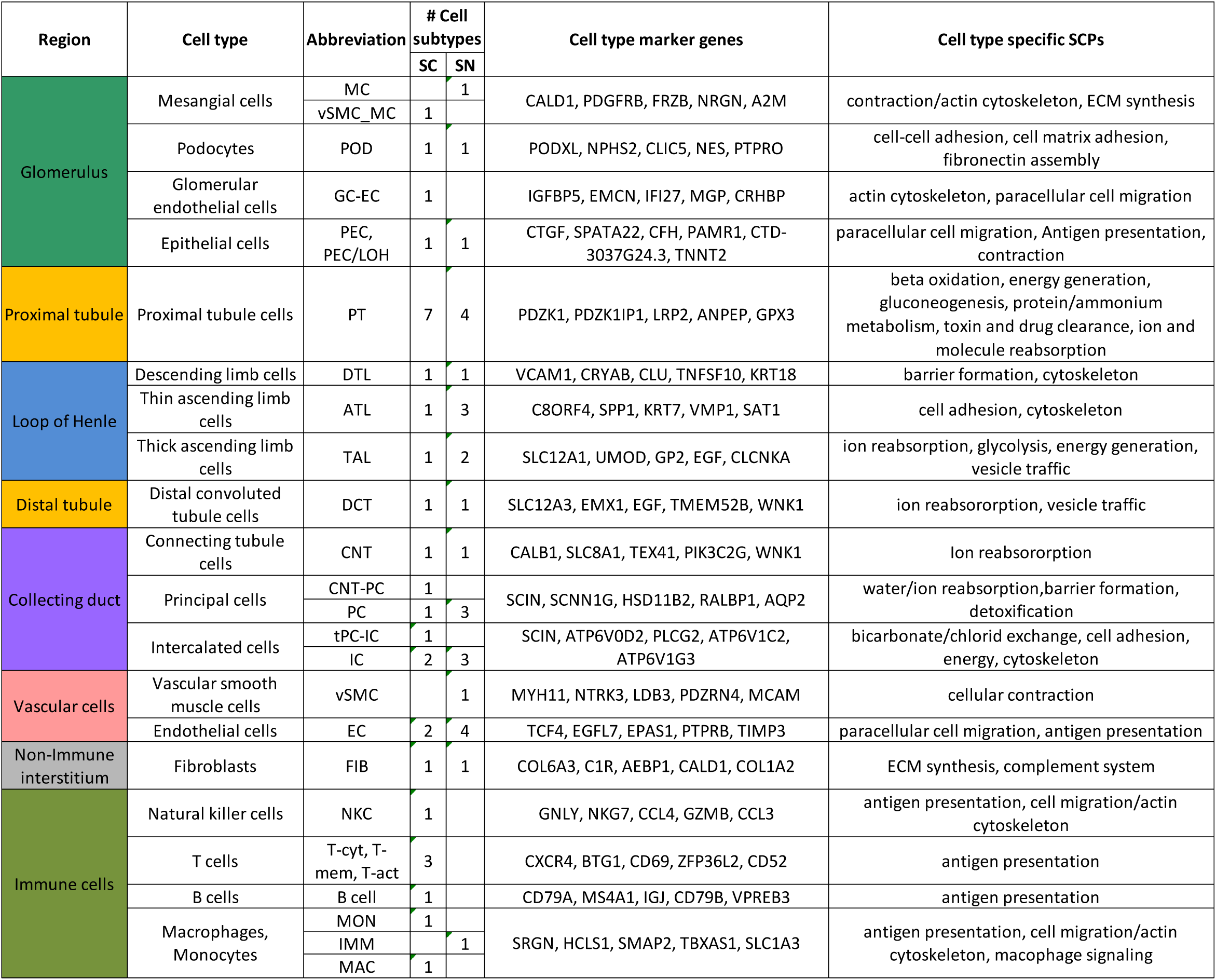
Overview of cell type specific marker genes and pathway activities.

### Limitations and future refinements

Several limitations of our study should be noted. Not all cell types are identified with the same certainty and depth, although our cell types agree well with other published studies (*9, 14, 17, 97, 113, 114*) Additionally our single cell and sn mRNA – Seq assays contain a relatively low number of cells by current standards. However, this is compensated for by use of multiple omic and other technologies, all which provide convergent conclusions in the identification of cell types.

As different cell types exist within the kidney tissue at various levels the numbers of cells for the various types of sc and sn RNAseq assays also vary widely. When fewer cells are detected, typically, we also identify a lesser number of marker genes and SCPs. Currently, we do not know if the relative number of cells we detect in the sc/sn RNAseq assays reflect the proportions *in situ*. Further experiments are needed to resolve this issue. It appears likely that to map SCPs to same depth in all cell types, additional subjects are needed. Nevertheless, for the major cell types of kidney, the *post hoc* power analyses indicate that we have sufficient power to map the core cell-level functions from SCPs. In addition, the number and functional identity of major cell subtypes need to be further studied. Currently, subtype identification is based on statistical reasoning used for the clustering algorithms. How many subtypes there are for a given kidney cell and whether these subtypes exist in all individuals requires further studies using spatial imaging technologies. Studies in other tissues, such as brain (*115*) and heart (*116*), have identified multiple subtypes of paraventricular interneurons and ventricular myocytes. Hence, it is likely that kidney cell types may also contain major subtypes. In spite of these limitations, this study provides a detailed functional view of a kidney map at single-cell resolution which can be used to understand major aspects of kidney physiology as demonstrated by the two examples described above.

Information becomes knowledge only when it is deliberately and systematically cataloged such that new cohesive insights can readily be drawn, as shown above for sphingomyelin related functions in podocytes. Ontology is an ideal tool that can logically represent the data and metadata in a human- and computer-interpretable manner. It can enable the generation of new knowledge, especially when such knowledge involves multiscale relationships between molecules, cell types and their subtypes and tissue level physiological function. In addition to the integrated analytics presented here, KPMP is also building a community-based Kidney Tissue Atlas Ontology (KTAO) (*117*). KTAO will systematically integrate different types of information (such as clinical, pathological, cell and molecular) into a logically defined tissue atlas, which can then be further utilized to support various applications. Taken together, the final knowledge environment and the kidney tissue atlas constructed by KPMP, which is available at www.atlas.kpmp.org, should be able to help molecularly characterize cellular types and subtypes in the kidney; improve patient care by providing new disease classifications; and may ultimately lead to new patient-specific novel therapeutic approaches.

## Supporting information

Supplementary Table 1 - Metadata

Supplementary Table 2 - LMD RNAseq

Supplementary Table 3 - LMD Proteomics

Supplementary Table 4 - NSC Proteomics

Supplementary Table 5 - Marker genes and proteins

Supplementary Table 6 - No of significant marker genes and proteins

Supplementary Table 7 - MBCO enrichment results

Supplementary Table 8A - Spatial Metabolomics 18-139

Supplementary Table 8B - Spatial Metabolomics 18-142

Supplementary Table 8C - Spatial Metabolomics 18-342

Supplementary Table 9 - humanbase

Supplementary Table 10 - Essential genes

Supplementary Table 12 - Curated sodium transport mechanisms

Supplementary Table 11 - Metabolism enrichment results

## Data Availability

All raw and processed data described in this manuscript is available through the KPMP Data Portal at kpmp.org.

## Acknowledgements

Kidney Precision Medicine Project acknowledges all the participants, patients, and the scientific officers from National Institute of Diabetes and Digestive and Kidney Diseases. KPMP was supported by NIH grants UH3 DK114923, UH3 DK114920, UH3 DK114933, UH3 DK114937, UH3 DK114907 and U2C DK114886. A complete list of all KPMP members can be found at kpmp.org. We thank Joseph Goldfarb for critically reading of the manuscript.

## Author Contributions

**Integrated analysis and interpretation:** Jens Hansen, Rachel Sealfon, Rajasree Menon, John Cijiang He, Jonathan Himmelfarb, Lisa Satlin, Olga G. Troyanskaya, Matthias Kretzler, Ravi Iyengar, Evren U. Azeloglu

**Pilot tissue procurement, data coordination and metadata curation:** Becky Steck, Abhijit Naik, Jeffrey B. Hodgin, Matthias Kretzler, Evren U. Azeloglu

**SC/SN RNASeq data generation and processing:** Rajasree Menon, Blue B. Lake, Jens Hansen, Edgar A. Otto, Jeffrey B. Hodgin, Laura Barisoni, Minnie Sarwal, Kun Zhang, M. Todd Valerius, Sanjay Jain Matthias Kretzler, Evren U. Azeloglu

**LMD transcriptomics data generation and processing:** Michael T. Eadon, Daria Barwinska, Pierre C. Dagher

**LMD and NSC proteomic data generation and processing:** Samir Parikh, John P. Shapiro, Tara K. Sidgel, Priyanka Rashmi, Minnie Sarwal, Brad Rovin

**Imaging data generation and processing:** Dejan Dobi, Seth Winfree, Tarek M. El-Achkar, Zoltan Laszik

**Spatial metabolomics data generation and processing:** Theodore Alexandrov, Dusan Velickovic, Christopher R. Anderton, Guanshi Zheng, Annapurna Pamreddy, Kumar Sharma

**Manuscript preparation:** Jens Hansen, Rachel Sealfon, Rajasree Menon, Michael P. Rose, Yongqun He, Ian H. de Boer, Lisa Satlin, Olga G. Troyanskaya, Matthias Kretzler, Ravi Iyengar, Evren U. Azeloglu. All authors commented and edited the manuscript and assisted in the assembly of the final version.

---

**Supplementary Figure 1:**
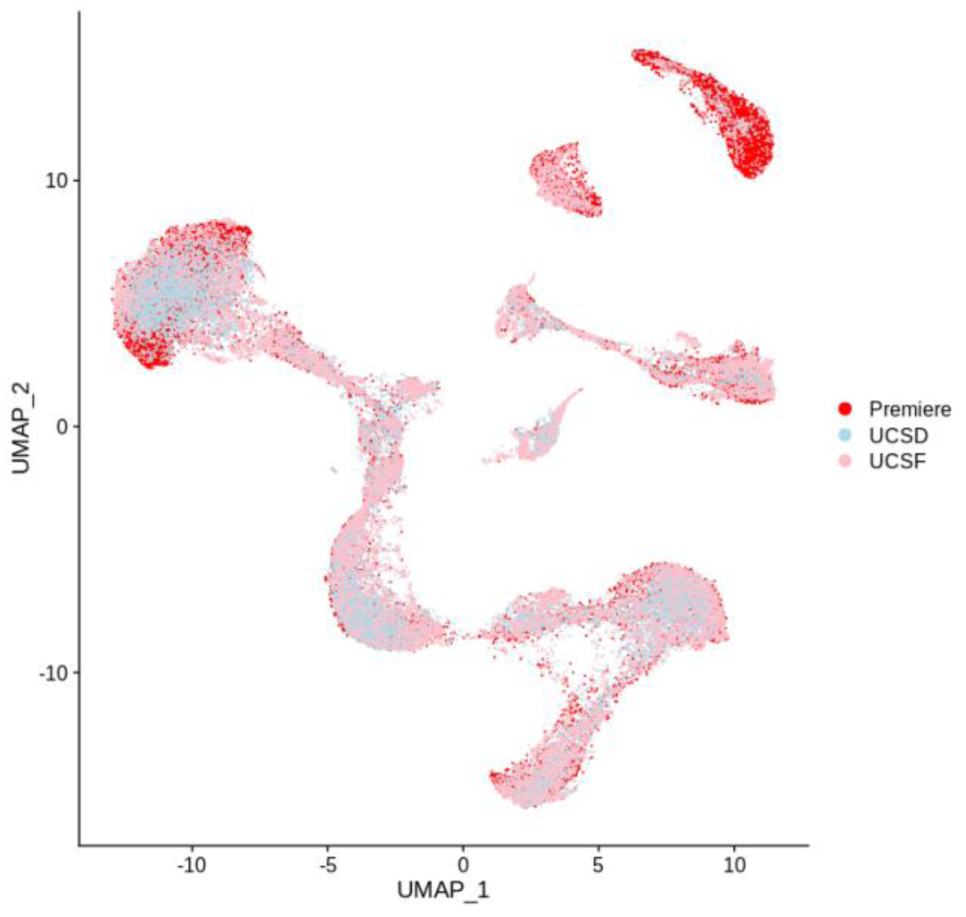
Coloring of cells and nuclei by dataset documents that each cell cluster contains cells and nuclei from each dataset.

**Supplementary Figure 2:**
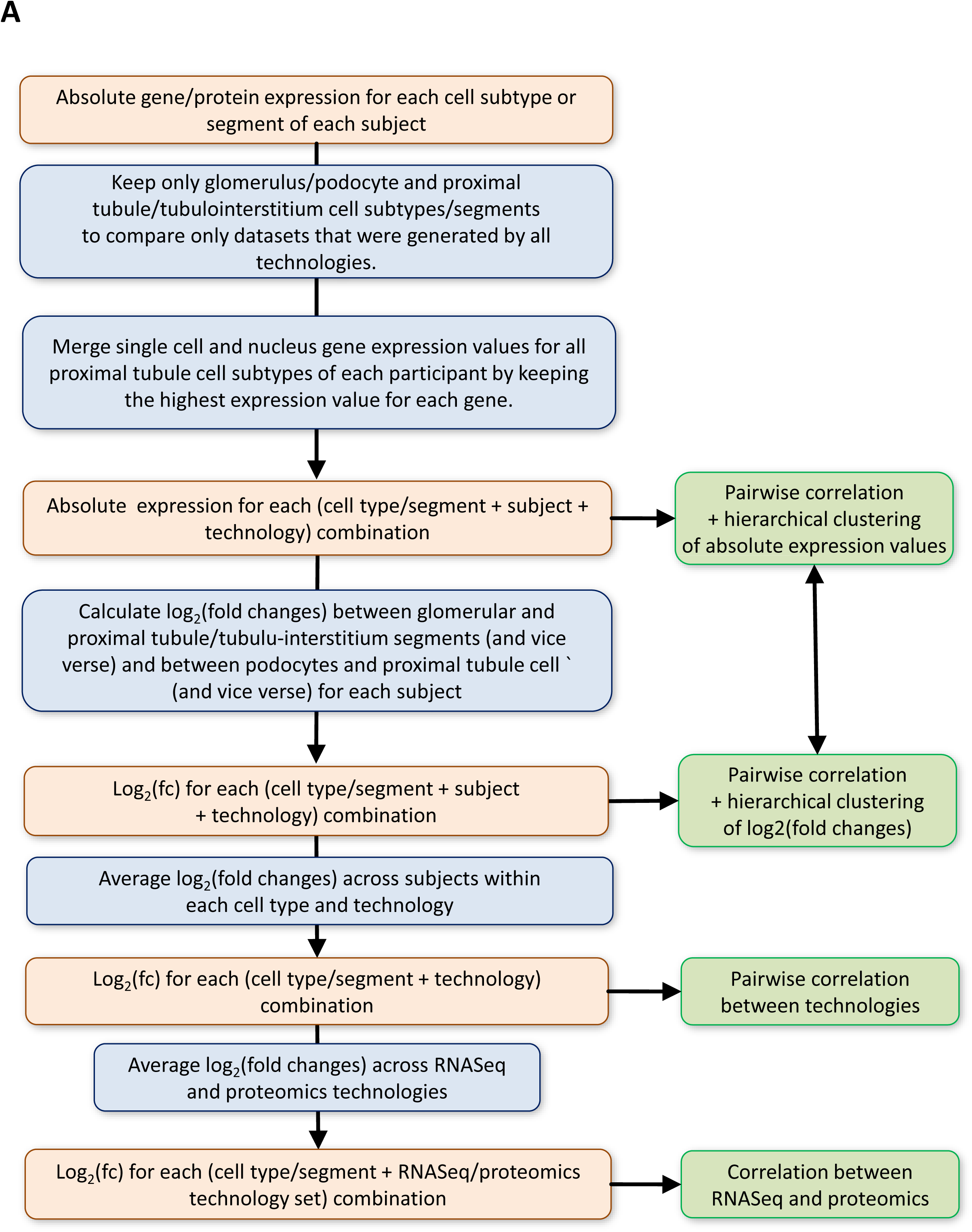
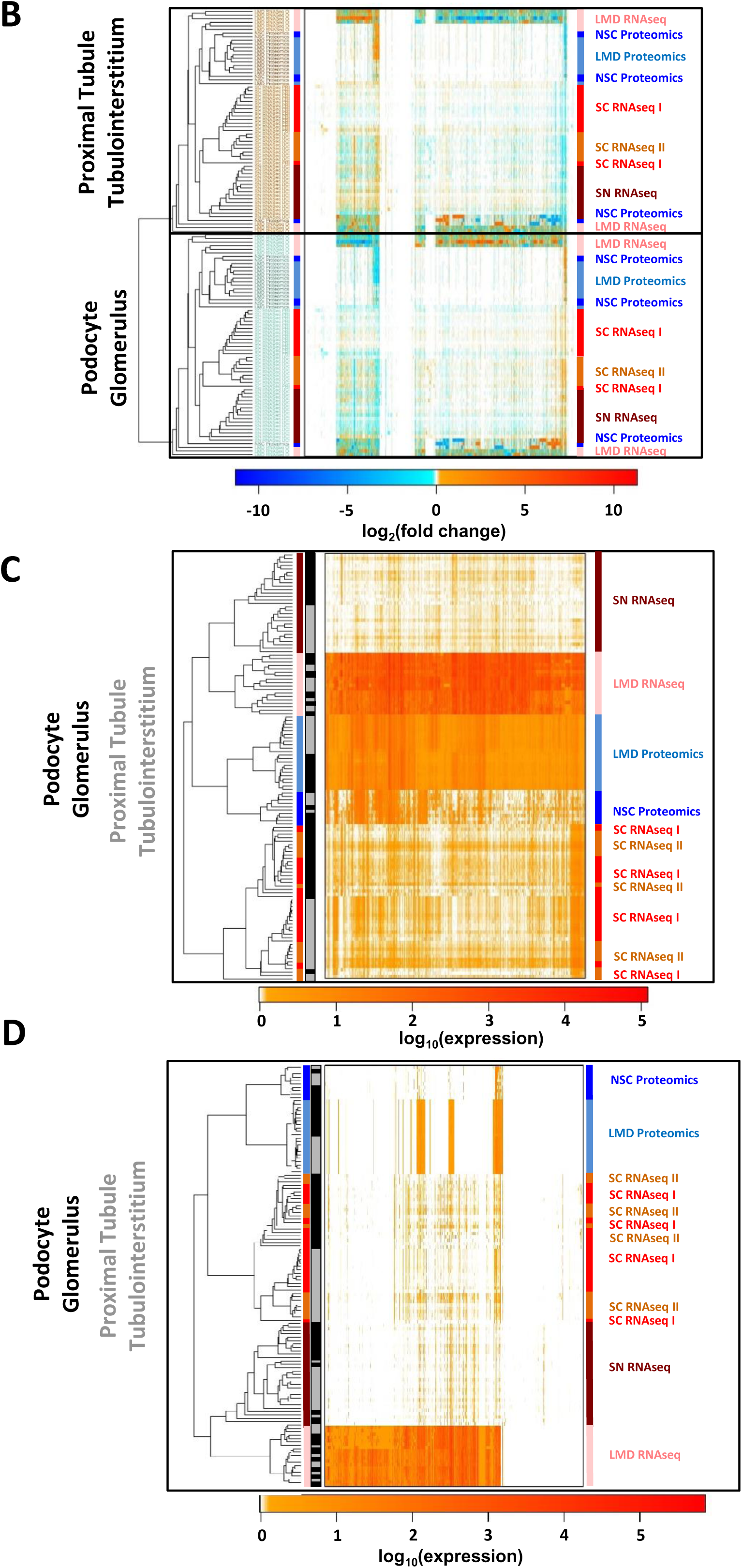
Cross-platform comparison of gene and protein expression. **(A)** Pipeline for correlation analysis across different omics technologies. See methods for details. **(B)** Hierarchical clustering of pairwise correlation coefficients between all samples based on the log_2_(fold changes) without removal of those genes and proteins that are not consistently detected across all assays also groups the samples by anatomical region and not technology. In contrast, pairwise correlation and hierarchical clustering based on logarithmized absolute expression values groups samples by technology, **(C)** with or **(D)** without removal of the not consistently detected genes and proteins.

**Supplementary Figure 3:**
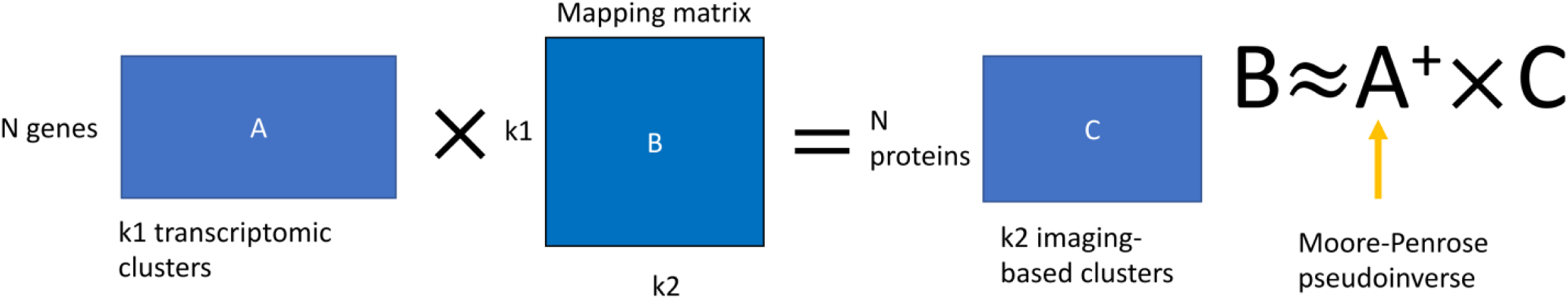
Illustration of method used for mapping of single cells/nuclei to CODEX.

**Supplementary Figure 4:**
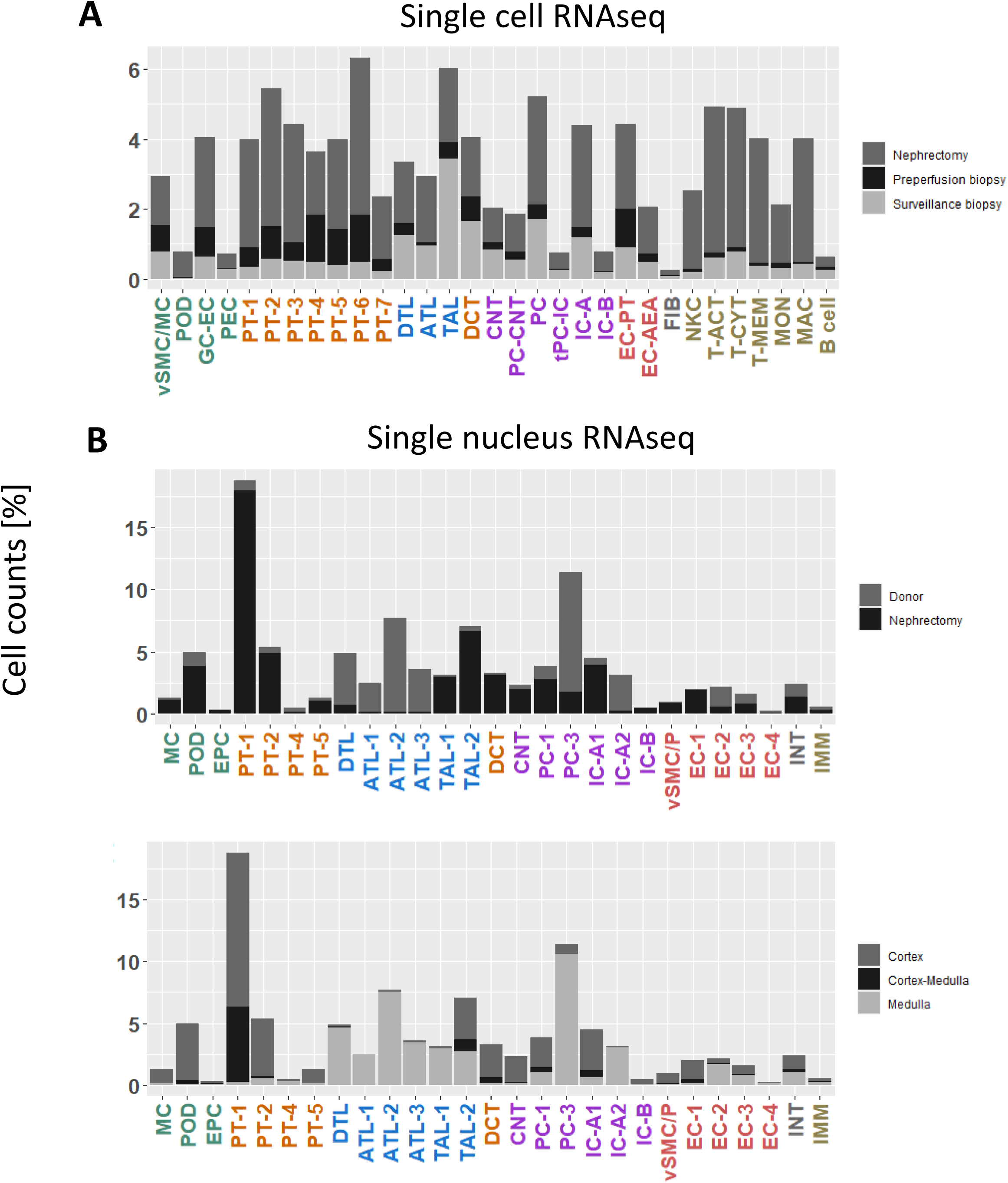
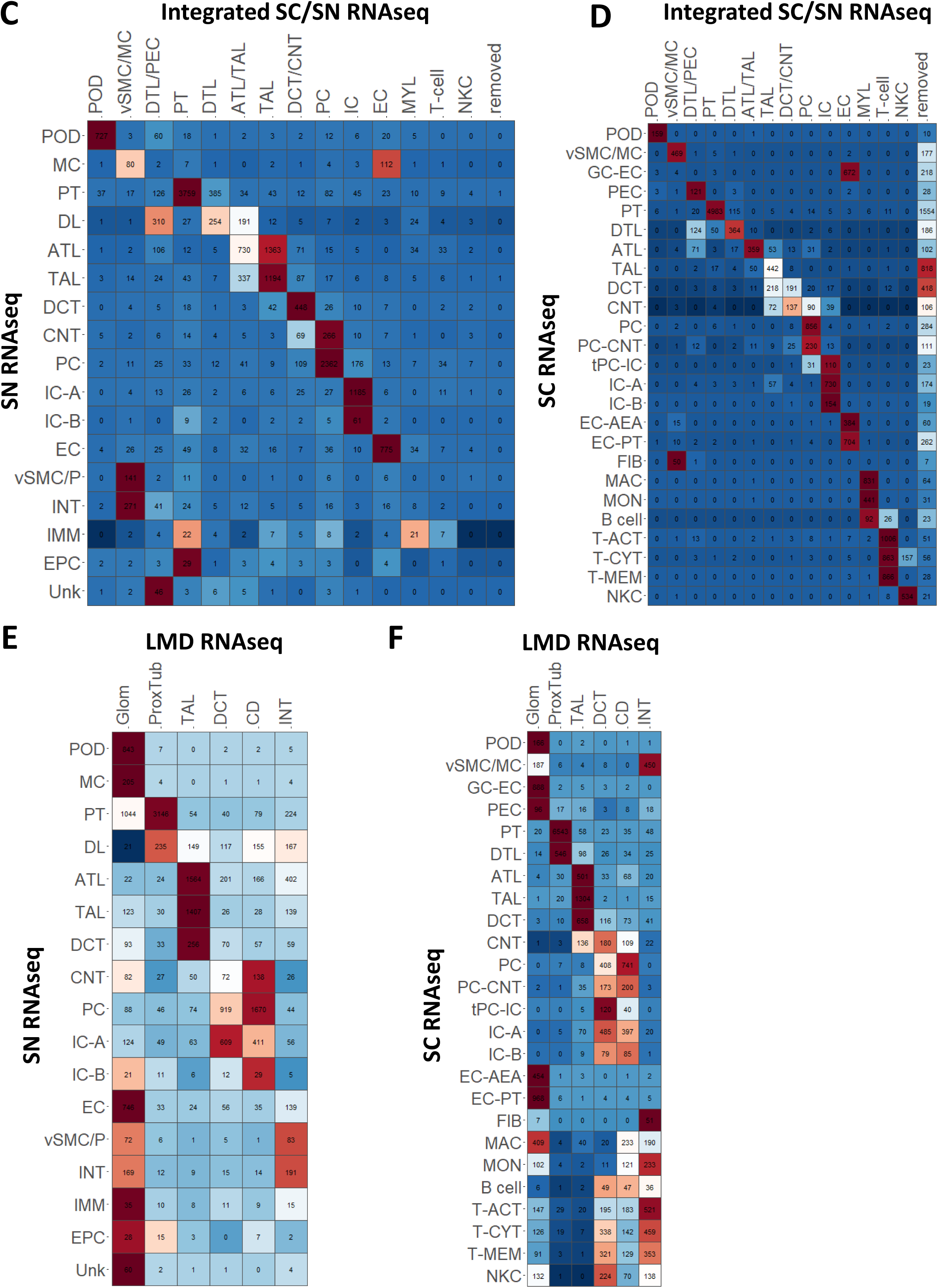
Separated and integrated analysis of sc and sn RNAseq datasets generate consistent cell type mapping. Cell types and subtypes identified by the separated analyses of the **(A)** sn and **(B)** sc RNAseq datasets. Bars indicate the percentage of all cells that mapped to a particular cell type or subtype, colors indicate the tissue collection method each particular cell was obtained by. Cell type assignments of separate clusters from **(C)** sn and **(D)** sc RNAseq datasets were compared to those obtained by the integrated analysis. Numbers indicate nuclei/cell counts; fields are colored by the percentage of cells within each field compared to the row margins. Note that in separated analyses of the sc RNAseq dataset, the applied cutoff for mitochondrial gene expression was higher (≤50% instead of ≤20%); consequently, some of the cells that were removed in the combined analysis were assigned to cell types in the separated analysis. Similarly, mapping of the **(E)** nuclei and **(F)** cells to LMD segments documents that the annotations obtained from the separated analyses map to their correct anatomical origin, as observed for the integrated analysis. All heatmaps are colored according to the number of cells assigned to each LMD subsegment, scaled so each row has mean of 0 and standard deviation of 1. See figure 2A for cell type abbreviations.

**Supplementary Figure 5.**
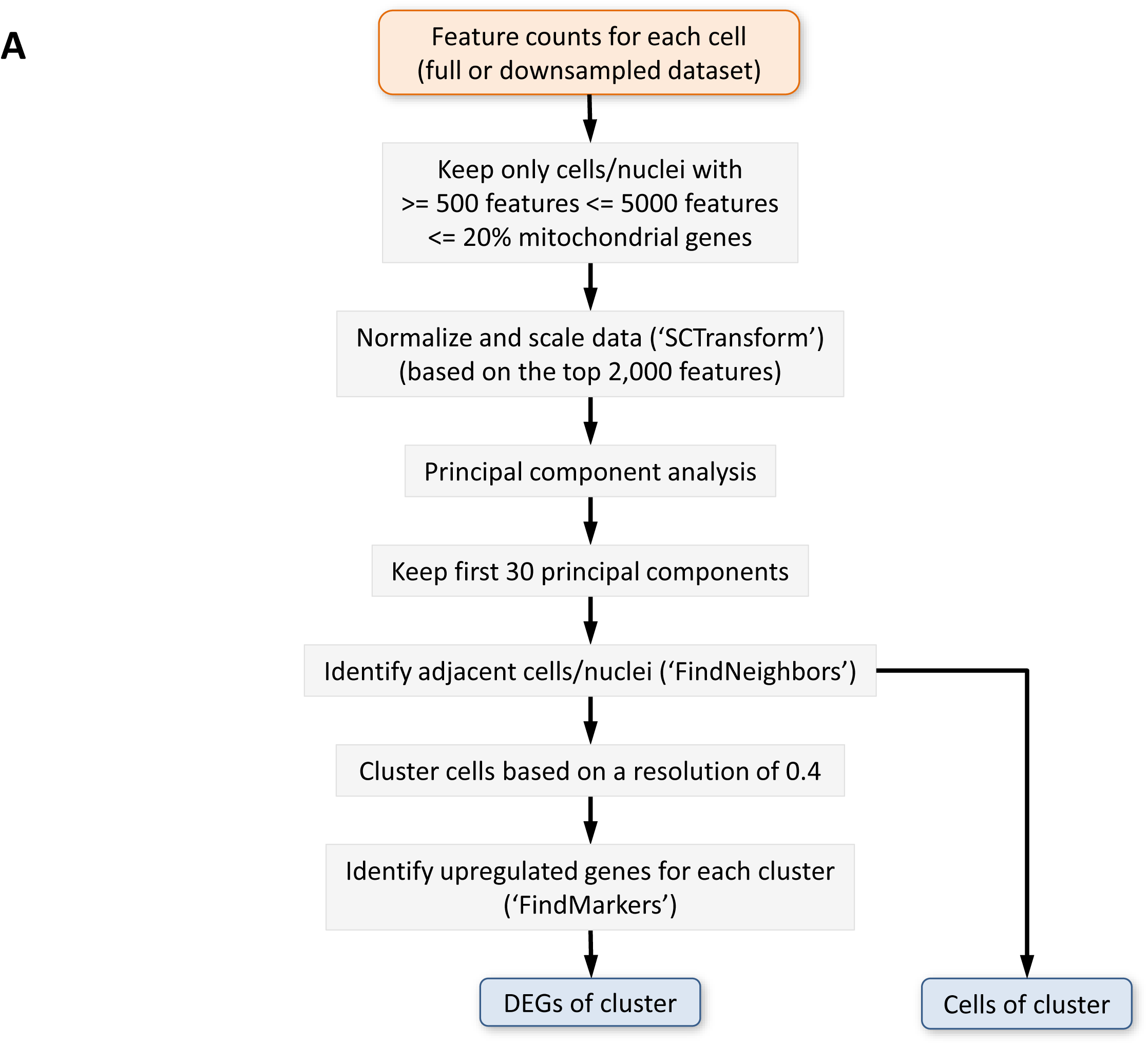
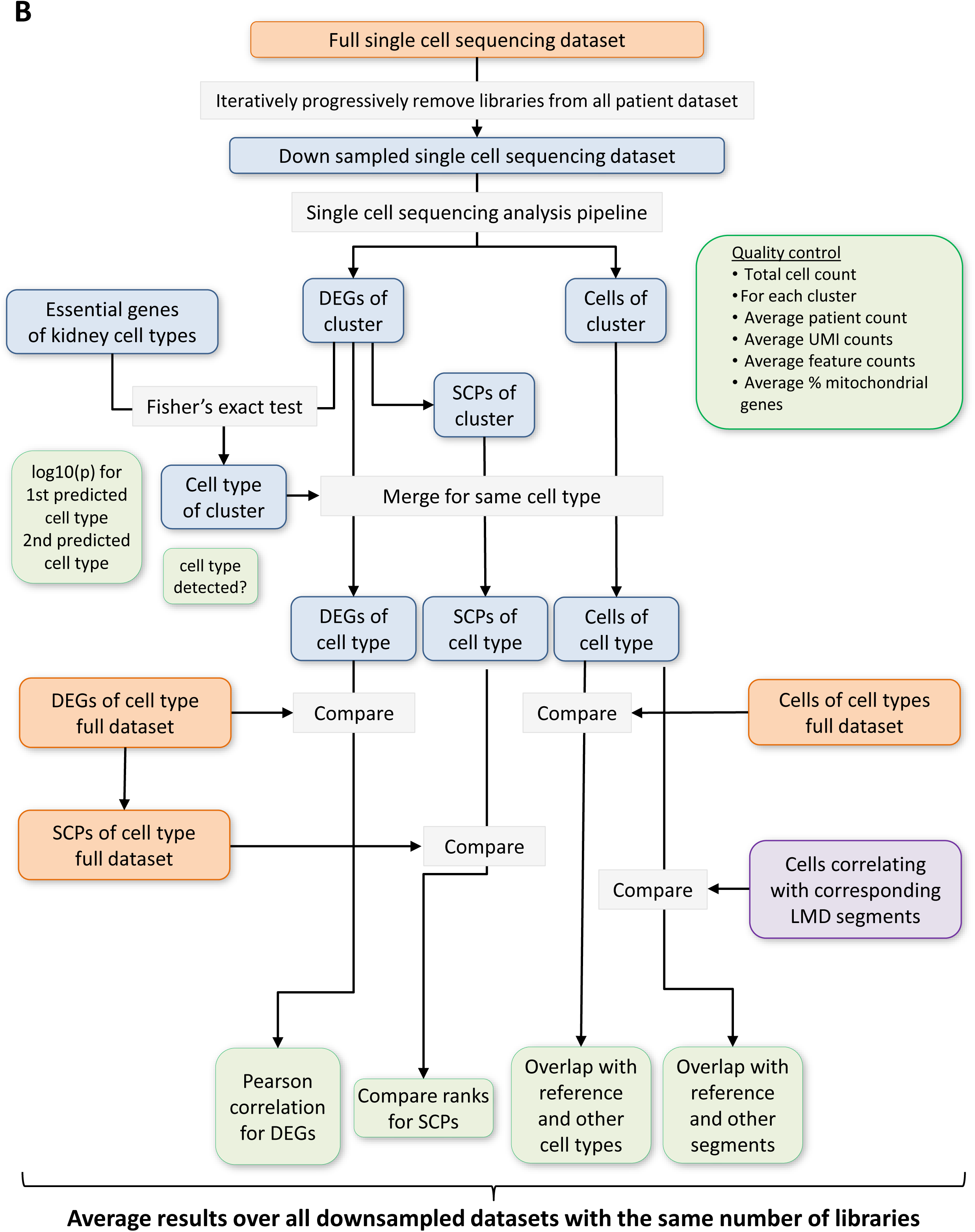
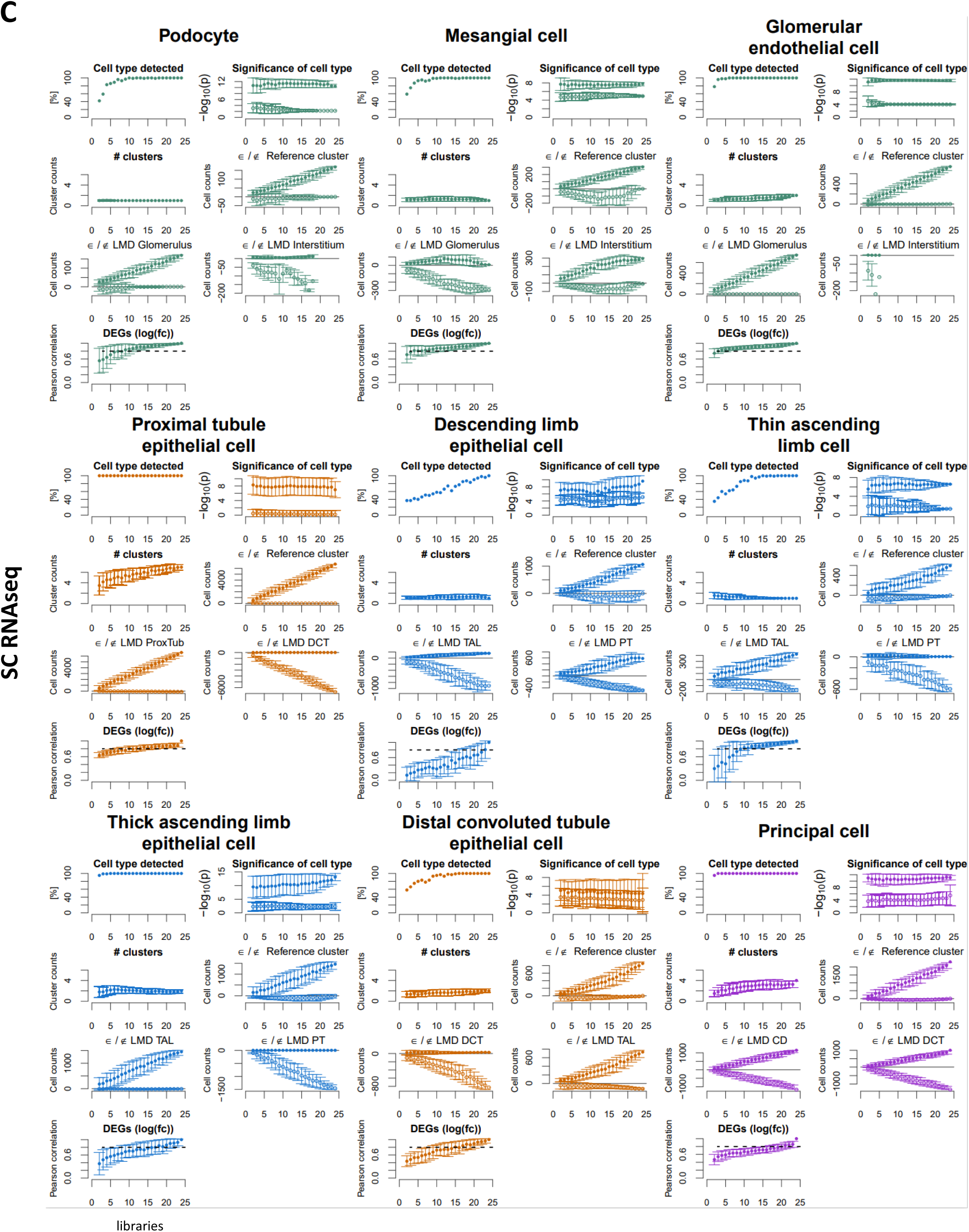
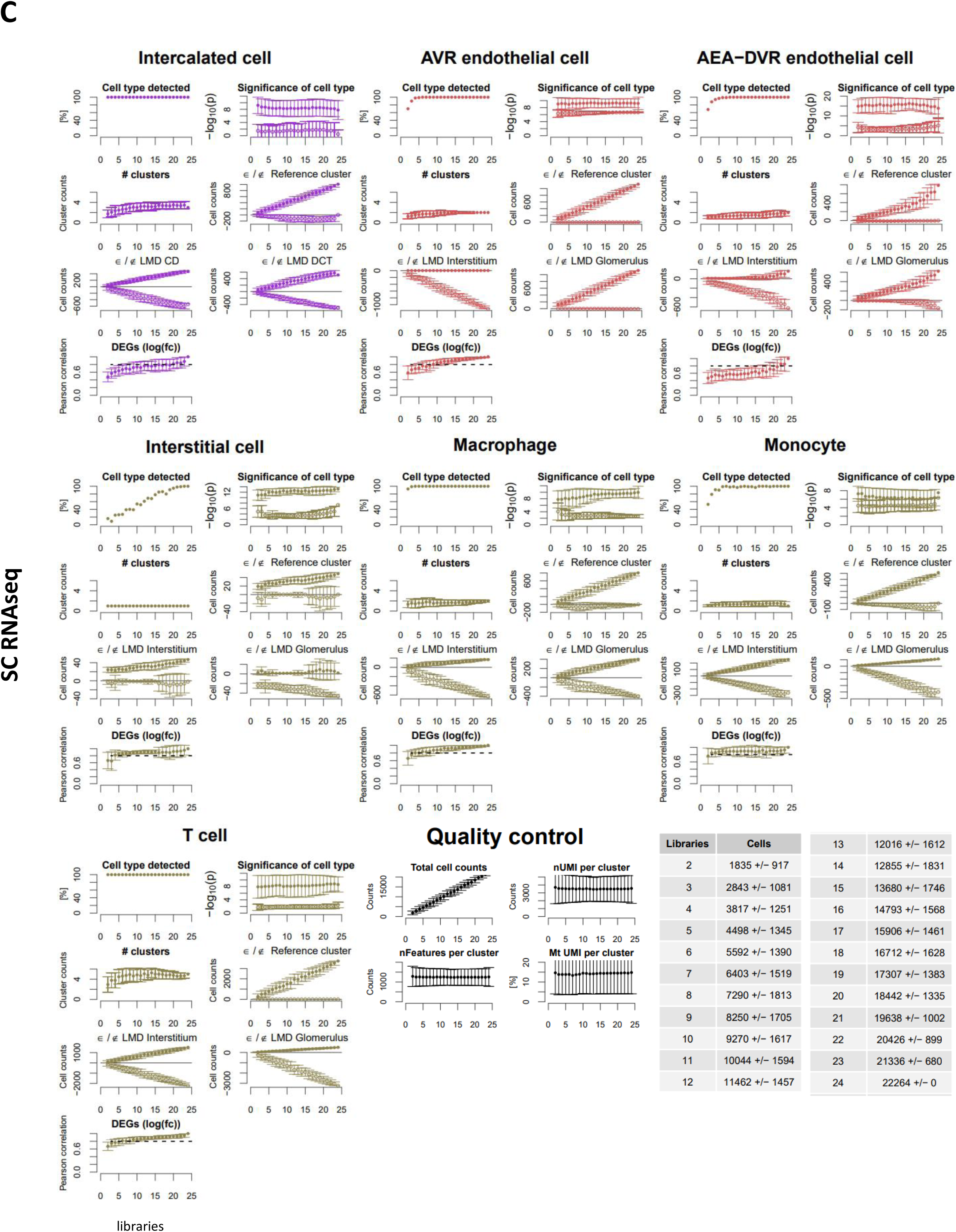
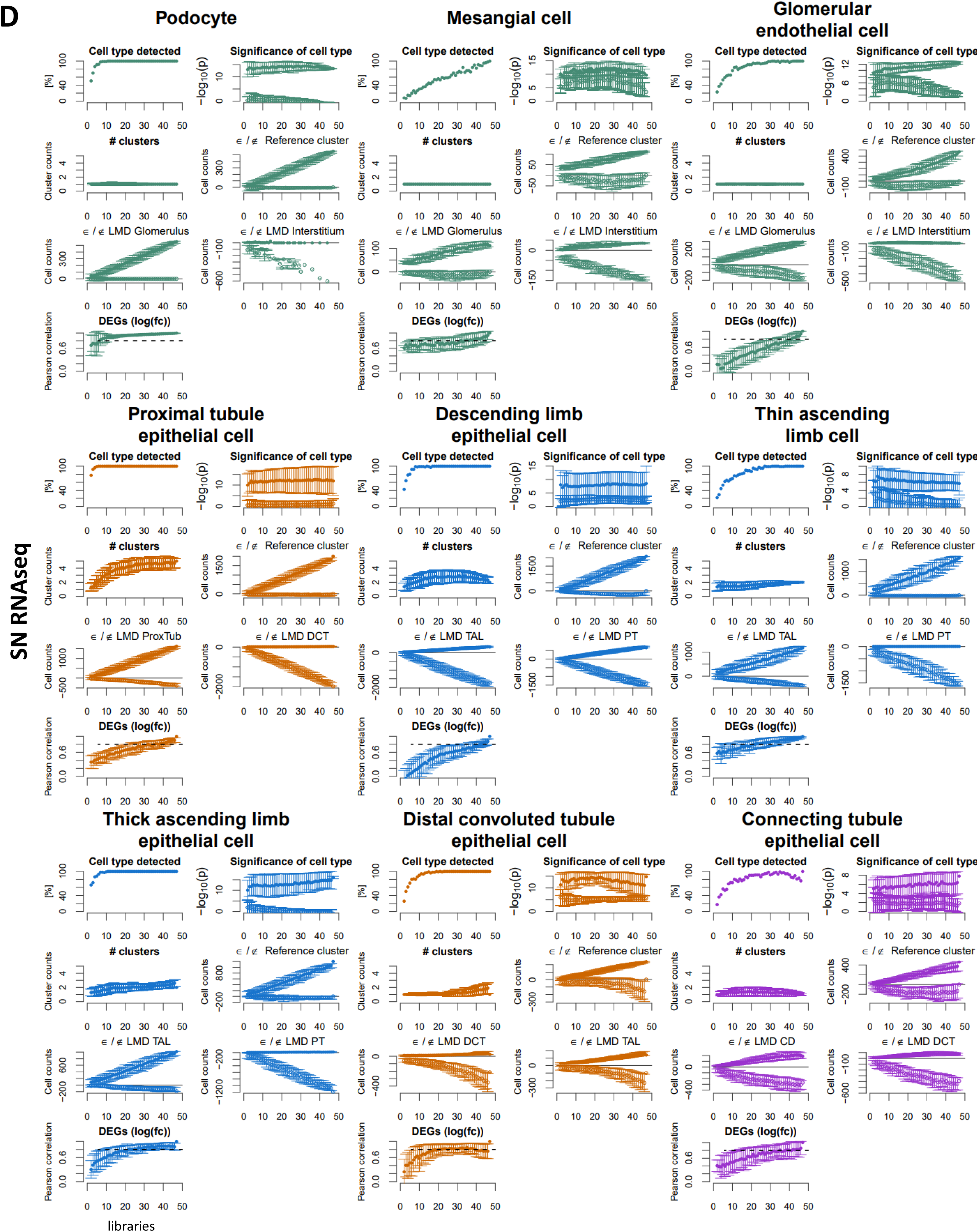
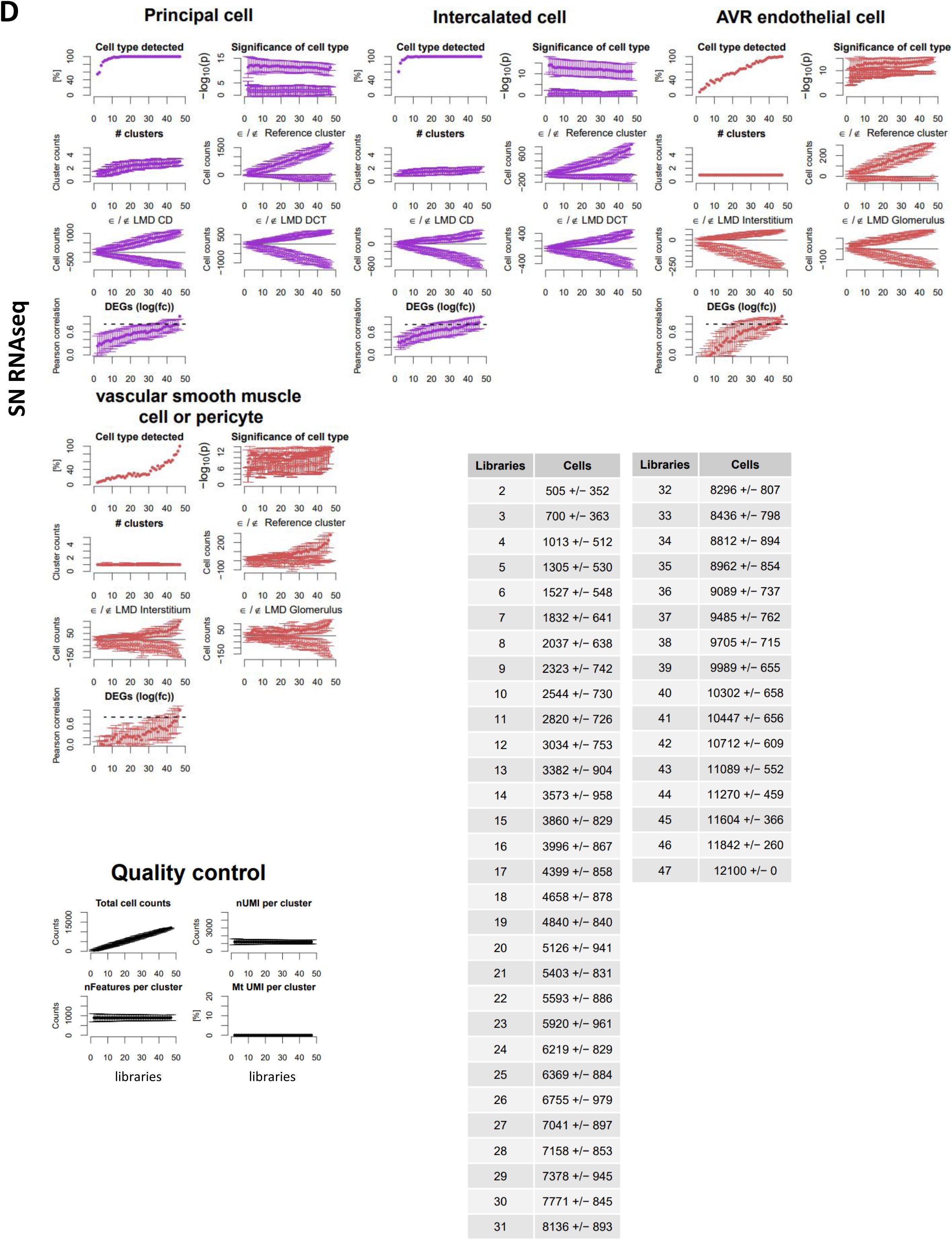
Complete results of single-cell/nucleus transcriptomic *post hoc* power analysis. Subject libraries or samples were randomly and progressively removed from the sc (24 libraries) and sn (47 libraries) RNAseq to generate at max 100 non-overlapping random groups for each number of remaining libraries. **(A)** Sc and sn datasets were subjected to an automated sc/sn data analysis pipeline. **(B)** Results were averaged for each number of subject libraries and compared between the downsampled and full datasets as indicated. *Post hoc* power results of the **(C)** sc and **(D)** sn RNAseq datasets. **‘Cell type detected’:** This plot documents how often (in percent) a particular cell type was detected in dependence of the number of analyzed libraries. **‘Significance of cell type’**: To assign cell types to each cluster we subjected cluster specific marker genes to enrichment analysis using Fisher’s Exact test and a list of literature curated cell-type specific essential genes. For each cluster predicted cell types were ranked by significance and the top ranked cell type was assigned to that cluster. The plot shows the -log_10_(p-values) of the first (i.e. the selected) and the second ranked cell type. Comparison of both p-values allows an estimation of the reliability of a particular cell type assignment. The larger the difference between both -log_10_(p-values), the more certain is that particular cell type assignment. **‘# clusters’** documents how many clusters were assigned to that particular cell type. **‘element/not element of Reference cluster’:** Cells/nuclei that were assigned to the same (above abscise, positive values, full circles) or to a different cell type (below abscise, negative values, open circles) as in the full dataset were counted in each downsampled dataset. **‘element/not element of indicated LMD subsegment’:** Using cell and nuclei mappings presented in Suppl. Figure 4E/F we counted how many cells/nuclei of a particular cell type mapped to the indicated LMD subsegment (above abscise, positive values, full circles) or to a different LMD subsegment (below abscise, negative values, open circles). **‘DEGs log(fc)’**: Correlation between the log fold changes of cell type specific markers obtained for the downsampled and full dataset. Notify that all comparisons were only done, if a particular cell type was detected (as indicated in the first diagram). See figure 2A for cell type abbreviations.

**Supplementary Figure 6:**
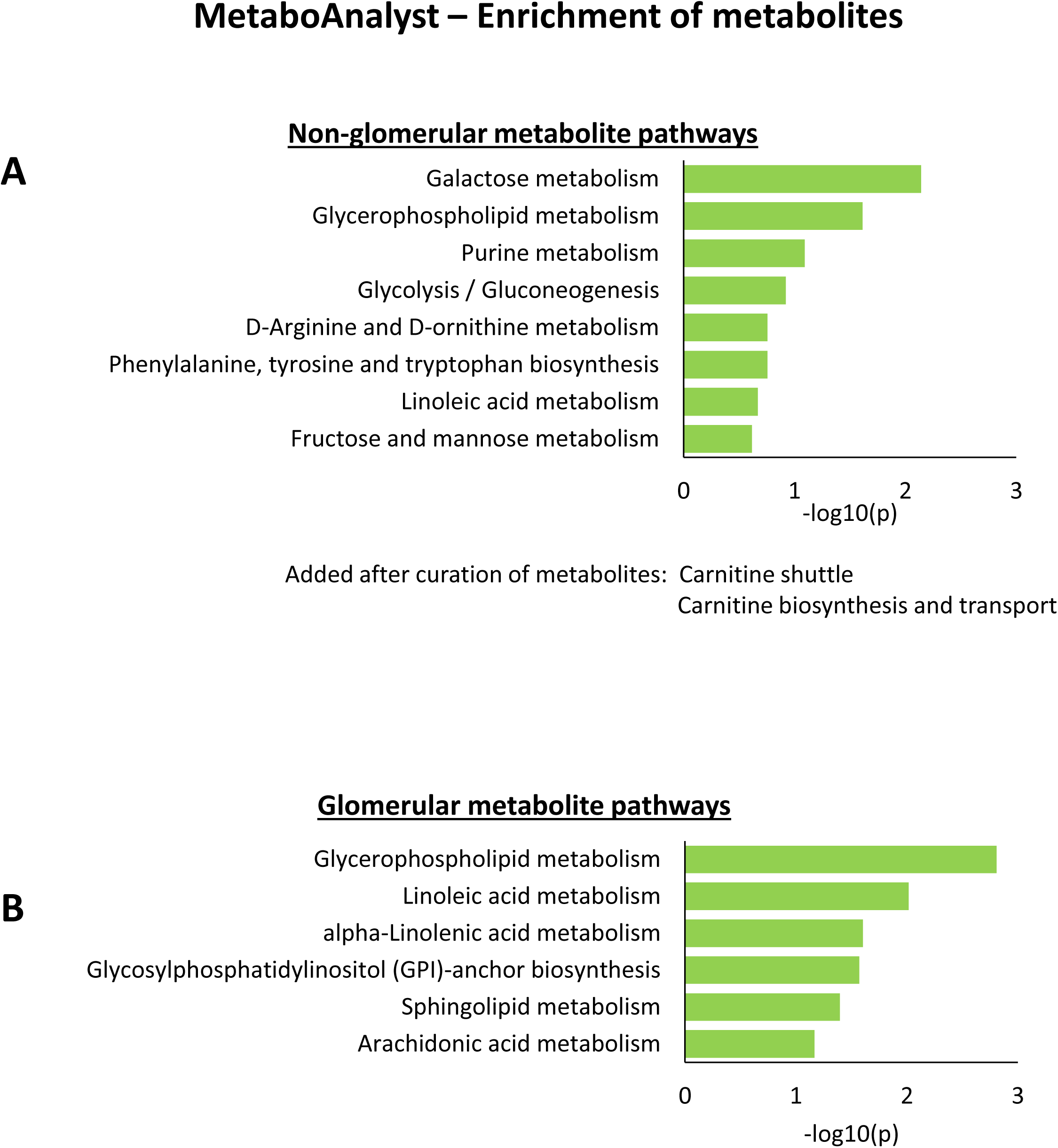
Pathway enrichment analysis of spatial metabolomics data. All **(A)** Non-glomerular and **(B)** glomerular metabolites obtained from the three nephrectomy samples were subjected to pathway enrichment analysis using MetaboAnalyst. Some pathways were predicted from metabolites that are general precursors for the synthesis of multiple products and participate in multiple pathways. To exclude such unspecific and consequently uncertain pathway predictions, we focused only on those pathways that were predicted from a pathway specific metabolite (see methods for details). To merge the metabolic pathways with the MBCO SCP-networks, we mapped the MetaboAnalyst pathways ‘Glycolysis/Gluconeogenesis’ and ‘Glycerophospholipid metabolism’ to the MBCP SCPs ‘Glycolysis and Gluconeogenesis’ and to ‘Phosphoglyceride biosynthesis’, respectively. Based on identified metabolites, we added the MBCO SCPs “Carnitine shuttle” and “Carnitine biosynthesis and transport” to the predicted MetaboAnalyst pathways (see methods for details).

**Supplementary Figure 7:**
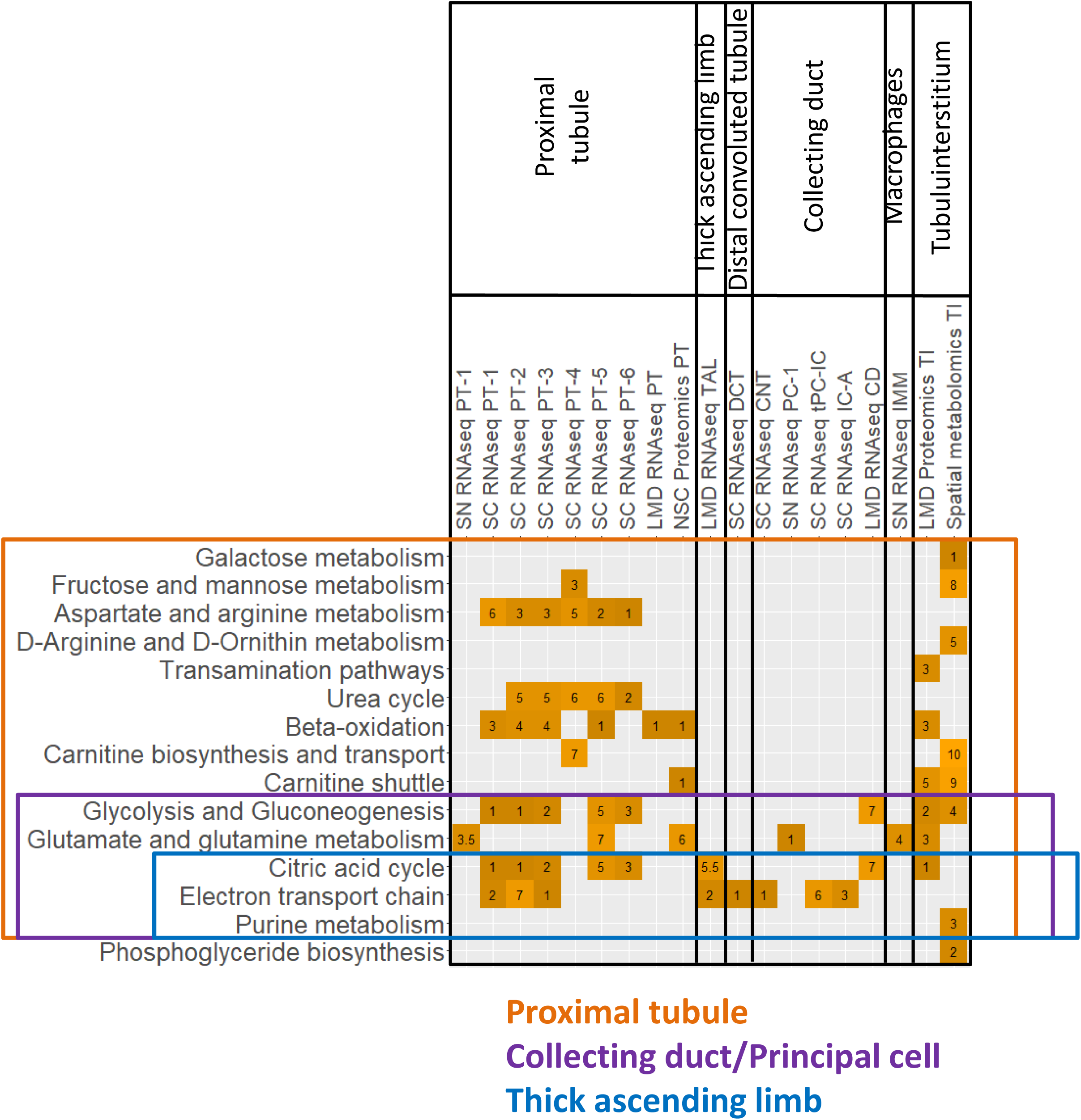
Mapping of tubulointerstitial SCPs to cell types. SCPs predicted by dynamic enrichment analysis for the tubulointerstitial segment by the LMD Proteomics and spatial metabolomics assays were mapped to one of three detected glomerular cell types, because they were either detected in that cell type as well or related to SCPs detected for that cell type. Numbers indicate at which rank a particular SCP was detected. Notify that dynamic enrichment analysis can predict single SCPs or combinations of up to three SCPs, and consequently the same rank can be given to multiple SCPs. When an SCP was predicted by multiple cell subtypes, the highest rank is visualized in this figure.

**Supplementary Figure 8:**
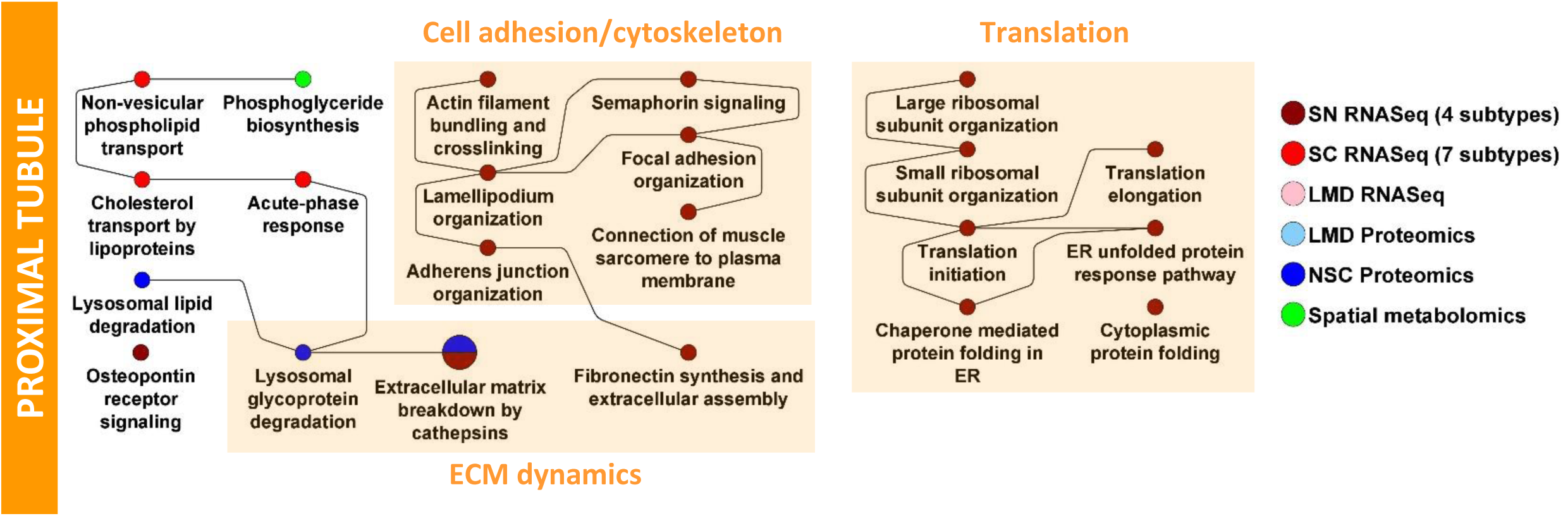
Enrichment analysis of differentially expressed genes and proteins in proximal tubule cells and subsegments. See Figure 5 for details. SCPs that were among the top seven predictions based on dynamic enrichment analysis of PT DEGs and DEPs and were removed from the main figure for space reasons.

**Supplementary Figure 9:**
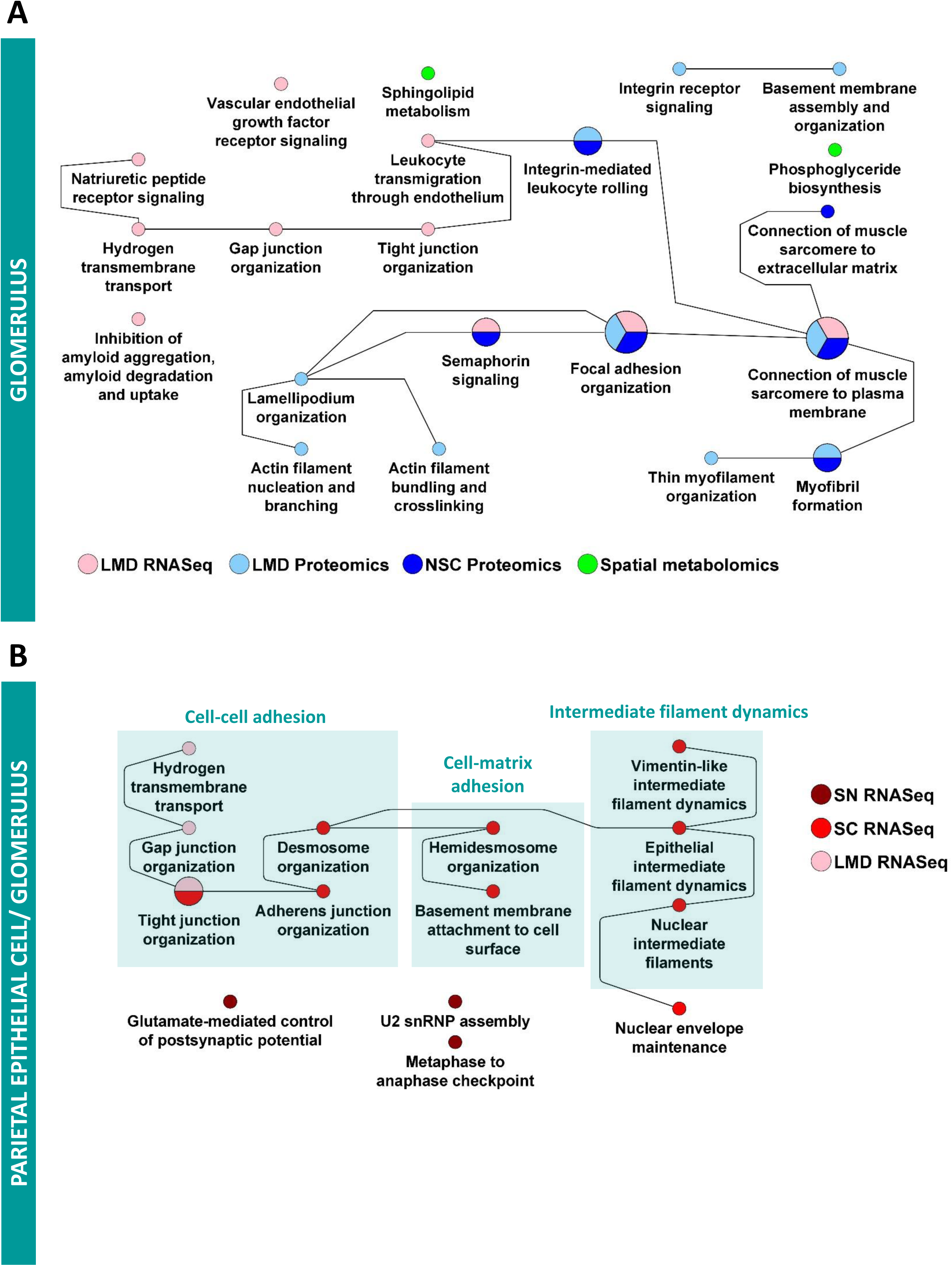
Enrichment analysis for glomerular datasets. **(A)** DEGs and DEPs identified by LMD RNAseq and Proteomics and NSC Proteomics were subjected to dynamic enrichment analysis. **(B)** SCP network predicted for parietal epithelial cells by the sn and sc RNAseq datasets were merged with those SCPs that were predicted by the glomerular segment-specific datasets and assigned to this cell type (Figure 5C).

**Supplementary Figure 10:**
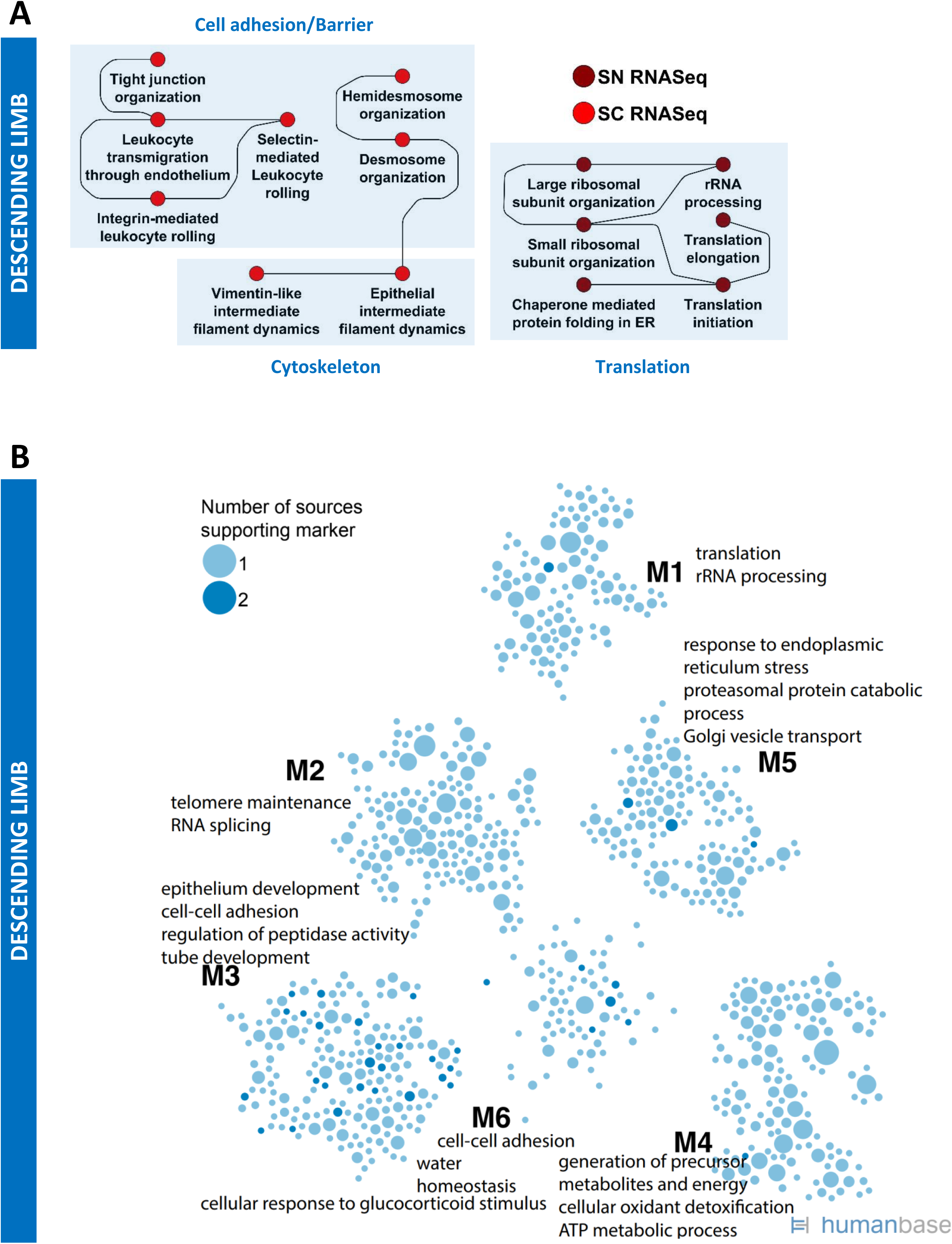
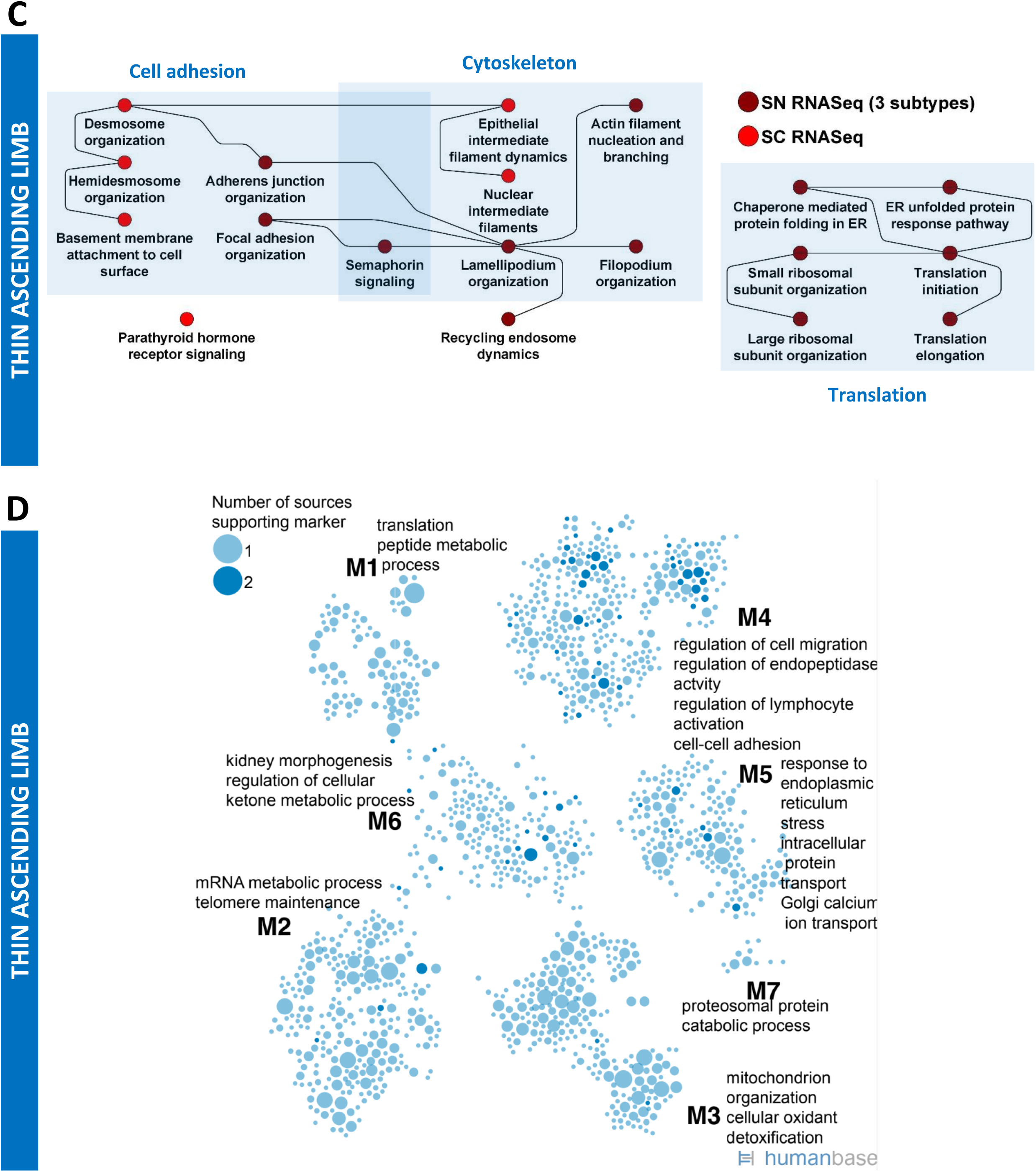
Enrichment analysis for the Loop of Henle. **(A/B)** Descending limb cell specific DEGs were subjected to dynamic enrichment (A) and module analysis (B). **(C/D)** Similarly, thin ascending limb cell specific DEGs were subjected to dynamic enrichment (A) and module analysis (B).

**Supplementary Figure 11:**
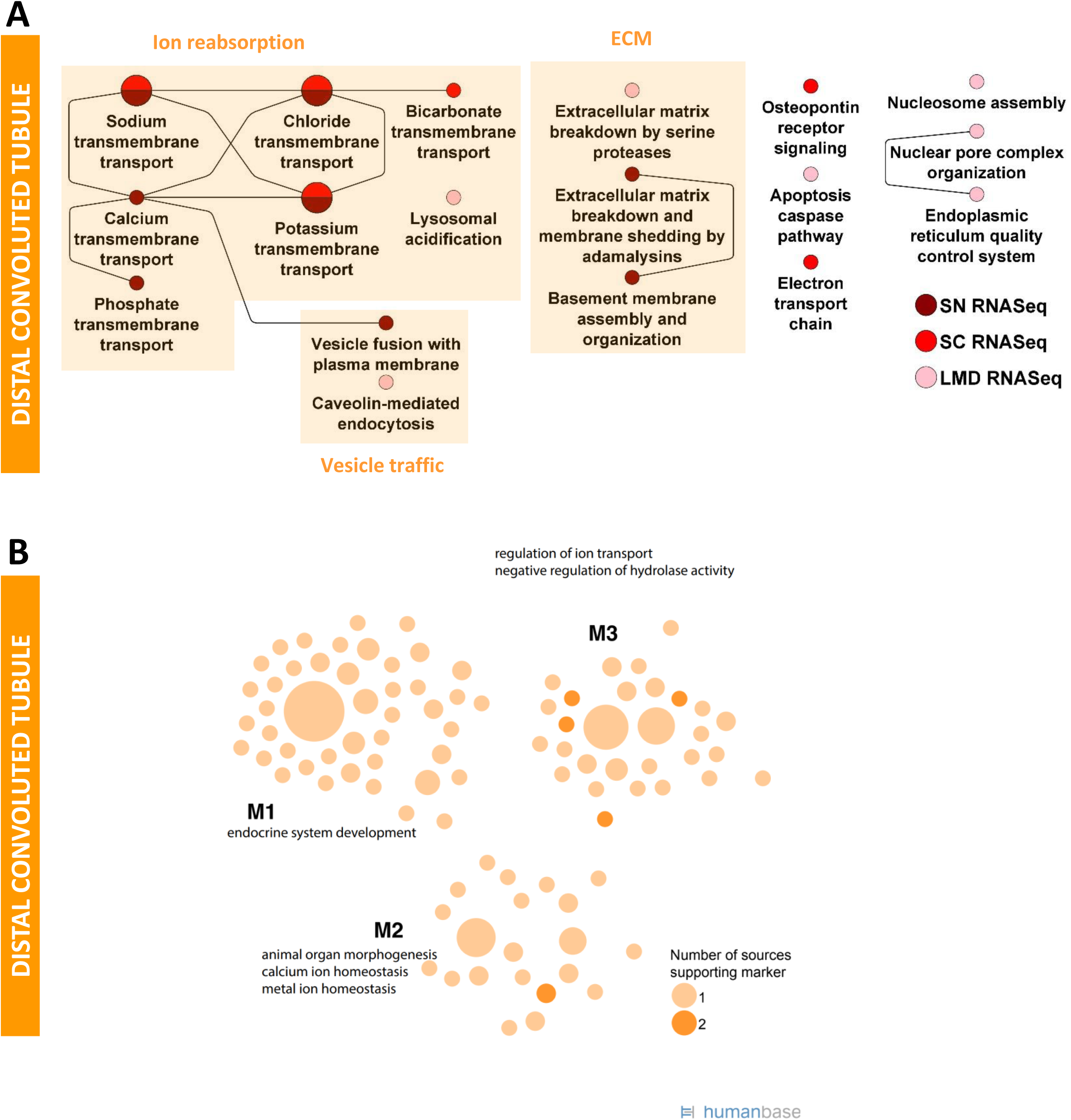
Enrichment analysis for the distal convoluted tubule. **(A/B)** Distal convoluted tubule cell and segment specific DEGs were subjected to dynamic enrichment (A) and module analysis (B).

**Supplementary Figure 12:**
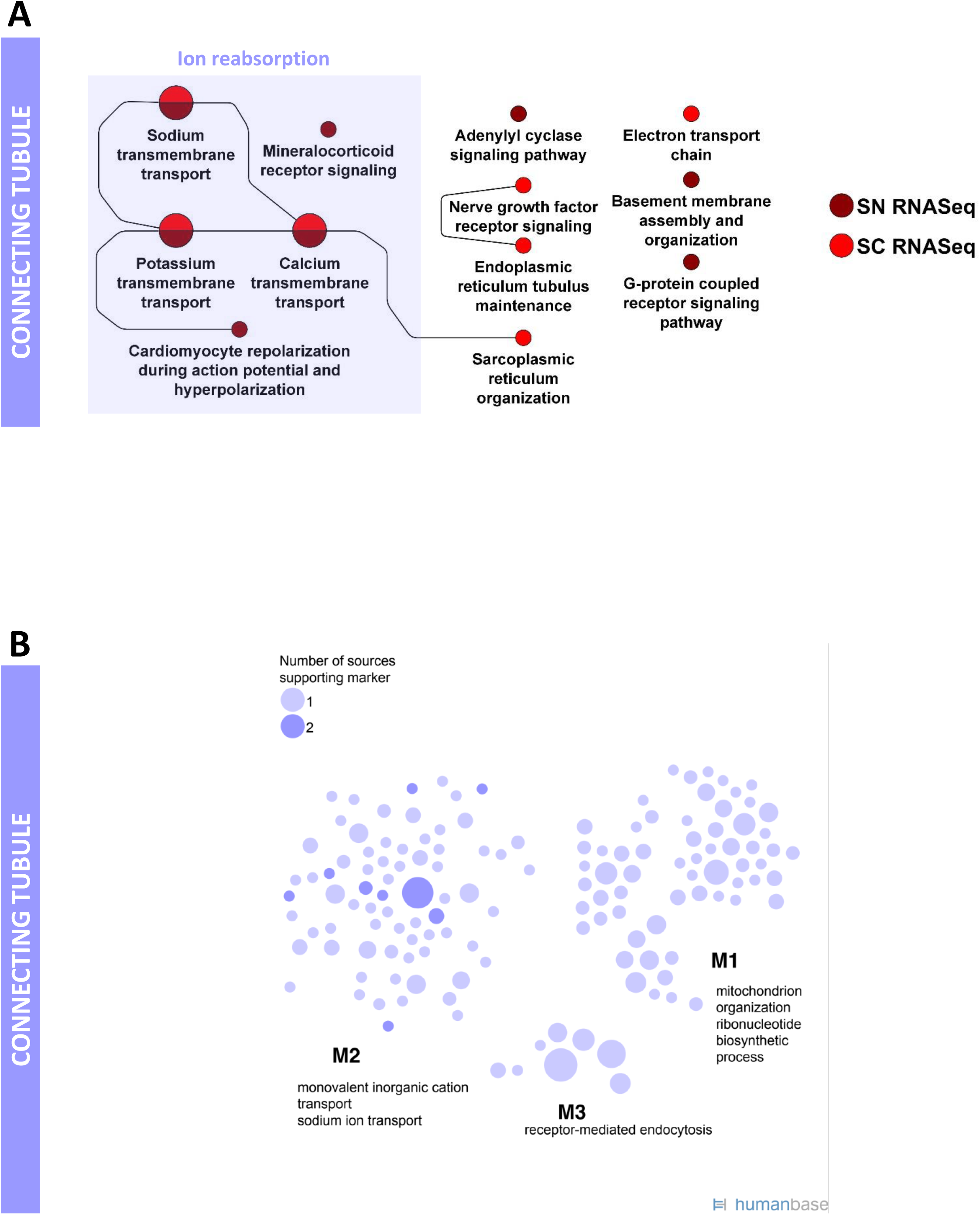
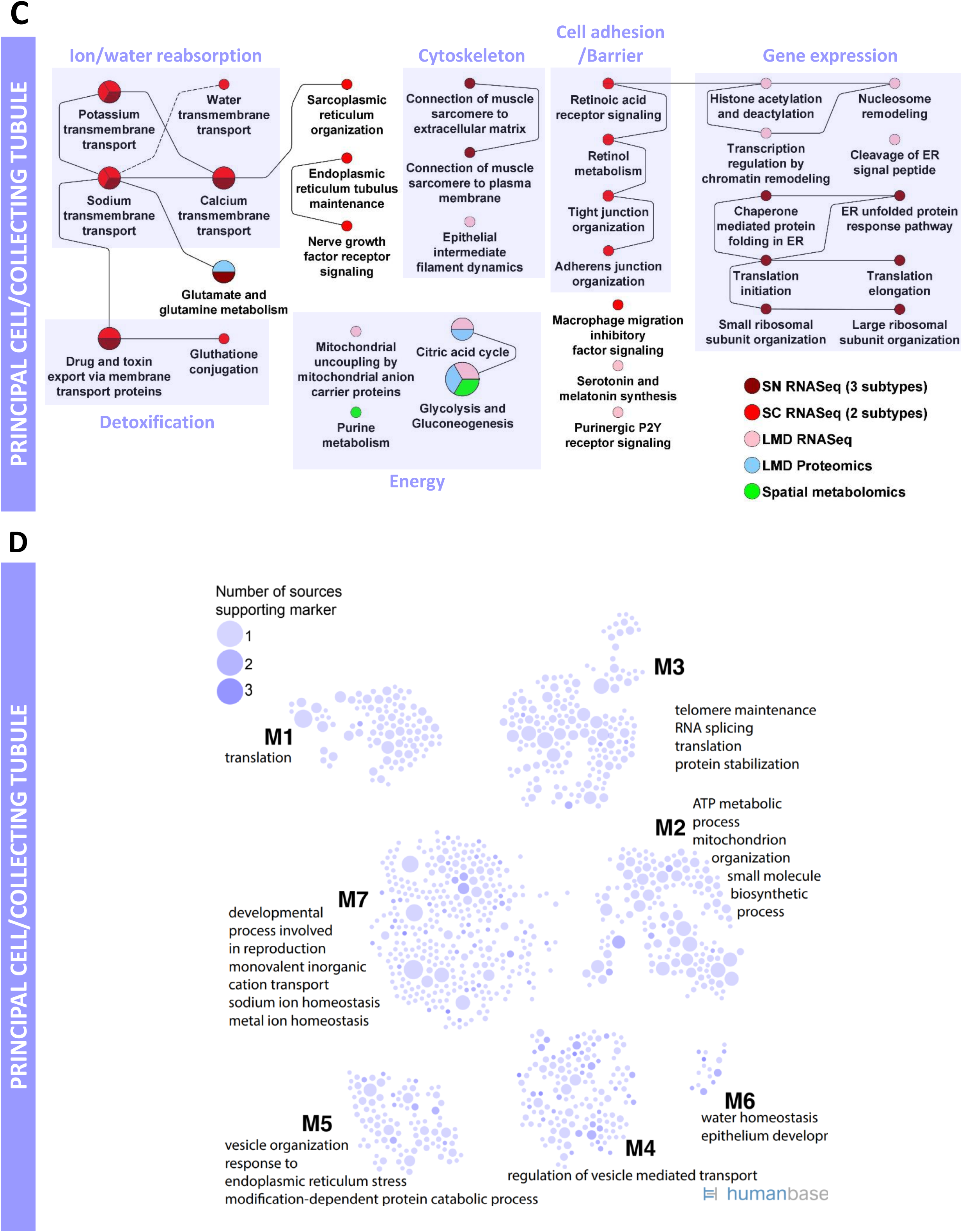
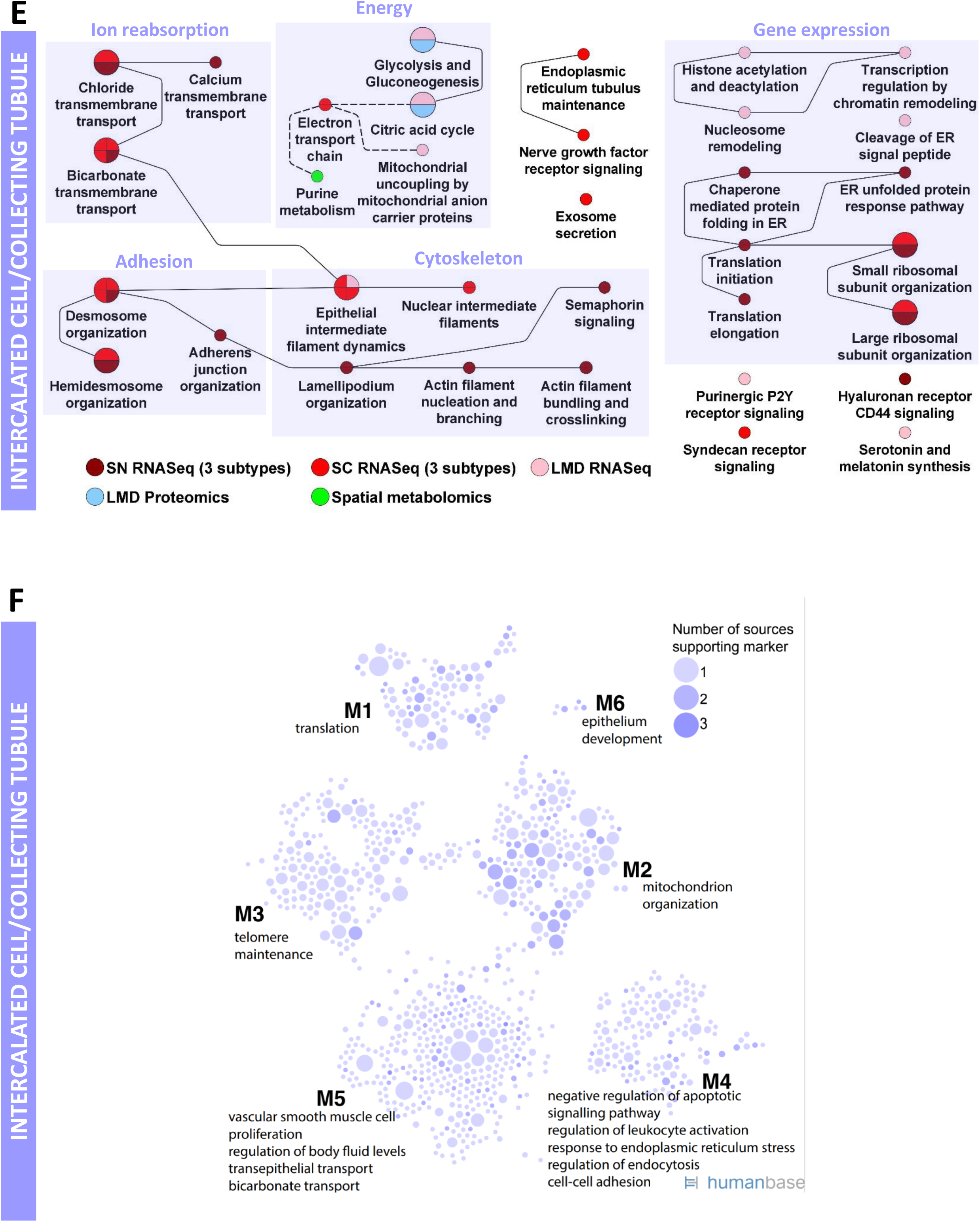
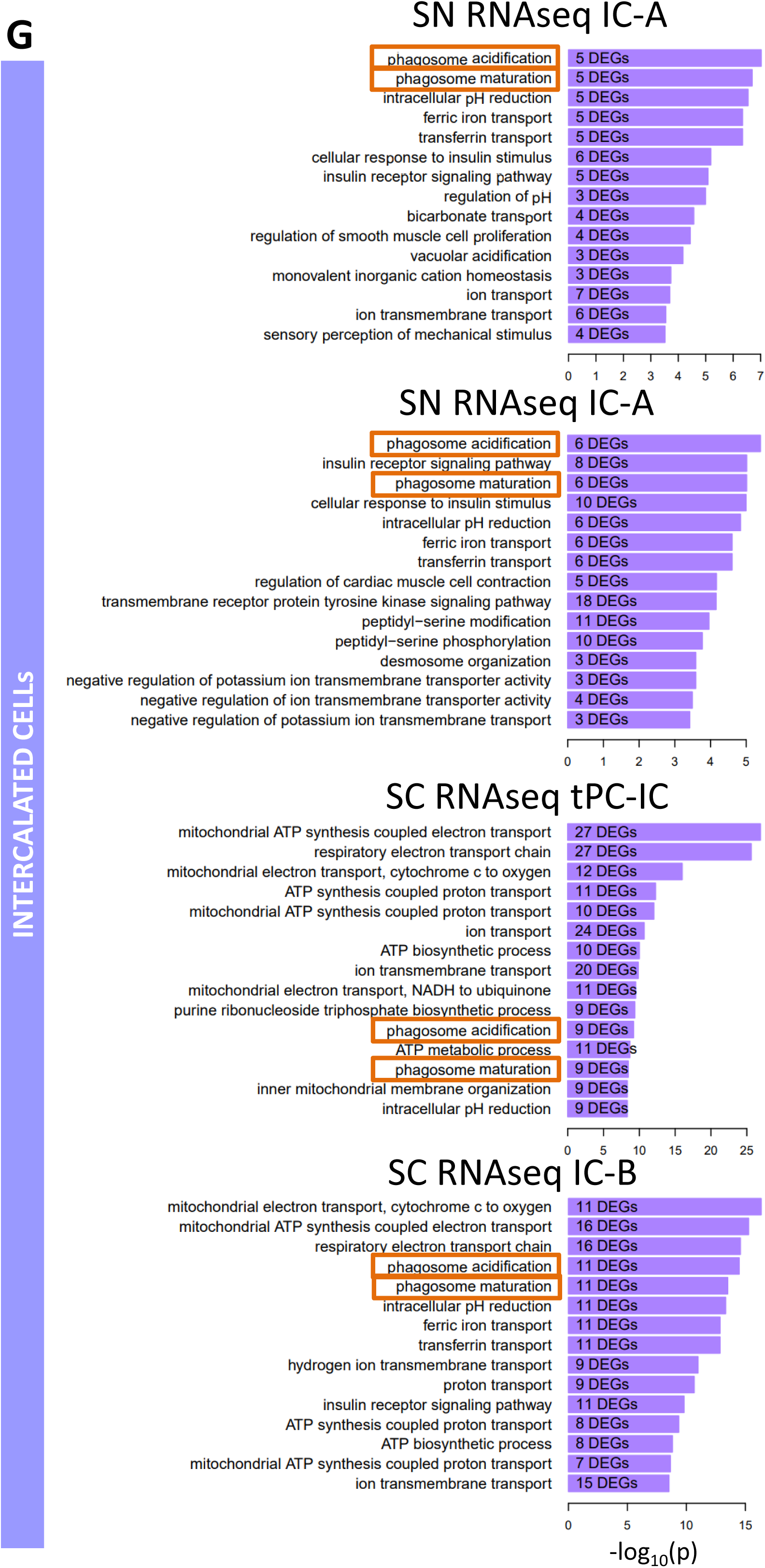
Enrichment analysis for the collecting duct. **(A/B)** Connecting tubule cell specific DEGs were subjected to dynamic enrichment (A) and module analysis (B). **(C/D)** Principal cell and collecting duct specific DEGs were subjected to dynamic enrichment (C) and module analysis (D). **(E/F)** Intercalated cell and collecting duct specific DEGs were subjected to dynamic enrichment (E) and module analysis (F). **(G)** Enrichment analysis of the marker genes for 4 different intercalated cell subtypes from sn and sc RNAseq using Gene Ontology Biological Processes identifies the pathways ‘Phagosome acidification’ and ‘Phagosome maturation’.

**Supplementary Figure 13:**
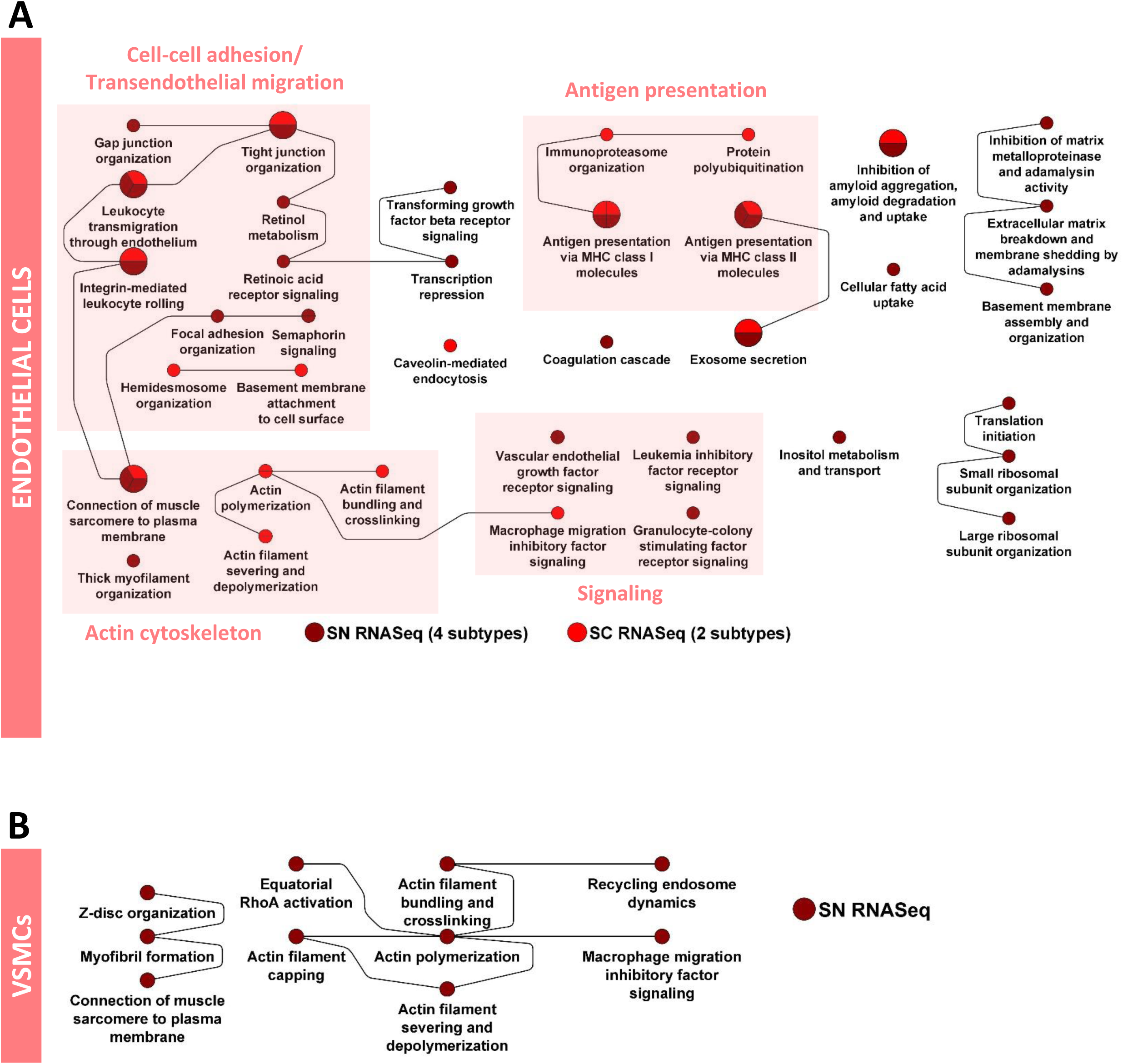
Enrichment analysis for vascular cells. **(A)** Endothelial cell specific DEGs were subjected to dynamic enrichment. **(B)** Similarly, vascular smooth muscle cell specific DEGs were subjected to dynamic enrichment analysis.

**Supplementary Figure 14:**
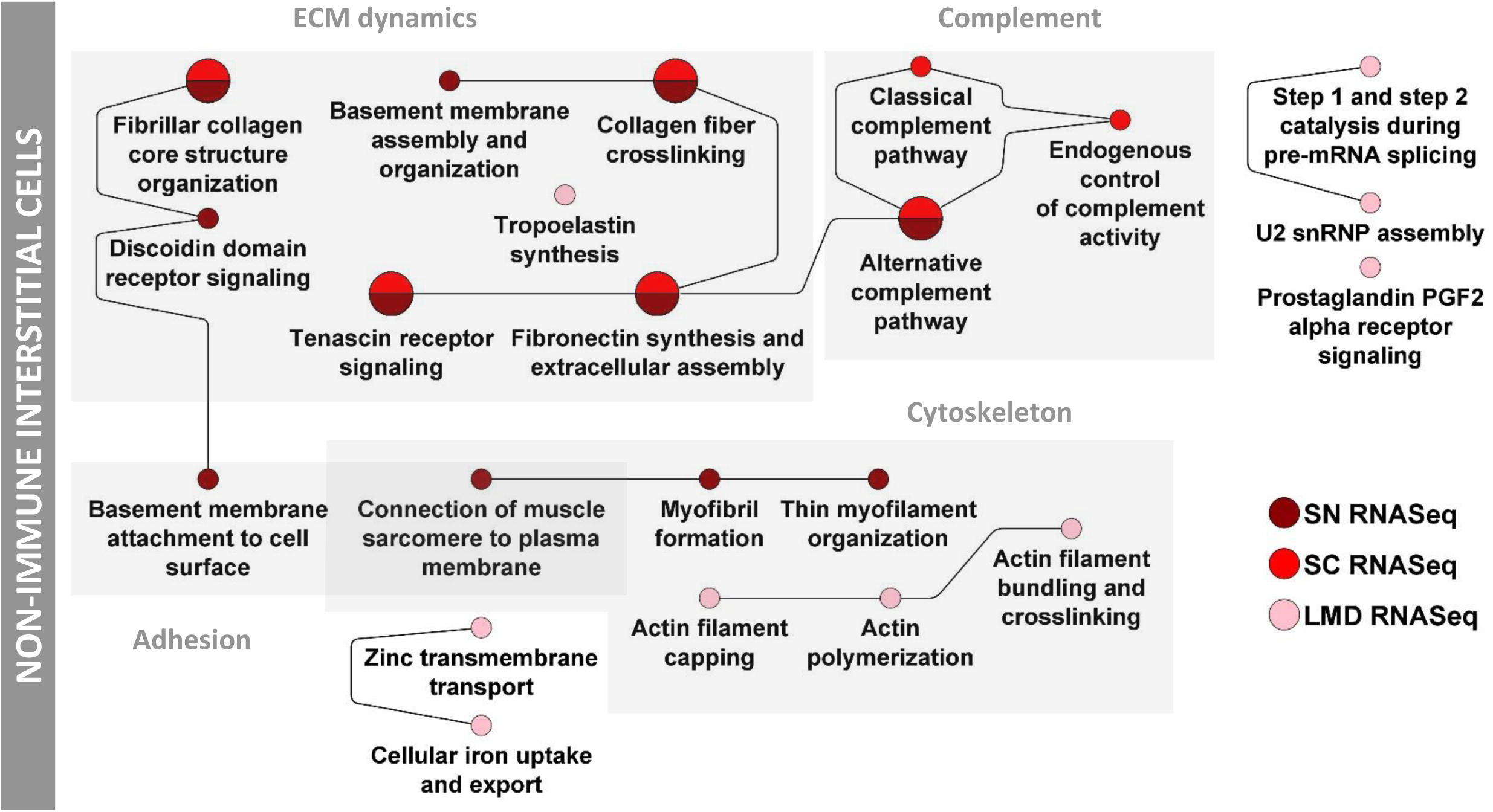
Enrichment analysis for interstitial cells. **(A)** Interstitial fibroblast cell and segment specific DEGs were subjected to dynamic enrichment analysis.

**Supplementary Figure 15:**
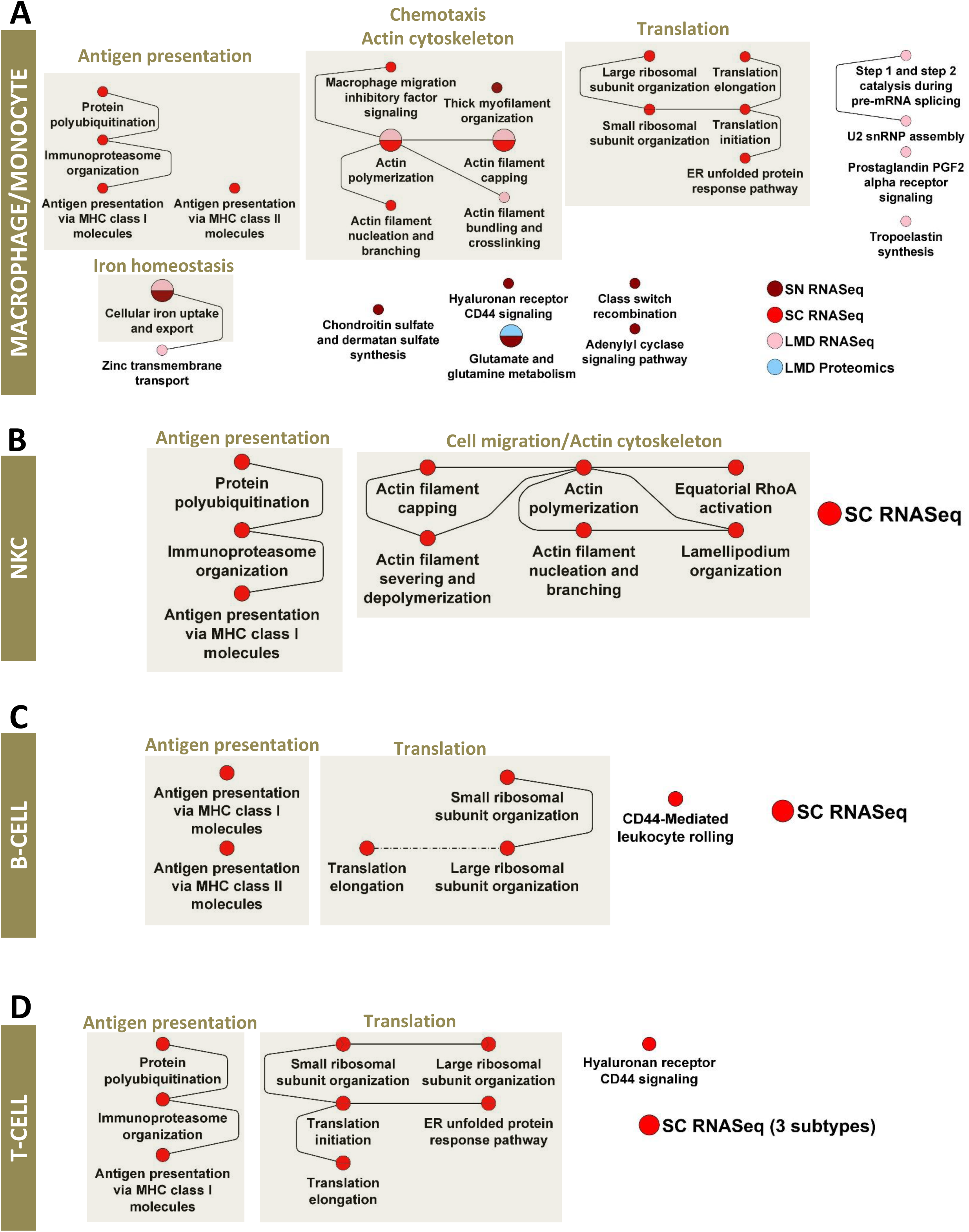
Enrichment analysis for immune cells. **(A)** Macrophage/Monocyte, **(B)** Natural Killer cell, **(C)** B-cell, and **(D)** T-cell specific DEGs were subjected to dynamic enrichment analysis.

**Supplementary Figure 16:**
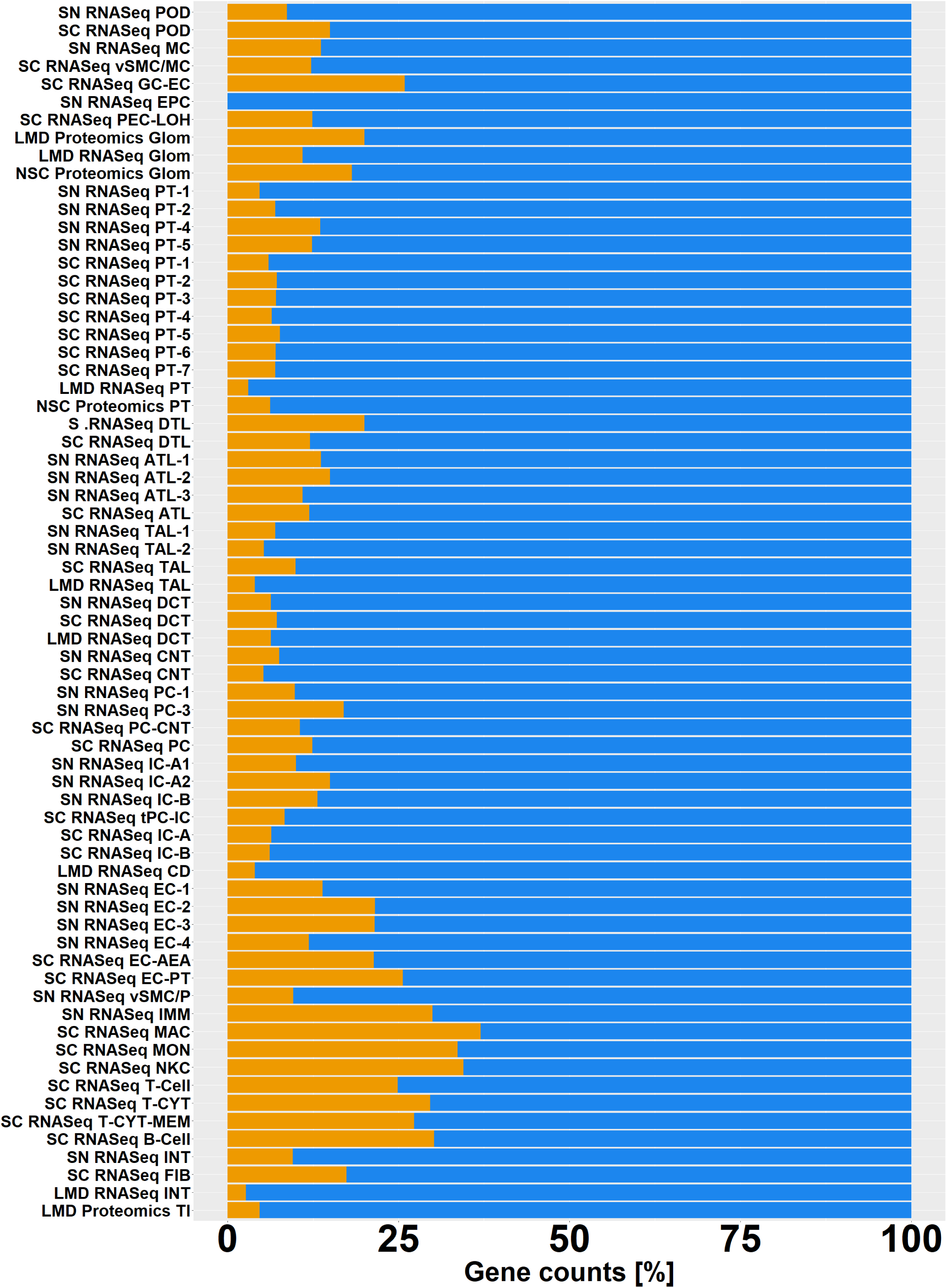
Expression of immune related genes in the kidney cell types. Using all genes that are assigned to the Gene Ontology Biological Process “immune system process” or any of its children processes based on the “is_a” or “part_of” relationships, we documented the percentage of immune system related genes (orange) in all cell type, subtype and segment-specific marker genes and proteins. See figure 2A for cell type abbreviations.

**Supplementary Figure 17:**
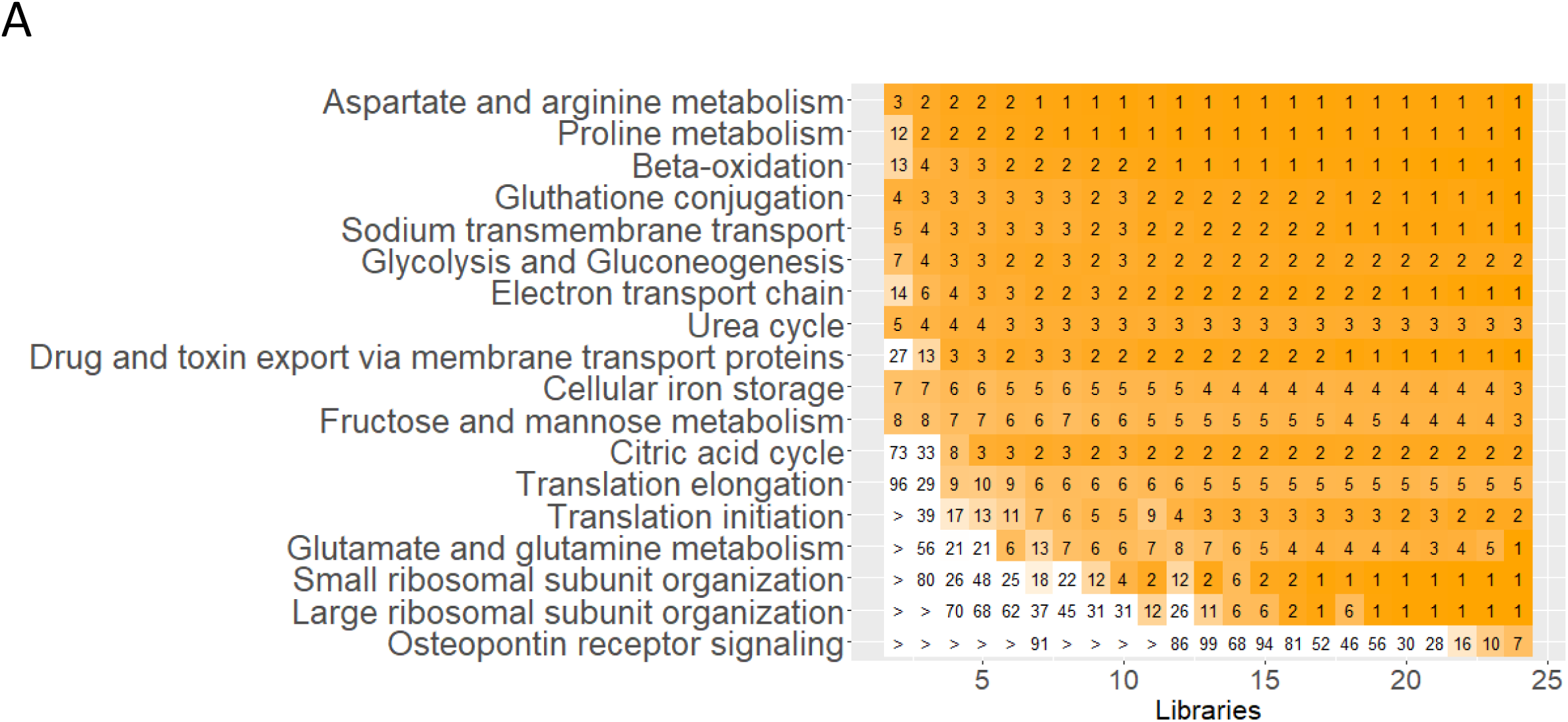
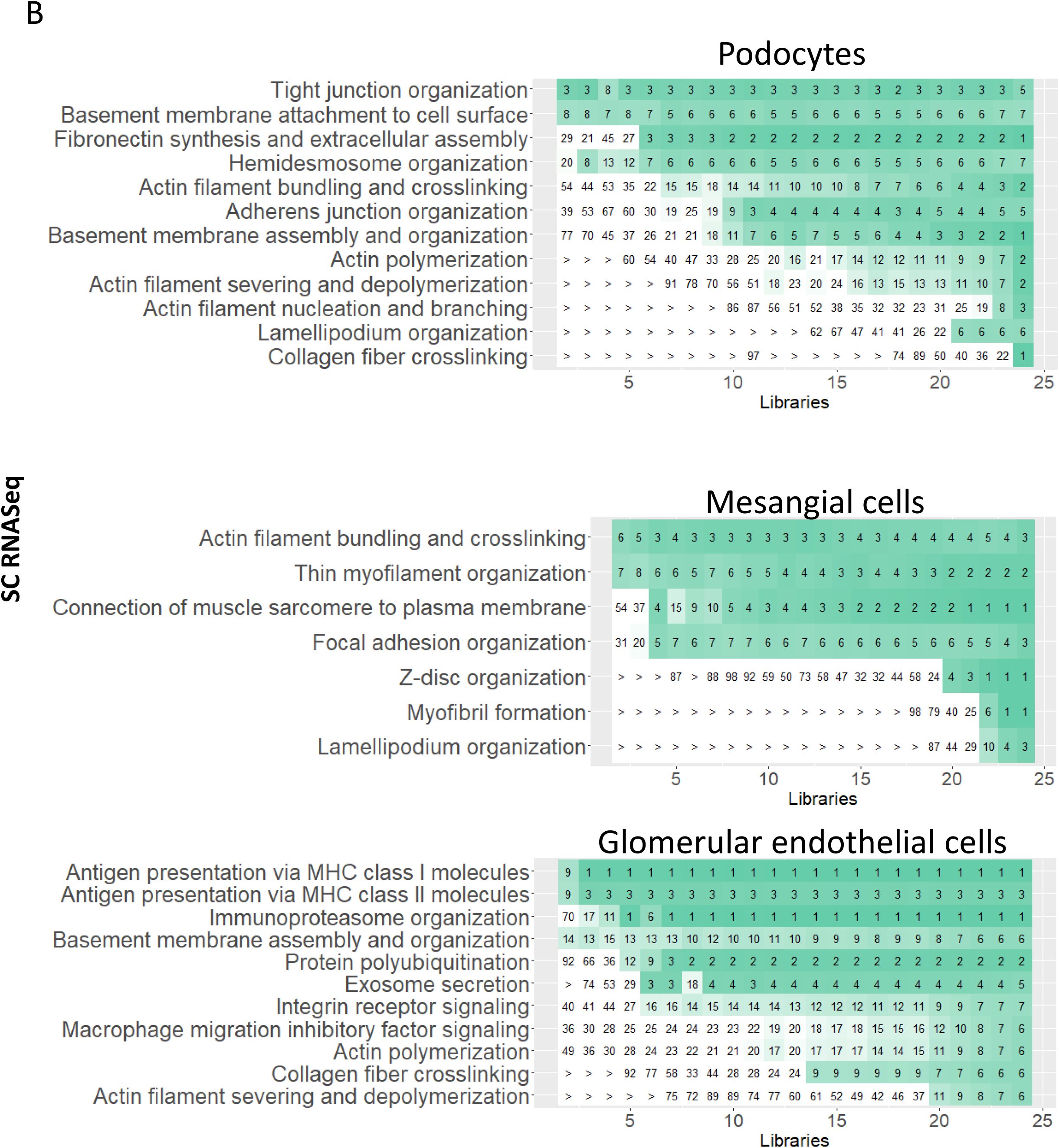
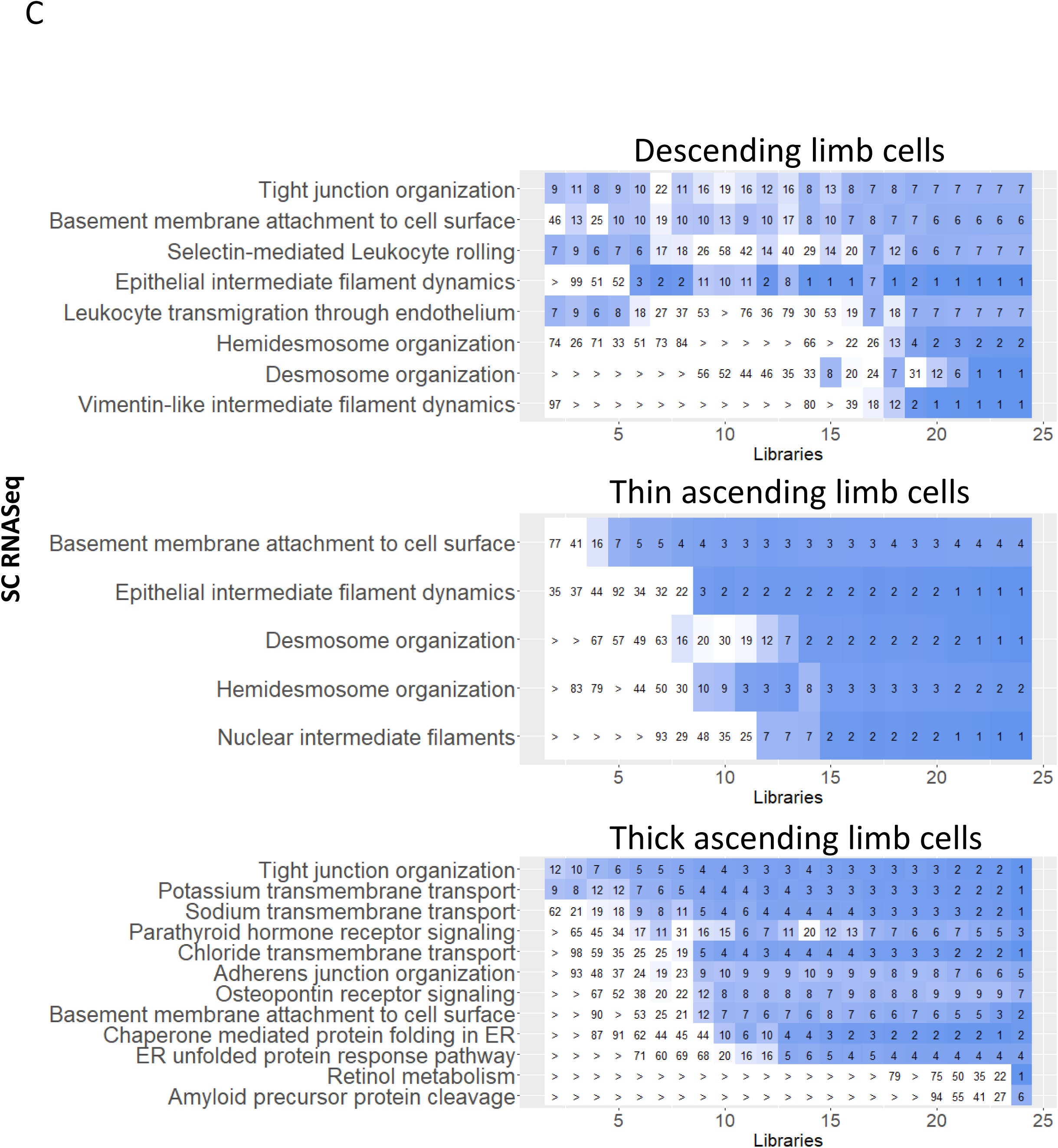
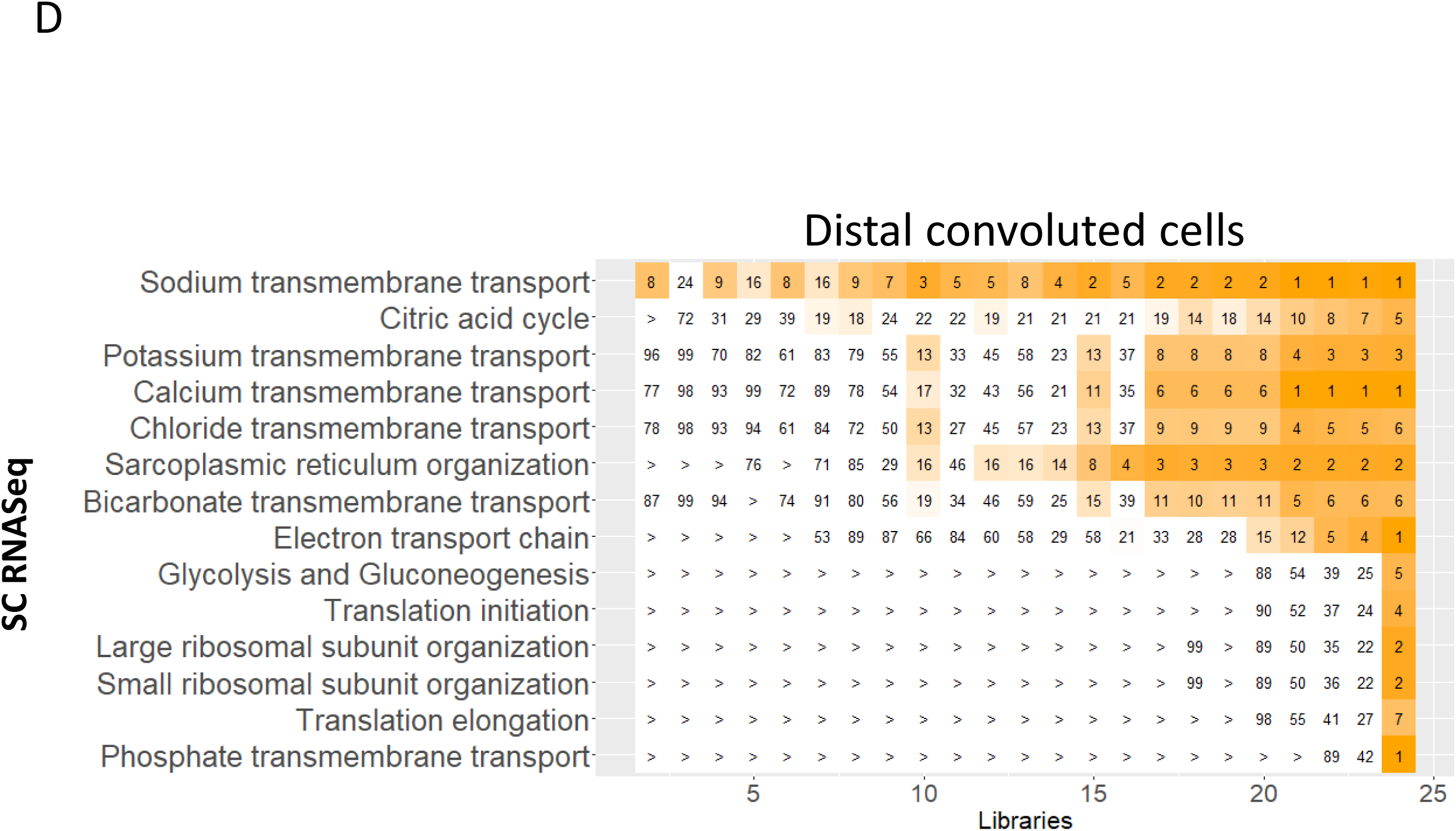
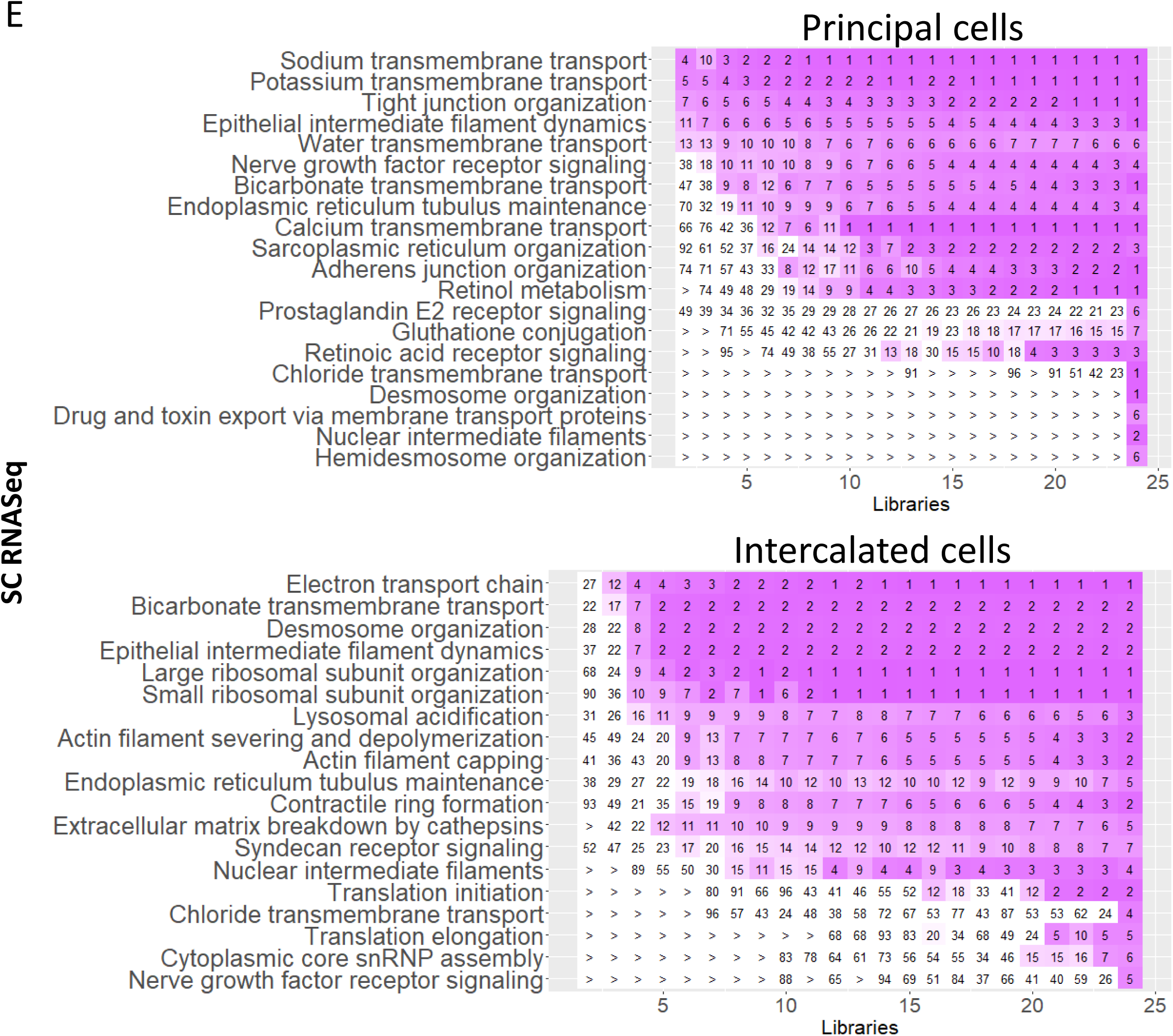
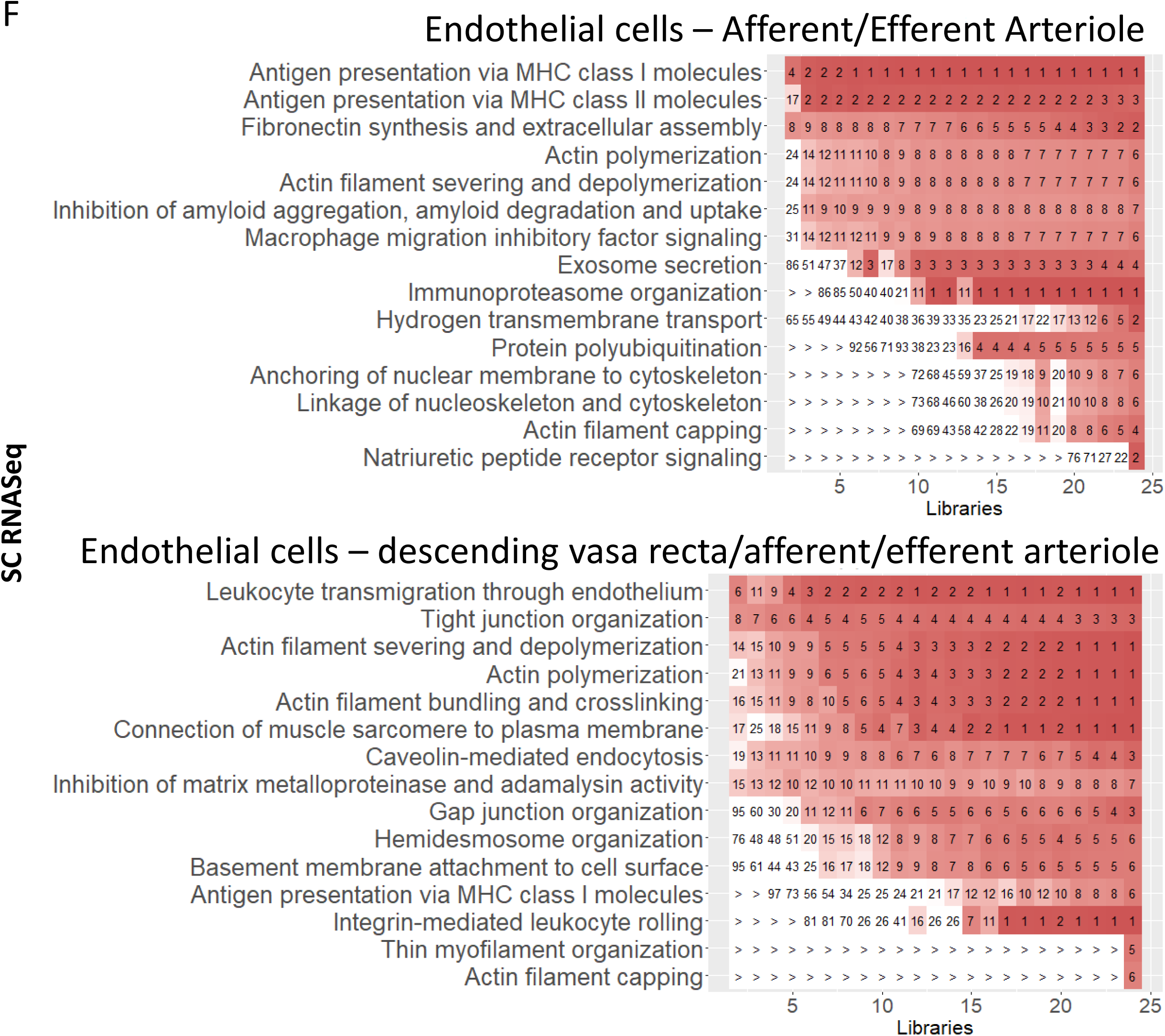
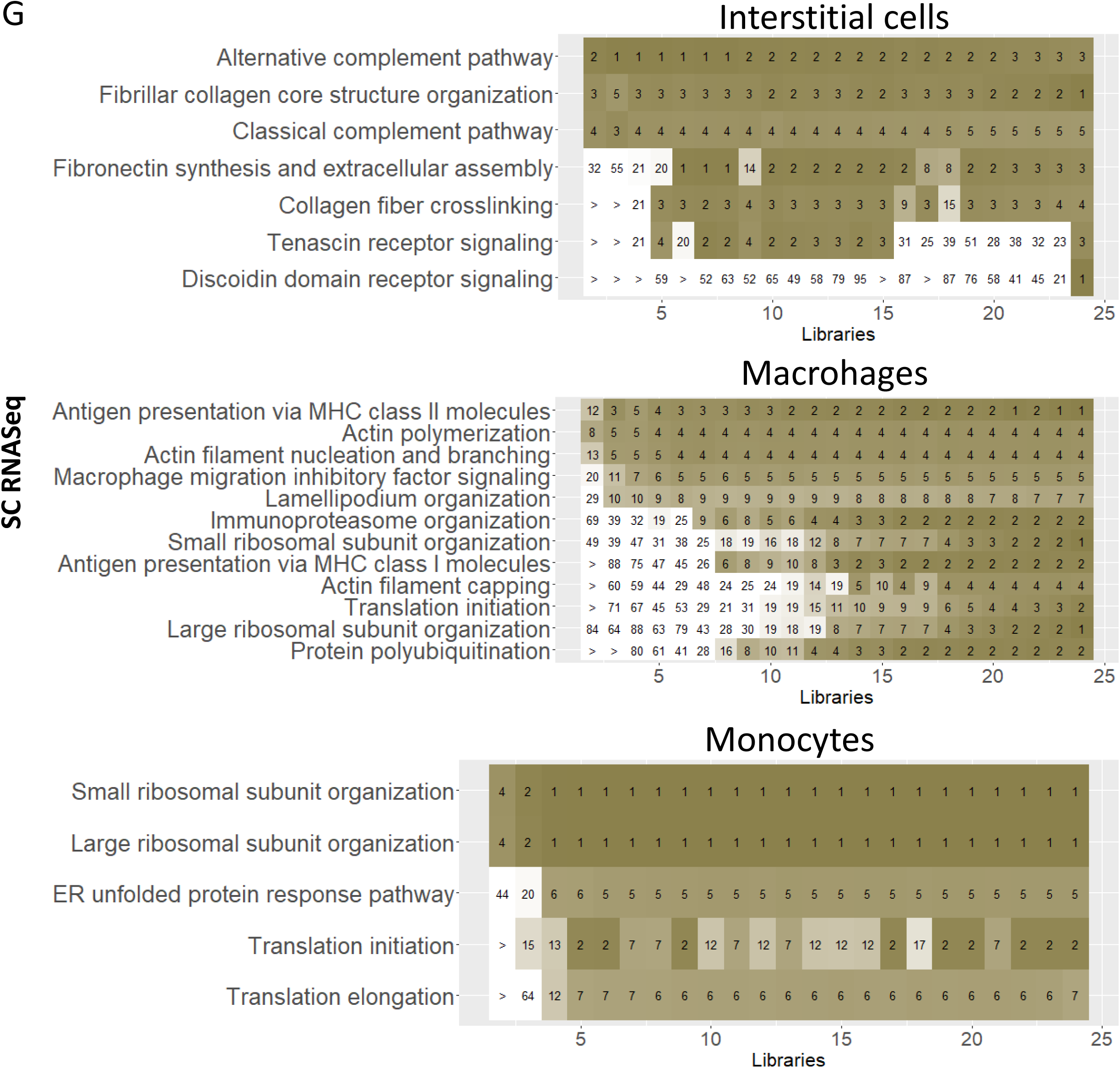
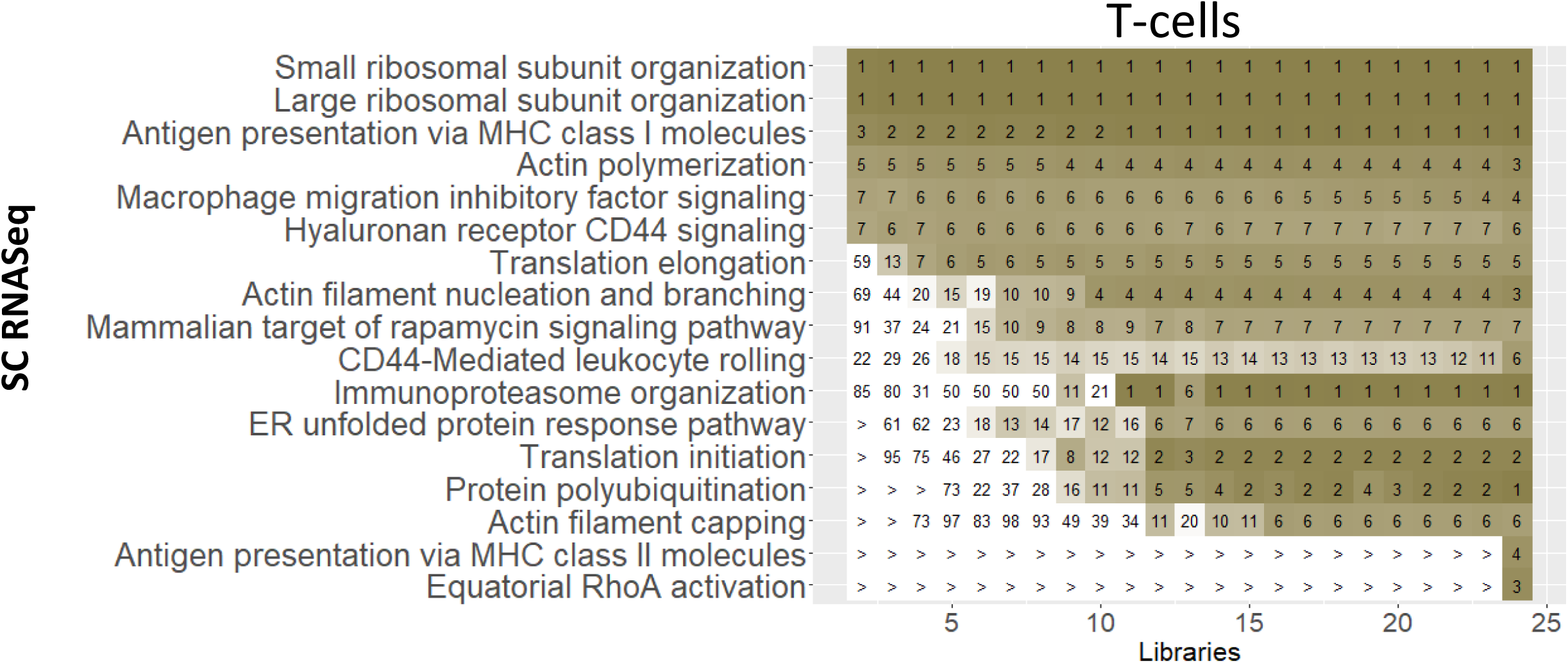
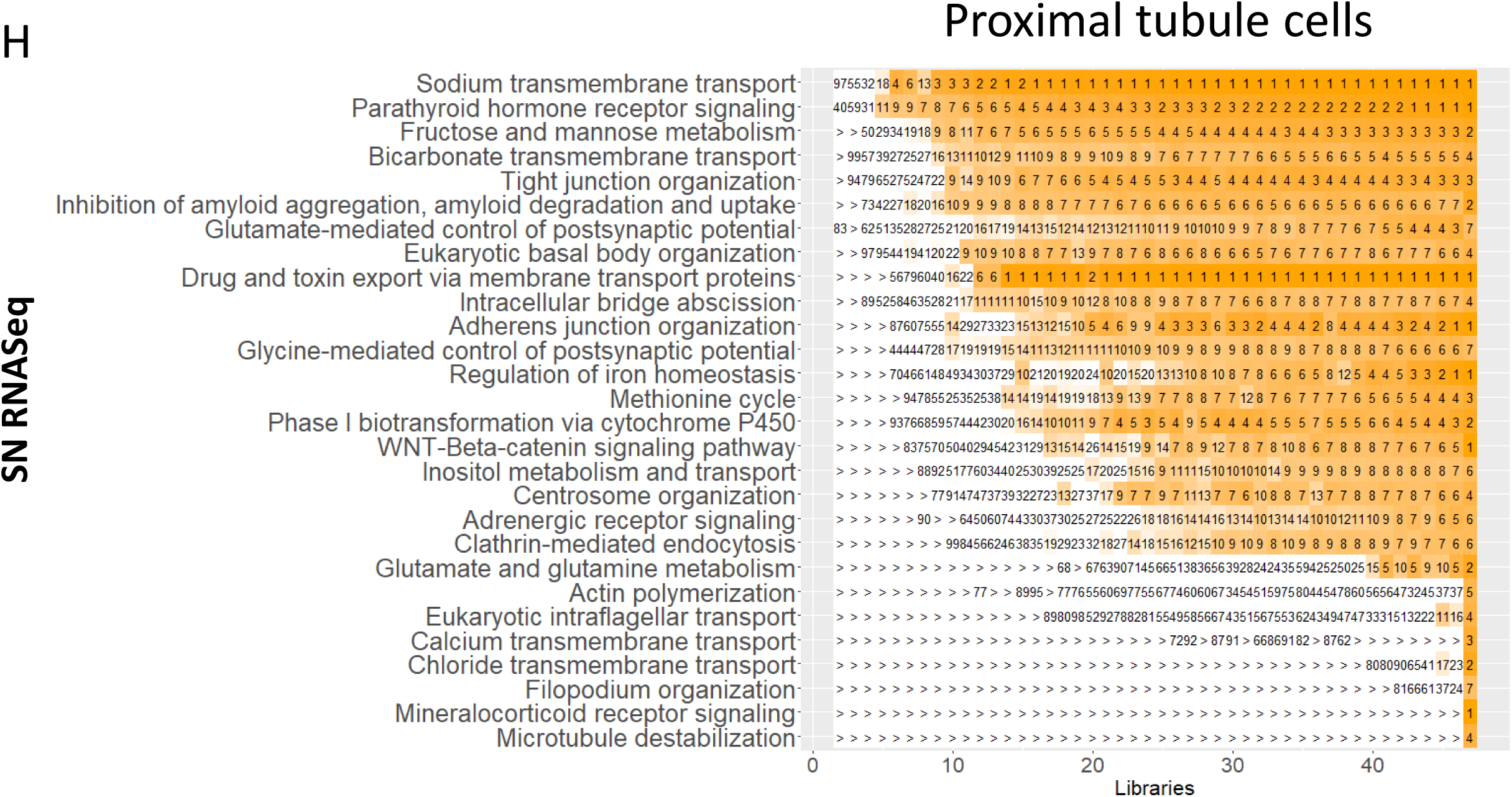
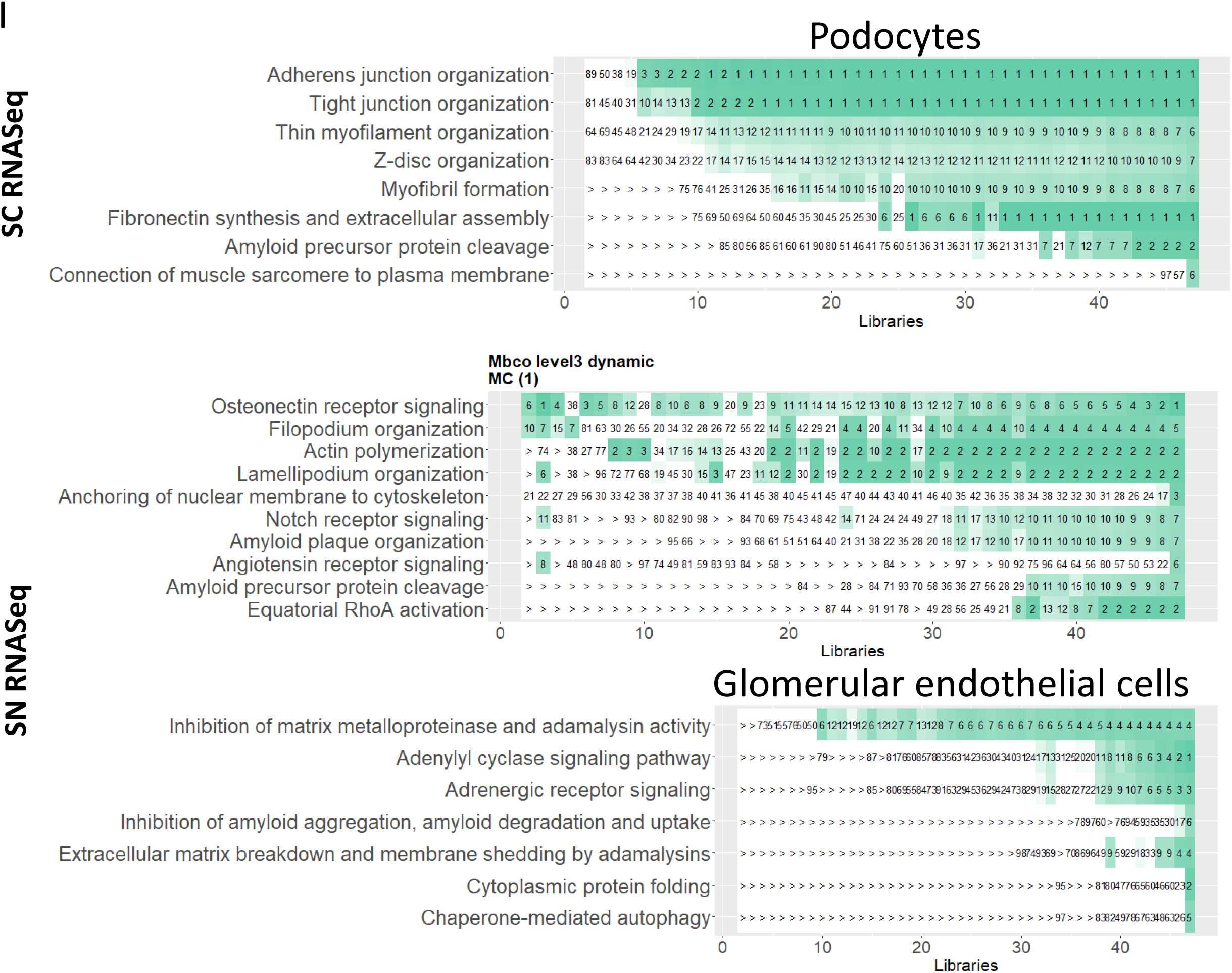
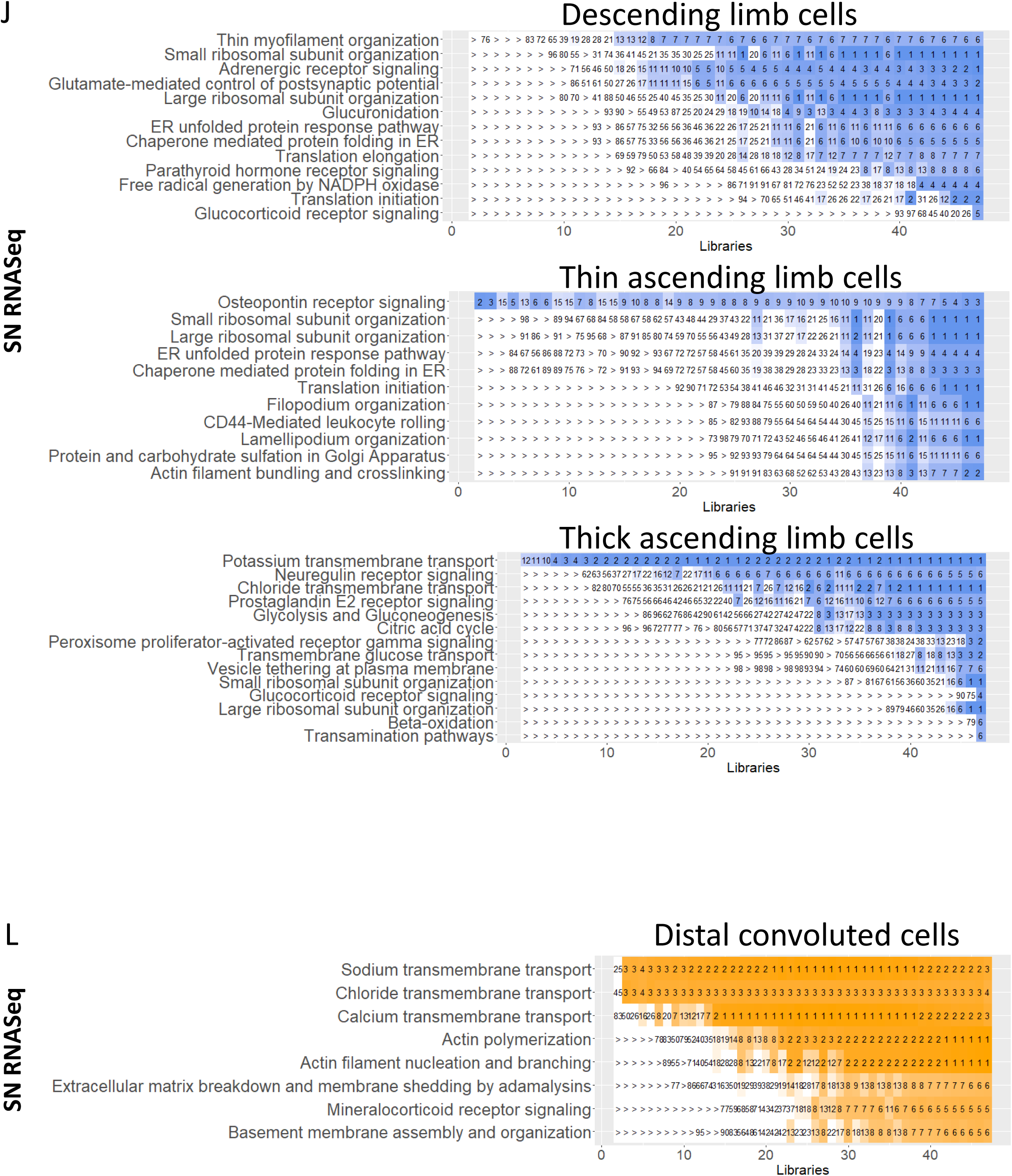
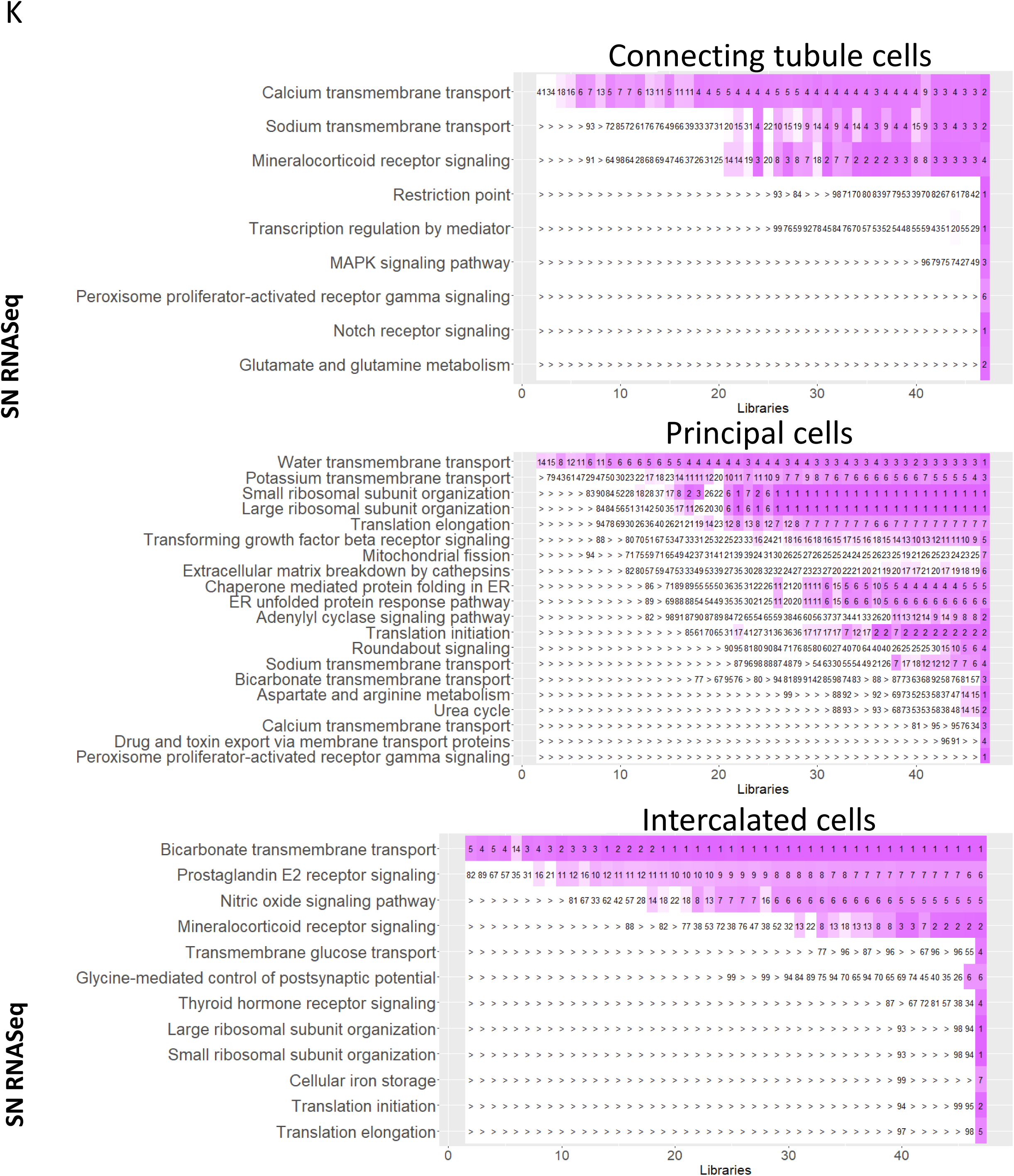
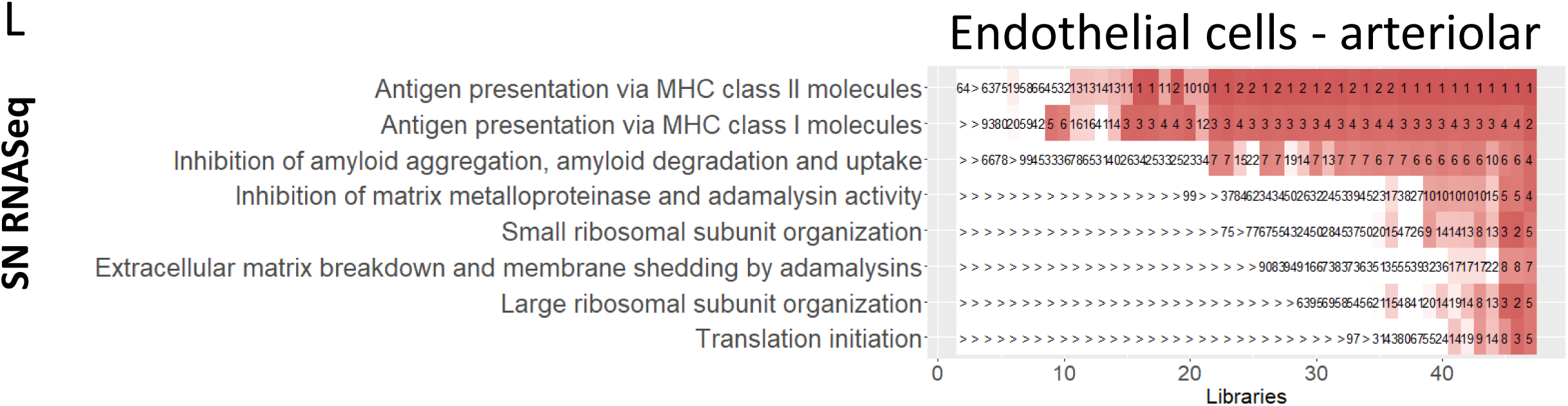
Cellular key functions are most consistently predicted by downsampled sc and sn RNAseq datasets. To analyze the reliability of predicted cell type-specific biology we subjected the top 300 cell-type specific marker genes that were obtained from the full or down-sampled sc and sn RNAseq datasets (Suppl. Figure 5C and 5D, respectively) to dynamic enrichment analysis. SCPs that were among the top 7 predictions for the full sc and sn RNAseq were identified. We identified the dynamic enrichment ranks of these SCPs in the down-sampled datasets and averaged them across all datasets with the same number of libraries. Color scale ranges from 1 (dark green/orange/purple) to 21 or higher (white). Notify that SCPs predicted for the full datasets are not necessarily the same as the one documented in figure 6, since the 2124, 4447 and 721 individual full and downsampled datasets were analyzed using an automated pipeline that did not allow manual *ad hoc* optimization. The first set of subfigures shows the predicted SCPs identified from the sc RNAseq dataset for **(A)** proximal tubule cells, **(B)** glomerular cell types, **(C)** cell types of the Loop of Henle, **(D)** of the distal convoluted tubule, **(E)** of the collecting duct, **(F)** vascular cells and **(G)** non-immune and immune interstitial cells. The second set of subfigures shows the predicted SCPs identified from the sn RNAseq dataset for **(H)** proximal tubule cells, **(I)** glomerular cell types, **(J)** cell types of the Loop of Henle, **(K)** of the distal convoluted tubule, **(L)** of the collecting duct and **(M)** vascular cells.

**Supplementary Figure 18:**
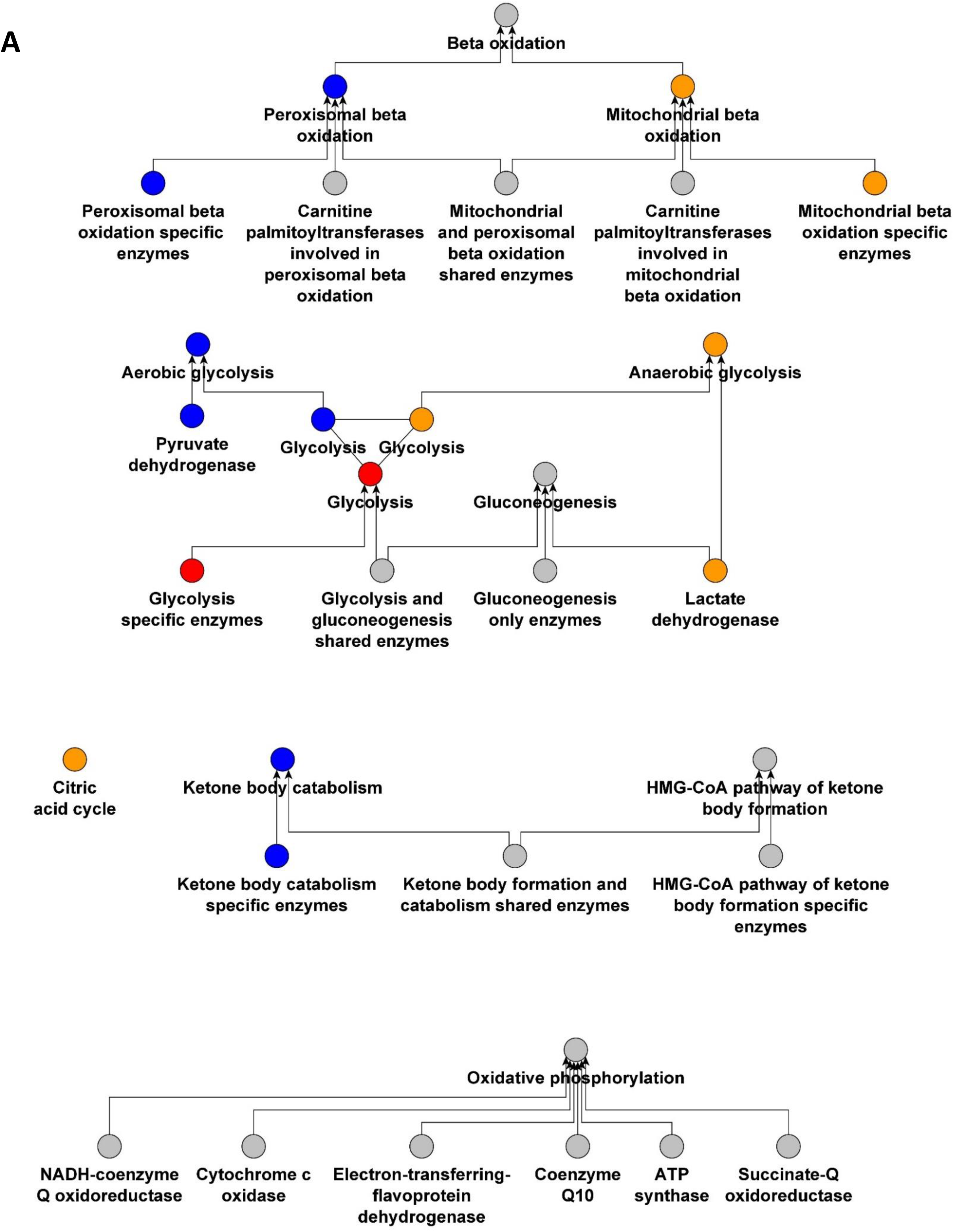
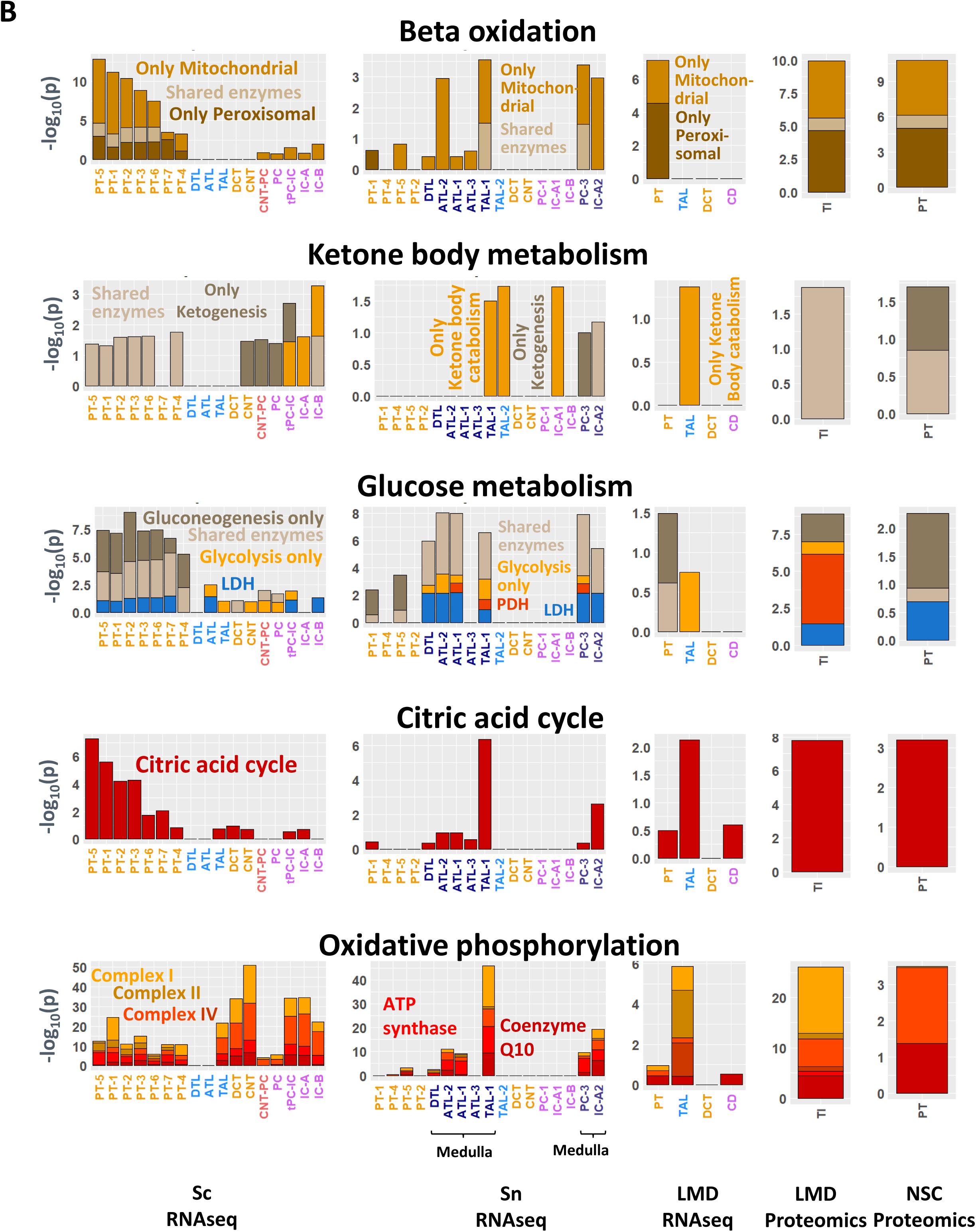
Prediction of cellular dependencies on aerobic and anaerobic metabolic pathway activities. **(A)** We designed a small ontology that allows distinguishing between aerobic and anaerobic as well as catabolic and anabolic reactions. Shown is the annotated pathway hierarchy. Colored pathways indicate parent and child pathway pairs, where the child contains only enzymes that are specifically involved in the function of its parent and of any other parent. Pathways were populated with genes by literature curation. Parents are populated with all genes of the child pathways. **(B)** Top 500 cell type, cell subtype and subsegment specific marker genes and proteins were subjected to enrichment analysis using the leaf pathways shown in A. Initial enrichment results determined with pathways were used for the analysis shown in figure 7. For each cell type, subtype and subsegment we only considered a higher level pathway, if the child pathway that contains the enzymes specifically involved in the higher level pathway activity was also predicted (as indicated by the colored pathway pairs in A). Cell types that contain many cells obtained from medullary samples are marked. See figure 2A for cell type abbreviations.

**Supplemental figure 19:**
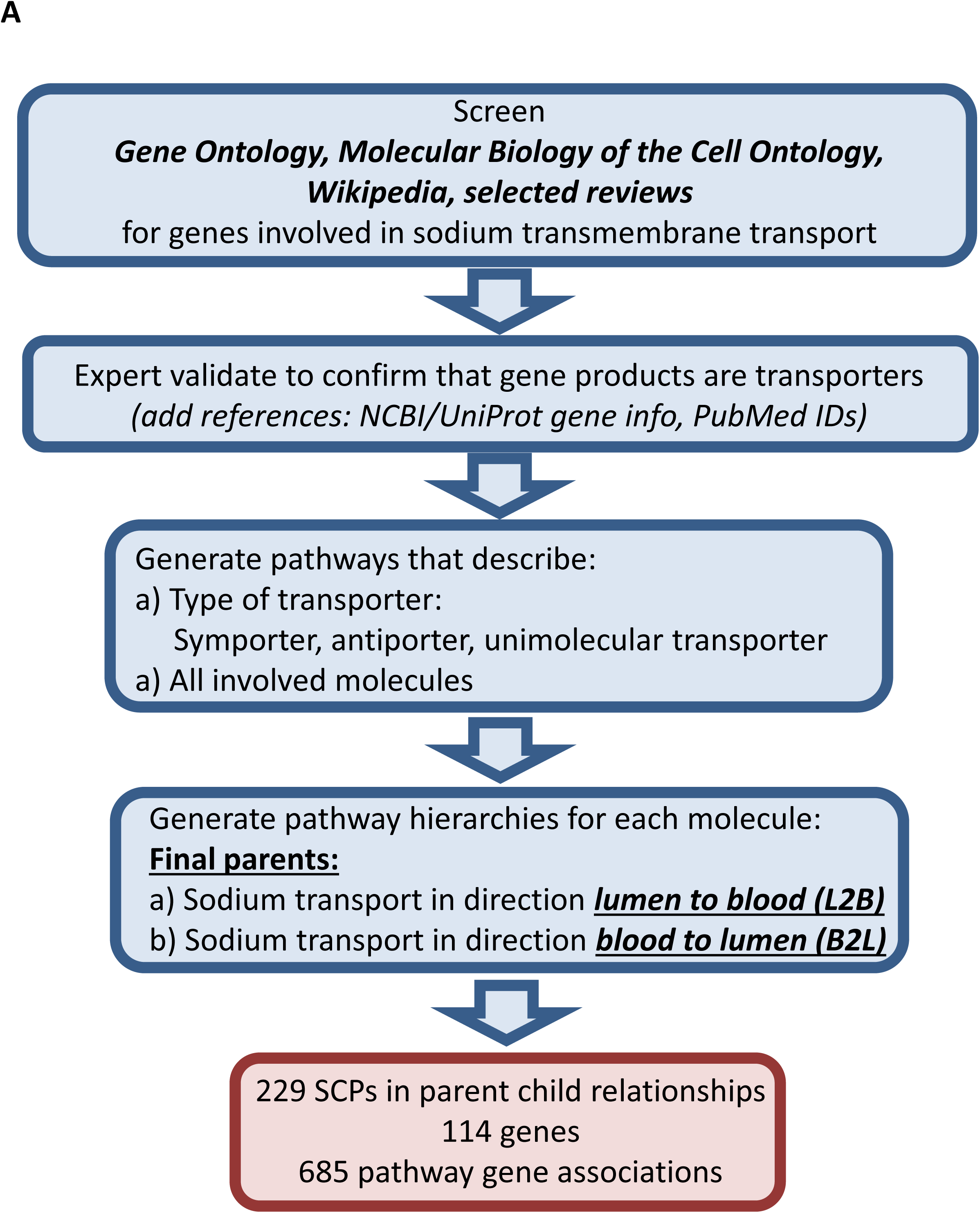
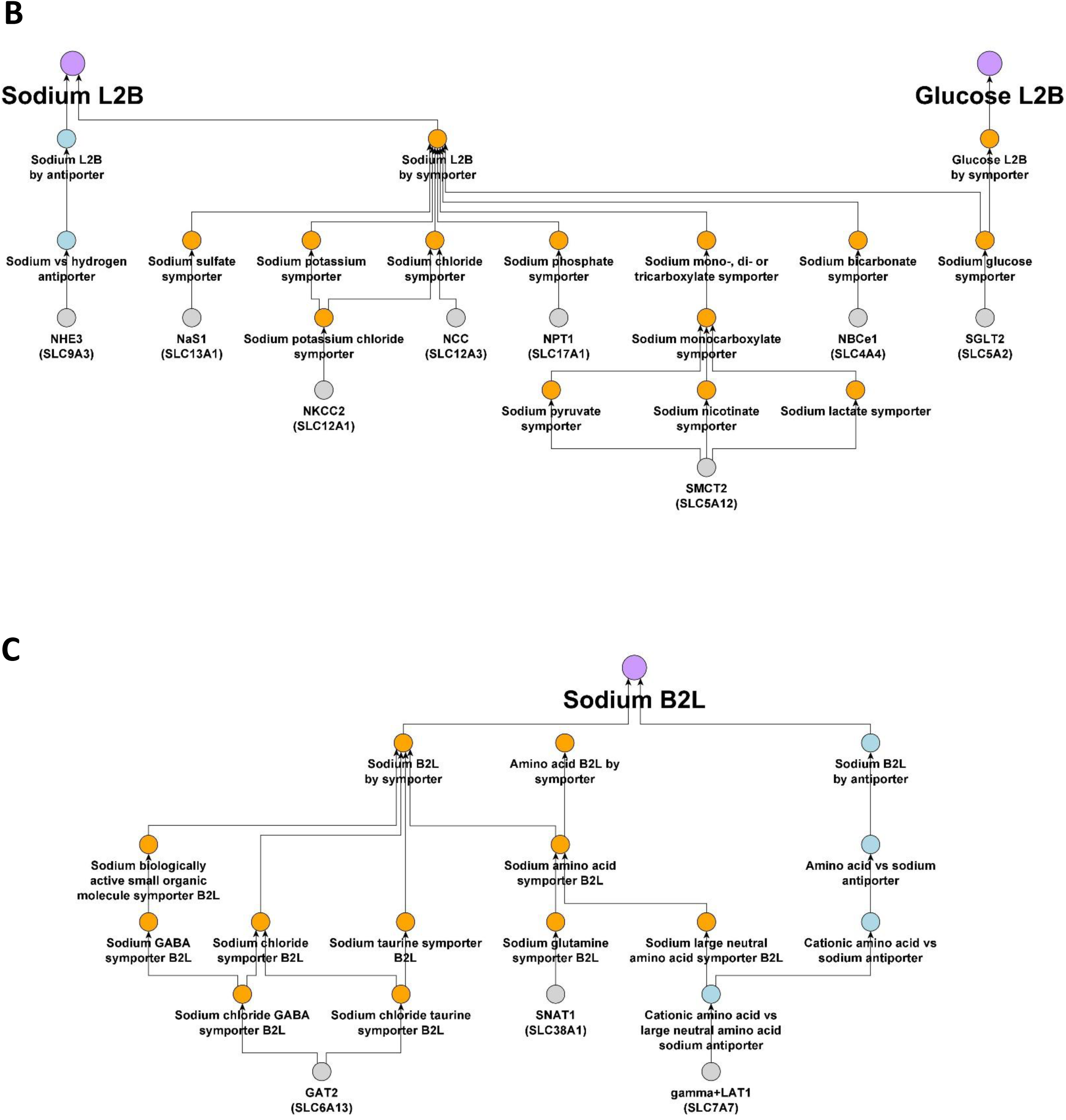
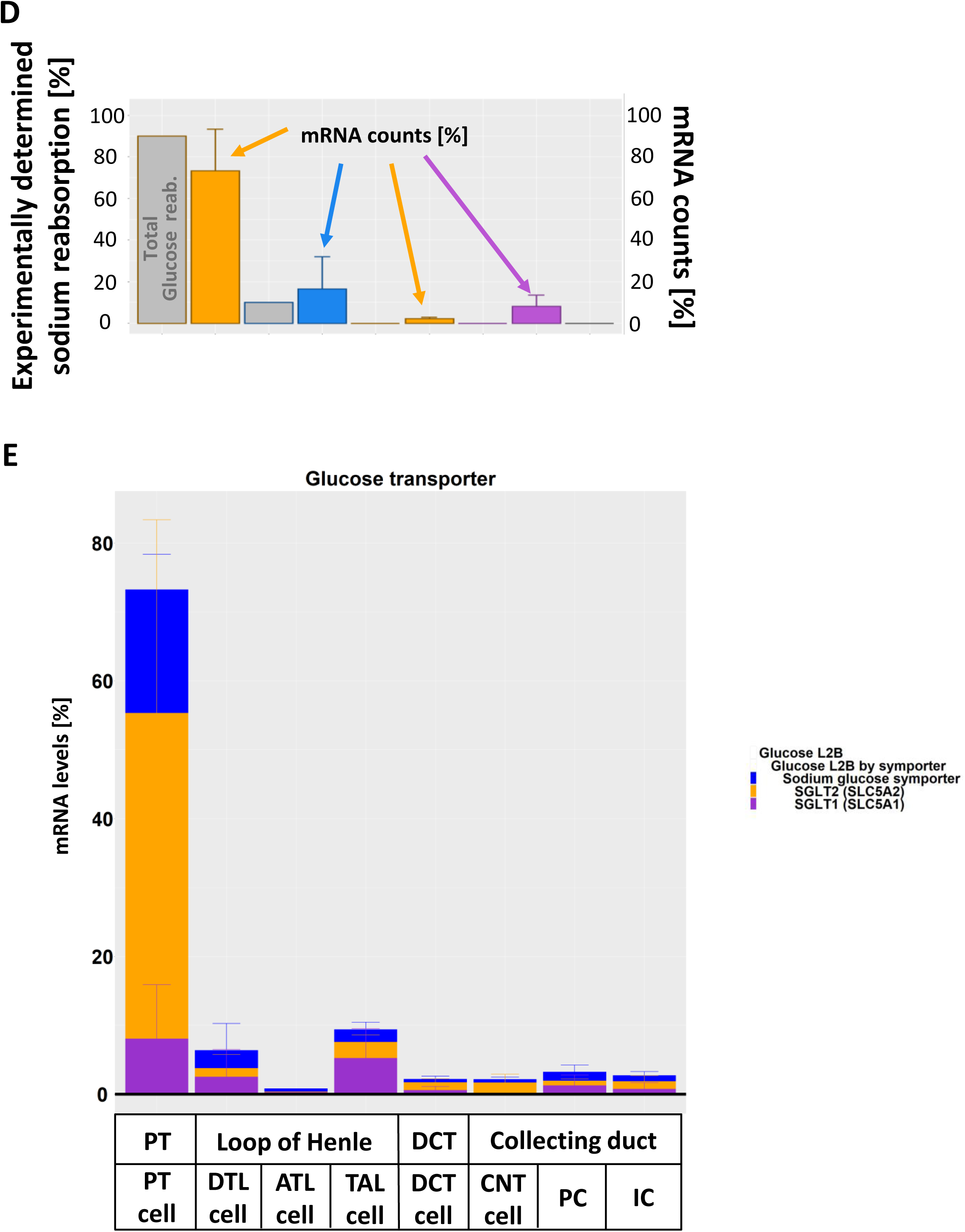
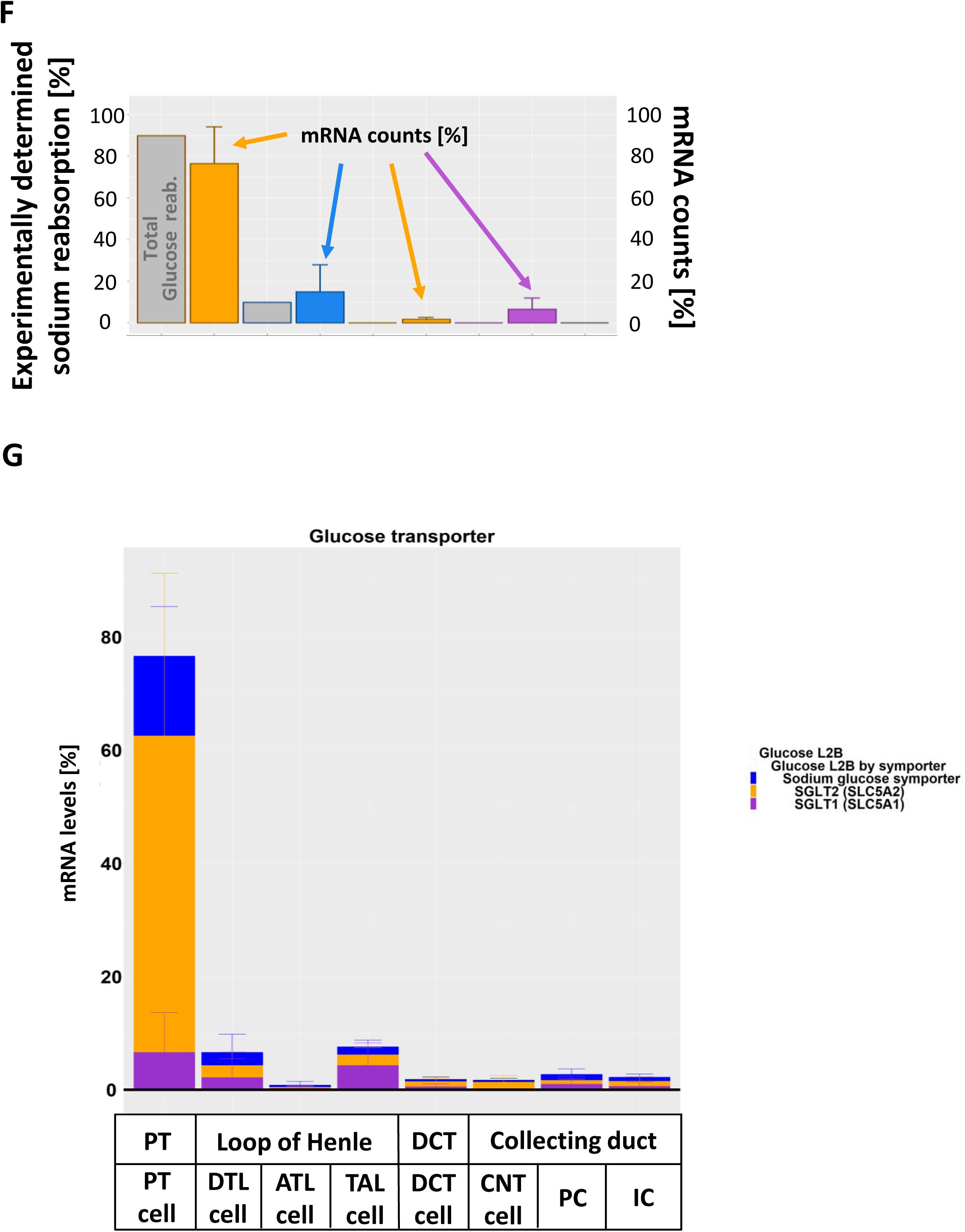
Generation of an ontology for transmembrane ion and molecule movements. **(A)** Flow chart documenting the steps involved in the generation of the ontology. (B) Shown are example transporters (gray) involved in sodium and glucose lumen-to-blood (L2B) transport and how they integrate into the hierarchy to finally converge on sodium and glucose lumen-to-blood transport. Symporter mechanisms are colored in orange, antiporter mechanisms in blue. **(C)** The figure illustrates SCPs involved in sodium blood-to-lumen (B2L) transport and their integration into the SCP hierarchy. **(D)** Reabsorption capacities for glucose transmembrane transport were calculated using the three sn RNAseq datasets as described in figure 8 and compared to experimentally determined glucose reabsorption profiles. Since only one physiology text book (*94*) documented the glucose reabsorption profiles, there is no standard deviation for the experimental values. Facilitated glucose transporters were excluded. **(E)** As for sodium, we analyzed the transport mechanisms involved in glucose reabsorption. **(F)** We compared the reabsorption capacities that were calculated using the three sn and the sc RNAseq datasets with the experimental reabsorption profiles, **(G)** followed by visualization of the individual transport mechanisms for glucose.

**Supplementary Figure 20:**
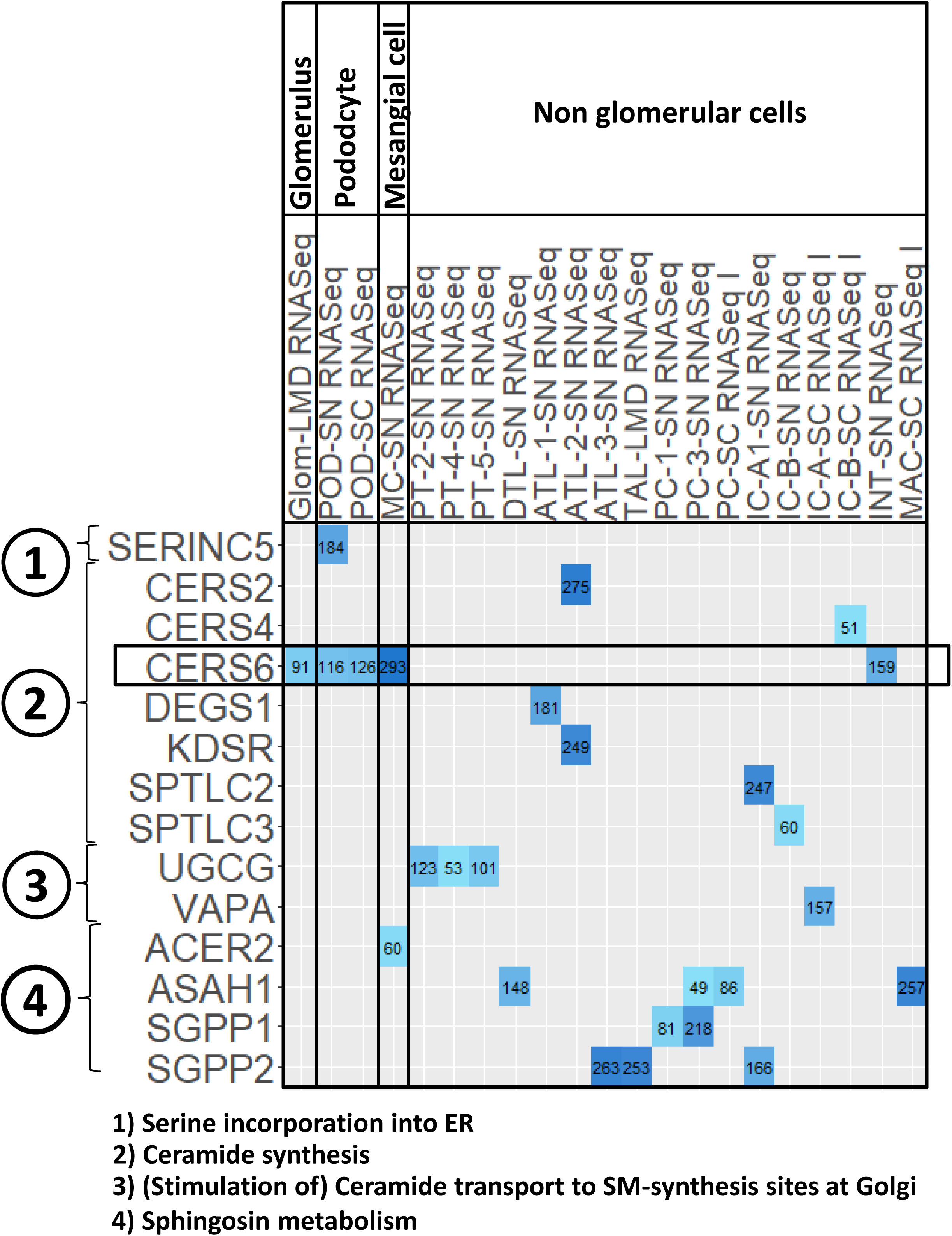
Expression of genes involved in sphingomyelin synthesis and sphingosine metabolism in all kidney cell types and segments. Expression of curated enzymes was detected in the indicated cell types/segments. Genes were ranked by significance and ranks were added to the figure.

## Supplementary Table Captions

**Supplementary Table 1.** Samples used for different analytical pipelines.

**Supplementary Table 2.** Laser microdissected (LMD) RNAseq gene expression

**Supplementary Table 3.** Laser microdissected (LMD) Proteomics protein expression

**Supplementary Table 4.** Near Single Cell (NSC) Proteomics protein expression

**Supplementary Table 5.** Top 2,000 marker genes and proteins predicted by each assay for each analyzed cell subtype, cell type and tissue subsegment. Marker genes and proteins are differentially expressed genes (DEGs) and proteins (DEPs) that were obtained by comparing each cell type, cell subtype or subsegment to all other cell subtypes, types or subsegments. Initially, we duplicated all subsegmental datasets and added them to each data integration term that describes a cell type localized in that particular segment. For cell type specific assignments of the subsegmental data see results section, Figure 5C and Supplementary Figure 8.

**Supplementary Table 6.** # of significant marker genes that were subjected to dynamic enrichment analysis.

**Supplementary Table 7.** Dynamic enrichment analysis results of the top 300 marker genes and proteins. We duplicated all predictions based on the subsegment specific LMD RNAseq and Proteomics and the NSC Proteomics and added them to each integration term that describes a cell type localized in that particular segment. From these results we assigned cell type specificity to the predicted pathways as described in the results sections and documented in Figure 5C and Supplementary Figure 8. Notify that the columns “Experimental_symbols_count” and “Scp_symbols_count” contain the experimental and scp genes after removal of all those genes that are not part of the background list of genes (See methods for details). Hence, they can be smaller than the gene counts documented in supplementary table 6.

**Supplementary Table 8.** Spatial metabolomics metabolite correlations for subjects 18-139 (A), 18-142 (B) and 18-342 (C).

**Supplementary Table 9.** Gene Ontology enrichments for modules in the kidney-specific functional network of top DEGs and DEPs in PT, podocytes, and principal cells.

**Supplementary Table 10.** Literature curated cell-type specific essential genes used for cell type identification.

**Supplementary Table 11.** Enrichment analysis of the top 500 significant marker genes and proteins using the generated metabolic ontology.

**Supplementary Table 12.** Sodium transporters identified in the sc and sn RNAseq datasets and their function in kidney sodium reabsorption.

## Supplementary Information

Omics and imaging assays used within KPMP target different types of molecular components with different resolution, sensitivity and precision. An important function of the KPMP Central Hub is to integrate the different types of data using a set of analytical techniques. This process is summarized in Figure 1. Throughout the paper, we consistently use the same continuous color-code to identify different assays or cell types. The experimental assays that generate the raw data and all their technical details including standard operating procedures are detailed under ‘Data generation and initial analysis’ and publicly released with all their technical details and version-controlled release dates on the KPMP protocols.io page (https://www.protocols.io/groups/kpmp/publications).

### Identification of differentially expressed genes, proteins and metabolites

We analyzed data from four types of transcriptomic, two proteomic, one imaging-based and one metabolomic tissue interrogation assays. The pilot data presented for each assay comprises 3 to 48 different datasets that are obtained from 3 to 22 subjects (Suppl. Table 1). Kidney tissue was procured from a spectrum of tissue resources including from unaffected parts of tumor nephrectomy specimen (n=38), living donor preperfusion biopsies (n=3), diseased donor nephrectomies (n=5), and normal surveillance transplant (n=5) and native kidney biopsies (n=4). Single cell and nucleus transcriptomics clusters were obtained from previous analyses (*20, 21*). Within each assay we generated lists of differentially expressed genes (DEGs), proteins (DEPs) and metabolites that describe those genes, proteins or metabolites that are upregulated or enriched in a particular single cell cluster, single nucleus cluster or kidney subsegment, if compared to all other clusters or subsegments.

For pathway enrichment analysis and module identification, cluster-specific differentially expressed genes (DEGs) were obtained from published analyses from PREMIERE TIS (Michigan, Princeton, Broad) single-cell RNA sequencing (RNAseq) (*21*) and UCSD/WU TIS single-nucleus RNAseq (*20*) datasets. We excluded the clusters PT cells-3 and principal cells-2 from the single-nucleus RNAseq dataset, since these clusters showed an inflammatory or a stress response. Similarly, we excluded the cluster “Unk” from the single nucleus and the clusters “Pax8positivecells” and “LOH/DCT/IC” from the single cell RNAseq assays. Laser microdissected (LMD) RNAseq and proteomics (OSUIU), near-single-cell (NSC) proteomics (UCSF) and spatial metabolomics (UTHSA-PNNL-EMBL) datasets were individually processed as described in supplementary methods. Only DEGs and DEPs that indicate genes and proteins that are higher expressed in a particular cell subtype, type or segment were considered for all analyses.

### Ranking of differentially expressed genes and proteins

In the case of the DEGs and DEPs that were used for dynamic enrichment analysis, (*28*) module identification, (*32*) and *post hoc* power analysis, single nucleus and single cell DEGs were first ranked by adjusted p-value and then by decreasing fold changes (i.e., fold changes were used as a tiebreaker). Top ranked 300 entities with a maximum adjusted p-value of 0.05 were subjected to downstream analysis. Similarly, DEGs and DEPs obtained for each kidney subsegment based on LMD bulk RNAseq (*24*), or LMD and NSC proteomics, were ranked first by p-value and decreasing fold changes and the top ranked 300 DEGs and DEPs with maximum nominal p-value of 0.05 were subjected to pathway enrichment analysis or module detection (see below).

### Dynamic enrichment analysis

Top DEGs and DEPs for each podocyte cluster/glomerulus, PT cell cluster/tubulointerstitium and principal cell cluster/collecting duct subsegment were separately subjected to dynamic enrichment analysis using the Molecular Biology of the Cell Ontology (MBCO, version 2021) level-3 subcellular processes (SCPs) (*28*) that can be found at github.com/SBCNY/Molecular-Biology-of-the-Cell and www.mbc-ontology.org. The annotated interconnected hierarchy of MBCO is enriched using a unique algorithm that infers weighted relationships between functionally related SCPs. For all analyses we consider the top 25% weighted relationships. Dynamic enrichment analysis uses the top relationships to generate context-specific higher-level processes by merging functionally related SCPs that contain at least one DEG or DEP. The context specific higher-level SCPs contain all annotated genes of the original SCPs and are added to the annotated ontology to generate a context specific ontology. The context specific ontology at this point contains single and merged SCPs. This list is then used for enrichment analysis of the DEPs or DEGs using Fisher’s Exact test. All SCPs that are among the first seven predictions are connected based on the top inferred relationships, using solid lines. All networks for a particular cell type and the corresponding segment were merged and each SCP was color-coded according to the source assay(s) that initiated its dynamic enrichment. SCPs predicted by multiple assays contain multiple slices that are color coded accordingly. SCP size is determined by the number of assays that identified a particular SCP. Multiple subtypes or a particular cell type (e.g., PT cells) are all color coded by the same assay specific color. If an SCP was predicted for more than one subtype, it contains multiple slices colored with the assay specific color. SCPs predicted by different assays for the same cell type or corresponding segment were connected based on the top 25% inferred MBCO relationships, using solid lines. Additional well-known functionally related SCPs were connected using dashed lines.

We used the right-tailed Fisher’s Exact test to calculate the likelihood of obtaining the observed or a higher overlap between a list of DEGs/DEPs and a list of genes/proteins annotated to a particular SCP. To calculate this likelihood, we consider which genes or proteins have a chance to be identified as differentially expressed. Only genes/proteins that are detected by a particular assay and are statistically analyzed for differential expression can be identified as DEGs/DEPs. Consequently, only these genes/proteins are considered as the experimental background set for the Fisher’s Exact test. Similarly, the ontology background set only contains genes that have a chance to be assigned to a given SCP. In the case of the single cell (*21*) and nucleus (*20*) RNASeq datasets, all genes that are part of the UMI (Unique Molecular Identifier) read count matrices comprise the experimental background genes. In the case of the LMD bulk RNASeq, and the LMD and NSC proteomics datasets, the experimental background genes/proteins were all genes/proteins that were statistically analyzed for differential expression (Suppl. Tables 2, 3 and 4, respectively). MBCO contains an SCP that is labeled ‘Background genes’ and contains all genes that were identified during its population via text mining. The intersection of the experimental and ontological background genes/proteins is called background genes/proteins and is different for every assay and ontology combination. For additional statistical accuracy we removed all genes and proteins that were not part of the background genes/proteins from the lists of DEGs, DEPs and SCP genes before each enrichment analysis.

### Module detection

In parallel to enrichment analyses, we also performed another network-based pathway enrichment technique, identifying modules of cell-type specific marker genes within the kidney-specific functional network using the HumanBase interface (hb.flatironinstitute.org). For each cell type, module detection was performed using all cell-type-specific DEGs detected by single cell and single nucleus RNAseq (adjusted p-value <0.01) and segment-specific DEGs and DEPs detected by the other 4 technologies (nominal p-value < 0.01). Module detection is a network-based approach described in Krishnan et al., and construction of the functional networks is described in Greene *et al.* (*31, 32*). In contrast to the prior knowledge-based MBCO networks, the kidney-specific functional network is constructed using a data-driven regularized Bayesian framework based on the information in thousands of datasets, which include co-expression, transcription factor binding, protein-protein interactions, and other data types. Modules are detected using a community clustering algorithm based on connectivity between genes in the kidney-specific functional network, and enrichment analysis is subsequently performed to identify functional enrichments in each module.

### Enrichment analysis for metabolites

All glomerular and nonglomerular metabolites that were identified for the three subjects were merged and subjected to pathway enrichment analysis using MetaboAnalyst (*30*). Pathway analysis with the selections: Hypergeometric Test, Relative-betweeness Centrality, Homo Sapiens (KEGG), website version 3/22/2021. We analyzed which metabolites were part of the top six predicted metabolic pathways. We removed those pathways among the top 8 predictions that were predicted based on metabolites that are shared substrates in multiple pathways and consequently unspecific for the identified pathway (i.e. we ignored the glomerular pathways ‘Linoleic acid metabolism’, ‘alpha-Linoleic acid metabolism’, ’GPI-anchor synthesis’ and ‘Arachidonic acid metabolism’ that were predicted based on the lipids ‘Phosphatidylethanolamine’ and ‘Phosphatidylcholine’ and the pathway ‘Phenylalanine, tyrosine and tryptophan biosynthesis’ that was predicted based on the central precursor ‘3-(4-Hydroxyphenyl)pyruvate’). We mapped the kept MetaboAnalyst pathways onto MBCO pathways whenever possible; if those pathways did not have a corresponding MBCO pathway, the original pathway names were preserved. Since the non-glomerular metabolites contained multiple carnitine derivates we added the MBCO pathways “Carnitine shuttle” (based on L-Acetylcarnitine, Malonyl-Carnitine, L-Palmitoylcarnitine and L-Carnitine) and “Carnitine biosynthesis and transport” (based on L-Carnitine and 3-Dehydroxycarnitine) to the pathways predicted from spatial metabolomics, assigning the ranks 9 and 10, respectively.

### Integration of single-cell/single-nucleus transcriptomics

In contrast to bulk mRNA sequencing, where the gene expression measurements reflect an average across all captured cell types, single-cell or single-nucleus mRNA sequencing allows the measurement and comparison of comprehensive gene sets obtained from individual cells. This approach enables mapping of cellular heterogeneity with high throughput. In the first phase of the project, three KPMP Tissue Interrogation Sites (TISes) performed this approach to generate single cell/single nucleus expression data from normal adult kidney tissue. In addition to locally acquired kidney tissue samples, each TIS also used a set of common KPMP pilot tumor nephrectomy tissue samples to generate the expression data. Single-cell transcriptomic data was produced by PREMIERE (24 libraries from 22 subjects) (*21*) and UCSF (10 libraries from 10 subjects), whereas the single-nucleus data was made by UCSD (47 libraries from 15 subjects) (*20*). Following is a brief description of the integration of the data from the three sites.

Data from each site were first processed using the Seurat 3.0 R package (*118*). As a quality control step, nuclei/cells with less than 500 and more than 5,000 features and more than 20% mitochondrial genes were removed. The processing steps included normalization and identification of highly variable genes. We then removed potential doublets using DoubletFinder (*119*) from each dataset. Next, we used the integration algorithm embedded in the Seurat R package to perform combined analysis of single-cell/single-nucleus transcriptomic data. The integration algorithm first identified a set of anchor genes in each processed dataset. These anchor genes were then used to harmonize the datasets. The downstream process included scaling, principal component analysis, batch integration using harmony, dimensionality reduction using Uniform Manifold Approximation and Projection (UMAP), and unsupervised clustering. The clustering was performed at a low resolutions (clustering granularity of 0.5). Enriched genes for each cluster compared to all other clusters were identified using the Wilcoxon rank sum test.

### Integration of single-cell, single-nucleus and laser capture microdissection bulk transcriptomics

To integrate single-cell sequencing, single-nucleus sequencing, and LMD bulk transcriptomic datasets, we first determined the overlap between genes identified both in the LMD dataset and in the corresponding single-cell transcriptomic dataset. From this set of shared genes, we restricted further analyses to a subset of genes showing variable expression in the single-cell dataset. We then computed the Pearson correlation between each individual cell in a scaled single cell/single nucleus dataset and the LMD transcriptomic dataset. For this correlation, we used the logarithmized mean fold change that was obtained by dividing the average expression of each gene within a subsegment by the average expression of the same gene within all other subsegments. Using this approach, we can assign each cell to the appropriate LMD segment that shows the highest correlation value. To evaluate the overall segment assignments for individual cell clusters, we examine the normalized distribution of cells assigned to each LMD segment within a given single-cell cluster and present this as a normalized heatmap that represents overlap between different transcriptomic assays.

### Proteomic-transcriptomic co-expression analysis

LMD and NSC proteomic datasets identified protein expression in two kidney subsegments: glomeruli and tubulointerstitium for LMD and glomeruli and proximal tubule (PT) for NSC. Here, we did not combine sequencing and proteomic results of multiple subjects to generate DEGs and DEPs, but compared the results obtained for each individual person. Since only one dataset per segment was generated from each individual person by the LMD and NSC technologies, we could not calculate p-values in this analysis. Furthermore, both proteomic technologies only generated results for 2 subsegments, i.e. the glomerular and PT segments for NSC proteomics and the glomerular and tubulointerstitial subsegments for the LMD proteomics. Consequently, we collectively calculated the fold changes between podocyte/glomeruli and PT/tubulointerstitial cells or subsegments for each individual subject.

For the single cell and nucleus transcriptomic datasets, we identified technology and subject specific cluster gene expression, using the “Average Expression” functionality embedded in the Seurat R package (RNA assay, counts slot) on the cells/nuclei assigned to the same clusters in the integrated single cell and nucleus RNAseq data analysis described above. The gene lists of all PT clusters of an individual subject and technology were merged (Suppl. Figure 2). If a gene was identified by more than one cluster, we defined the highest expression value as the merged expression value for that gene. For each technology we characterized all genes/proteins that were identified in at least one cluster or subsegment of at least one subject and defined these genes/proteins as a technology specific background set. The intersection of all background sets was defined as the set of common genes. Subject-specific podocyte or glomerular gene and protein expression was calculated by dividing gene and protein expression in podocytes, or glomeruli, by gene and protein expression in PT cells or PT/tubulointerstitial subsegments, after adding 1 to prevent division by 0. Ratios were inverted to describe PT/tubulointerstitial specific gene expression. Log_10_ absolute expression values and log_2_(ratios) of all genes/proteins or all common genes/proteins were subjected to pairwise correlation, followed by hierarchical clustering. Log_2_ ratios were averaged over each subject within each technology and pairwise Pearson correlation coefficients were determined between the different technologies using the set of common genes. Mean log_2_ ratios were averaged across the four RNAseq platforms and the two proteomic platforms, followed by determination of the Pearson correlation coefficient using the set of common genes.

### Comparison of cell type-specific imaging and transcriptomic expression data

To integrate cell type-specific imaging and transcriptomic data, we first constructed matrices with average expression values for each gene in each cell type cluster for both the set of 16 normalized integrated transcriptomic clusters and the CODEX clusters. We normalized each gene in both transcriptomic and CODEX matrices to have a mean of 0 and standard deviation of 1. We then filtered both datasets to include only genes represented in both the transcriptomic and the imaging datasets and computed the average expression of each gene/protein in each cell type. We next considered the problem of constructing a matrix to computationally map transcriptomic cell clusters to the imaging cell clusters. Specifically, let A be the N x k1 matrix of average protein expression values by imaging data clusters, C be the N x k2 matrix of average gene expression values by transcriptomic clusters, and M be the k1 X k2 matrix that maps A to C. We want to find M such that AM ≈ C. We can approximate M by taking the Moore-Penrose pseudoinverse of A, denoted A+, with M ≈ (A+)(C). M then provides a set of weights that map the imaging cell types to the transcriptomic cell types, with a large value for an entry in M in position (i, j) indicating that the imaging cell type i makes a large contribution to approximating the expression vector of transcriptomic cell type j as a linear combination of imaging cell types. Before visualizing matrix M as a heatmap, we first normalized each row to have mean of 0 and standard deviation of 1 in order to identify the transcriptomic cell types that are weighted most heavily in the mapping to each imaging cell type.

### *Post hoc* power analysis

The single-cell RNAseq and single-nucleus RNAseq datasets were obtained from 22 and 15 subjects, respectively, whose samples were sequenced in 24 and 47 libraries. We used these datasets to assess the reproducibility and reliability of both assays in a *post hoc* power analysis. This analysis compares results by the full datasets with the results by down-sampled datasets where libraries are randomly and systematically removed from the full data.

Both full datasets were separately subjected to a standardized Seurat pipeline for the identification of single-cell (or -nucleus) clusters and DEGs. Nuclei and cells with less than 500 and more than 5,000 features as well as more than 50% mitochondrial genes were removed. ‘SCTransform’ was used for data normalization and scaling (based on top 2,000 features), followed by principal component analysis. The first 30 principal components were used for dimensionality reduction before identifying single nucleus/cell clusters (resolution = 0.8).

Top 300 DEGs of each cluster were identified (adjusted p-value: 0.05) and compared with literature-curated cell-type specific essential genes (Suppl. Table 10) using Fisher’s Exact test to assign a kidney cell type to each cluster. The assigned cell type is that cell type whose essential genes had the most significant enrichment among the DEGs of that cluster. To document the reliability of that cell type assignment we compared its p-value to the p-value of the second prediction (that cell type whose essential genes had the second most significant enrichment among the DEGs of that cluster). The larger the distance between both p-values, the more reliable the cell type assignment. The number of clusters that were assigned to each cell type was documented. Nuclei and cells that were assigned to a particular cell type and map or do not map to the corresponding LMD tissue subsegment were counted as well, based on the subsegmental correlation analysis as described above. The top 300 DEGs were subjected to dynamic enrichment analysis using MBCO. All SCPs among the top seven predictions were further investigated.

We progressively and randomly removed libraries from the full (reference) datasets to generate 100 non-overlapping downsampled datasets for each number of remaining subjects. Downsampled data was subjected to the same analysis pipeline and results were compared with the reference results. We calculated the percentage of downsampled datasets for each number of remaining libraries that identified a particular cell type. If a particular cell type was identified in a down-sampled dataset we counted how many of its nuclei/cells were assigned to the same or to a different cell type in the reference analysis. To visualize both counting results in the same plot, we defined those cell counts that mapped to a different cell type to be negative, so these counts are plotted below the abscissa. Similarly, we counted how many nuclei/cells of a particular cell type mapped and did not map to a particular tissue subsegment that is indicated in the title of the plot. Here, we also defined those cell counts that mapped to a different subsegment to be negative. We calculated the Pearson correlation between the DEGs of each cell type in the downsampled datasets and the reference datasets based on log_2_(fold changes).

Pathway enrichment analysis normally involves identification of the most significant pathways irrespective of their p-values. To document the reliability of the identified SCPs we identified the ranks of the SCPs that were among the top seven predictions in each downsampled dataset. Ranks were averaged for each SCP and number of analyzed libraries.

### Documentation of cellular metabolism

We generated a small ontology that contains the major metabolic pathways involved in energy generation and sphingomyelin synthesis. We defined parent-child relationships, where child pathways described sub-functions of their parent pathways (Suppl. Figure 17A). Pathways were populated with genes curated from the literature, parent pathways also populated with the genes of their child pathways. The ontology is publicly available at github.com/SBCNY/Molecular-Biology-of-the-Cell and mbc-ontology.org.

We subjected the top 500 significant marker genes and proteins (sc/sn RNAseq: adjusted p-value 0.05, LMD RNAseq, LMD/NSC proteomics: nominal p-value 0.05) to enrichment analysis using this ontology and Fisher’s Exact Test. Investigation of the predicted pathways that are specific for a particular reaction allowed to decide in which reaction(s) those enzymes participate that are shared by multiple pathways. Child pathways that specifically describe the function of their parent pathways are visualized in the same color in supplementary figure 17A. If only pathways that contained the shared reactions of multiple parent pathways were predicted, we assumed that they participated in the default parent pathways “Glycolysis”, “Keton body catabolism” or “Aerobic glycolysis”.

Since the sc RNAseq data was derived from cortical, medullary and mixed samples (Suppl. Figure 4B), we distinguished between medullary (DTL, ATL1-3, TAL-1, PC-3 and IC-A2) and cortical cell types (all other cell types of the renal tubule of the nephron). All other datasets were assigned as cortical. Enrichment result negative log_10_(p-values) were first averaged across the different cell subtypes of the same cell type and then across the different transcriptomic datasets. In case of the sc and sn RNAseq assays, we considered the amount of cells assigned to each subtype of a particular cell type. The averaged negative log_10_(p-values) is representative of the cell counts of each cluster.

### Comparison of experimental reabsorption and gene expression profiles

Experimentally determined reabsorption capacity profiles that describe what percentage of a filtered sodium or glucose is reabsorbed in a particular nephron segment were curated from standard medical and physiological text books (*92–94, 120*), followed by averaging of the curated numbers for each ion or molecule. As these are widely used medical school textbooks, we assumed that the information is correct and did not further track down the values given in these books to the primary papers from which these values were obtained. Also we assumed conversation of physiological processes across mammalian species and we did not ascertain if all values were derived from the same species or more than one species.

#### Generation of a transmembrane transport ontology

Since human single cell and single nucleus RNAseq datasets (*20, 21, 97, 98*) contain gene expression profiles in all major nephron cell types, we reasoned we could compare segment specific gene expression levels of the transporter or channel of interest with these physiologically measured reabsorption profiles. Using Gene Ontology ^7^, Molecular Biology of the Cell Ontology (*28*), Wikipedia articles and selected reviews as sources (*66, 99, 121-124*), we generated a comprehensive ontology of transmembrane transporter at the plasma membrane (Supplemental Figure 19A). Within GO we focused on all Biological Processes and Molecular Functions that were children of “sodium ion transport”, “sodium ion transmembrane transporter activity” as well as “glucose transport”, “glucose transmembrane transporter activity”, as defined by the “is_a” and “part_of” relationships. From MBCO we added all genes assigned to the subcellular processes (SCPs) “Sodium transmembrane transport”. The initial list of transporter candidates was manually investigated to validate their transporter activity. True positives were assigned to de novo subcellular processes (SCPs) that describe the movement mechanism (i.e., transport via symporter or antiporter), the movement direction (i.e. lumen-to-blood or blood-to-lumen) and all ions or molecules that are transported by that mechanism. In case of antiporters, we specified which ions or molecules move in opposing directions by separating them with the term ‘vs’. If the protein translated from a particular gene had a unique name that is commonly used and is different from the official NCBI gene symbol, we assigned the gene to that particular protein name (e.g., SLC12A1 and SCL12A3 were assigned to NKCC2 and NCC, respectively). Here, we did not describe the activity mediated by that protein (e.g., “Sodium potassium chloride transport by the symporter NKCC2”), since this would create unnecessarily long names in our figure legends. Nevertheless, in all analyses these proteins were processed as if they were SCPs. Consequently, whenever we use the term SCP in the manuscript, we refer to these proteins as well. Each SCP-gene association was supported by at least one reference that could be the NCBI gene summary, UniProt gene summary or a PubMed ID for a supporting article (Supplementary Table 12). To allow systematic analysis and grouping of transmembrane movements, we integrated all SCPs into a SCP hierarchy of parent and children SCPs (Supplementary Figure 19B) using a strategy we have described for the MBCO ontology (*28*). This hierarchy converges children on parent SCPs that describe more generalized shared transport mechanisms. For example, the SCP “Sodium potassium chloride symporter” (that is the parent of the SCP “NKCC”) is the child of the two parent SCPs “Sodium chloride symporter” and “Sodium potassium symporter”. We left out the SCP “Potassium chloride symporter”, since here we focused on sodium and glucose transmembrane transport. These SCPs are then connected to the higher-level SCP “Sodium lumen-to-blood transport by symporter”. For both sodium and glucose all SCPs finally converge on either one of two different overall parent SCPs describing transcellular lumen-towards-blood transport and transcellular blood-towards-lumen transport. Discussed example of SCP relationships and two additional examples are shown in supplemental figure 18B. Supplemental figure 18C shows the hierarchical organization of all SCPs involved in sodium lumen to blood and blood to lumen transport. All parent SCPs were populated with the genes of all of their children SCPs. Finally, we kept only those genes in the ontology that localize to the plasma membrane based on the jensenlab human compartment ontology with a minimum confidence score of 4 (out of maximal 5) (www.compartments.jensenlab.org)

#### Calculation of predicted reabsorption capacities

Besides our own single cell (sc) and nucleus (sn) RNAseq dataset (*20, 21*), we utilized two different snRNA seq datasets that were generated from undiseased tissue as well (*97, 98*). All datasets document how many mRNA molecules are transcribed from each gene in each individual cell. These numbers are described as UMI (Unique Molecular Identifier) counts (*125*), but in this study we use the term mRNA counts or levels to indicate that it is a quantitative measure of mRNA levels of a certain species. The cells and nuclei in the sc and sn RNAseq data sets were previously grouped into clusters using standard software packages (Seurat XX), followed by identification of cluster specific marker genes and cell type and subtype annotation (*20, 21, 97*). We analyzed the raw UMI matrix (GSE114156) (*98*) using the seurat package (as outlined in Suppl. Figure 5A) and annotated kidney cell types based on cell type specific gene expression (Suppl. Table 10).

We assumed that that mRNA molecule counts (i.e., UMI counts) of each transporter or channel in each cell reflect the capacity of that particular cell for transmembrane movement of that particular ion or molecule. The following explanation of how we predicted movement capacities from those mRNA levels is summarized in supplemental figure 18A. We initially processed all four datasets, i.e., one sc RNAseq and three sn RNAseq datasets, independently. For each dataset and SCP of our transmembrane movement ontology we summed up all mRNA molecules that are expressed in all cells of a particular cell type or nephron segment and map to genes involved in that SCP. It should be noted that we documented total and not mean capacities, because we did not divide the mRNA count sums by the number of cells in each particular cell type or segment. If a particular cell type or nephron segment contains more cells, it is assumed to contribute more to the reabsorption of a particular ion or molecule, if the appropriate transporter is present. Measured physiological reabsorption profiles describe net lumen-to-blood transport values in each segment. To account for the different transport directions predicted from SCPs that are involved in lumen-to-blood and blood-to-lumen transport, we defined all mRNA levels mapping to blood-to-lumen transport SCPs as negative. This allowed the calculation of net lumen-to-blood transport capacities by adding up all mRNA counts involved in lumen-to-blood transport and all (negative) mRNA counts involved in blood-to-lumen transport of each ion or molecule. Since the physiological profiles document how much percent of a particular ion or molecule is reabsorbed in a particular nephron segment, we expressed all SCP capacities in percent of the net lumen-to-blood transport capacities of the corresponding ions or molecules. Consequently, the sum of all predicted transport capacities along the nephron is 100% for both sodium and glucose (mRNA levels assigned to blood-to-lumen transport are still defined as negative), allowing the direct comparison of mRNA levels and reabsorption profiles. Any SCPs that mediate the transport of multiple ions or molecules, were normalized independently for each ion and molecule to calculate the relative contribution of that SCP to the total reabsorption capacity of each ion or molecule. Final percentages of the same SCPs predicted by the three single nucleus RNAseq datasets or all 4 datasets were averaged.

### Identification of cell-type markers

Marker genes and proteins of each cell subtype or segment were ranked as described above. If multiple cell subtypes were identified based on the single cell or single nucleus RNAseq datasets, we calculated the average rank for that cell type. Original or averaged ranks were averaged again over the different assays for each cell type and selected segment, followed by re-ranking. Top five re-ranked genes/proteins were selected as cell type markers in Table 1.

### Generation of nephron schema

We used BioRender.com to create the nephron schema in Figure 7.

### Data generation and initial analysis

Seven different RNAseq, proteomics, metabolomics and imaging datasets were generated and analyzed by five different TISes. The PREMIERE TIS (composed of Michigan, Princeton, Broad) generated single cell RNASeq data, the USCD/WashU TIS generated single-nucleus data, the UCSF TIS generated single-cell RNASeq, near-single-cell proteomics and Codex imaging data, the IU/OSU TIS generated laser microcapture dissection (LMD) RNASeq and LMD proteomics data and the UTHSA-PNNL-EMBL TIS generated spatial metabolomics data.

### Single-nucleus RNAseq (UCSD/WashU) and Single-cell RNASeq (PREMIERE)

UMI count matrixes and list of differentially expressed genes were downloaded from published analyses for the PREMIERE TIS (composed of Michigan, Princeton, Broad) single-cell RNA sequencing (RNAseq) (*21*) and UCSD/WashU TIS Single-nucleus RNAseq (*20*) datasets. We excluded the PT cells-3 and principal cells-2 clusters from the single-nucleus RNAseq dataset, since these clusters showed an inflammatory or a stress response.

### Subsegmental LMD Transcriptomics (IU/OSU)

A comprehensive Laser MicroDissection (LMD) protocol is published on protocols.io (https://www.protocols.io/view/laser-microdissection-8rkhv4w). Briefly, 12 μm frozen sections are obtained from an Optimal Cutting Temperature (OCT) preserved tissue block and adhered to LMD membrane slides (Leica, Buffalo Grove, IL). Tissue undergoes a rapid staining protocol involving acetone fixation, washes with RNAse-free PBS, and antibody incubation in 10% bovine serum albumin. Slides undergo dissection with a Leica LMD6500 system with pulsed UV laser. After collecting a minimum tissue area of 500,000 μm^2^ in an RNAse-free micro-centrifuge tube, RNA is isolated using the PicoPure RNA IsolationKit according to manufacturer’s instructions (Applied Biosystems, Cat# KIT0204). RNA quality is assessed by bioanalyzer, ribosomal RNA is depleted, and cDNA libraries are prepared using the SMARTer Universal Low Input RNA Kit (Takara, No. 634938). Sequencing was conducted on an Illumina HiSeq4000. Mapping was performed using STAR (v2.5.2b) and read counts were quantified with featureCounts (subread v.1.5.0). Total read counts mapping to each gene were generated with edgeR, normalized, and converted to expression ratios.

Segment specific gene expression was compared to the gene expression in all other subsegments using an unpaired ttest with equal variance. Subsegment specific gene expression ratios were calculated similarly.

### Subsegmental LMD Proteomics (IU/OSU)

A comprehensive Laser MicroDissection (LMD) proteomics protocol is published on protocols.io https://www.protocols.io/view/laser-microdissection-for-regional-transcriptomics-8rkhv4w?version_warning=no. Our LMD proteomic methods have also been previously published in detail (*126, 127*). Briefly, 10 μm frozen sections are obtained from an OCT preserved tissue block and adhered to polyethylene naphthalate (*PEN*) membrane *slides* for LMD. Frozen sections are fixed in 70% ethanol, incubated in H_2_O to remove OCT, briefly stained with hematoxylin, and dehydrated in ethanol. LMD is performed and glomeruli and tubuloninterstitial samples are collected separately in 0.5% Rapigest/50 mM NH_3_HCO_3_ solution. The collected samples are then boiled for 20 minutes for protein retrieval and digested overnight with trypsin. Peptides are dried, re-suspended in acetonitrile/formic acid and analyzed using liquid chromatography tandem-mass spectrometry (LC-MS/MS) analysis using an Easy-nLC 1000 HPLC coupled to an Orbitrap Fusion mass spectrometer (Thermo Scientific, Waltham, MA). Data is searched using Proteome Discoverer 2.1 (Thermo Scientific) and searched against a human Uniprot database (version 05/26/18). Data are analyzed following global normalization of spectral counts.

Glomerular gene expression was compared to the tubulointerstitial gene expression using an unpaired t-test with equal variance. Glomerular to tubular specific gene expression ratios were calculated similarly.

### 3-D Immunofluorescence Imaging and Tissue Cytometry (IU/OSU)

The entire 3-D fluorescence imaging and tissue cytometry protocol is published on protocols.io (<u>dx.doi.org/10.17504/protocols.io.9avh2e6</u>). Briefly, frozen cores are sectioned at 50 μm using a cryostat and fixed using 4% paraformaldehyde. A panel of up to 8 antibodies was incubated to identify renal and immune cell types. Images were acquired in up to 8 channels using a Leica SP8 Confocal Microscope. Volume stacks spanning the whole thickness of the tissue were taken using a 20× NA 0.75 or 40× NA 1.3 objectives with 0.5- to 1.0-*μ*m spacing. Large scale confocal imaging of overlapping volumes was performed with an automated stage and stitched using Leica LASX software (Germany). 3-D image rendering was done using Voxx v2.09d. The 3-D tissue cytometry was performed on image volumes using VTEA, which was developed as a plugin for ImageJ/FIJI as previously described (*128*).

### CODEX Imaging (UCSF)

The CODEX system is the combination of an (1) oligo-nucleotide based antibody labeling-detection technique, (2) a microfluidics instrument coupled with an inverted microscope capable of whole slide scanning, and a (3) software suite that consists of an image processor and an ImageJ-based image analysis solution (*129*). First, a section from an optimal cutting temperature compound-embedded tissue block is cut and incubated manually in a single step, with a set of antibodies each tagged with a unique oligonucleotide sequence. The following phase consists of iterative cycles of detection, imaging, and dye removal. In each cycle, a maximum of three targets are revealed by spectrally distinct dyes (AF488, Atto 550, and Cy5) tagged with oligonucleotides complementary to the oligonucleotide tag of a given antibody.

The acquired images are processed by the CODEX processor in a set sequence of steps: shading correction, tile registration, deconvolution, drift compensation, overlap cropping, background subtraction, best focus detection/interpolation, stitching, cell segmentation, and spillover compensation.

The output of the cell segmentation step of image processing is an .fcs file (similarly to flow-cytometry solutions). This file contains the individual fluorescent intensity values (can range from 0 to 65k) of each cell for each marker. Fluorescent intensity values allow the definition of cell populations by manual gating of the segmented cells using visual assessment of the image and previous literature data on the expression pattern of our marker set in human kidney.

Native renal biopsies taken at University of California, San Francisco from patients with minimal change disease (n = 3), thin-basement membrane disease (n = 1), and post-surgical biopsies from tumor nephrectomies (n = 2) were used. In addition, case 18-162 from KPMP pilot sample pool was also processed (Suppl. Table 1).

### Spatial Metabolomics (UTHSA-PNNL-EMBL)

10 μm thick renal cortical tissues were sectioned using a cryostat (Leica Microsystems), thaw mounted on indium tin oxide coated slides (Bruker Daltonics), and prepared for matrix-assisted laser deposition/ionization mass spectrometry (MALDI-MSI) by spraying with 2,5- dihydroxybenzoic acid (DHB; 40 mg/mL in 50% MeOH:H_2_O) using the TM-Sprayer automated spraying robot (HTX Technology). The following spraying parameters were used: 80 °C nozzle temperature, a flow rate of 0.05 mL/min, 10 passes, a N_2_ pressure of 10 psi, a track spacing of 3 mm, and a 40 mm distance between the nozzle and sample was maintained. MALDI-MSI was performed using a MALDI-FTICR imaging mass spectrometer (Bruker Daltonics) set at a 120,000 resolving power at m/z 400 or a MALDI-Orbitrap mass spectrometer (Thermo Scientific) set at the 120,000 resolving power at m/z 200. The data was inspected following the quality control guidelines as developed within KPMP and converted into the imzML centroided format using the SCiLS software (Bruker Daltonics) or ImageInsight software (Spectroglyph, LLC), followed by the submission to METASPACE and annotation against the SwissLipids and HMDB molecular databases with the false discovery rate of 20%, as described in Reference ^33^.

We have developed an approach to find glomeruli markers in MALDI-MSI data by using METASPACE and co-localization analysis. First, we have selected a template marker that was localized within the glomerular regions, as confirmed by the histology. This ion was annotated by METASPACE as ceramide phosphate CerP(d34:1) ^33^. Then, we performed a spatial co-localization analysis by calculating for all other detected metabolites and lipids their spatial correlation with CerP(d34:1) using the cosine score. The molecules with the correlation above 0.2 were considered and manually curated to show the co-localization with the glomeruli regions by overlaying every ion image with the histological image. The resulting 30 markers were uploaded to the KPMP DataLake and were used for the multiomics integration analysis.

